# CHARON: Estimating the drift time and the number of individuals in environmental DNA with diploid individuals

**DOI:** 10.1101/2025.05.26.656139

**Authors:** Jan van Waaij, Mads Vodder Hartmann, Peter Wad Sackett, Gabriel Renaud

## Abstract

Environmental DNA (eDNA) offers a promising avenue for reconstructing the genetic diversity and demographic history of ancient populations. However, the analysis of eDNA from humans or forensic DNA, presents significant challenges, including low coverage, DNA degradation, and the uncertainty of the number of individuals contributing to a sample. This study introduces CHARON, a novel statistical method for jointly estimating the number of individuals and drift times between a human eDNA or forensic sample and a given population with known allele frequencies. We validate our method through simulations and synthetic empirical data and show that we can reliably estimate the number of individuals up to 8 at a coverage between 2X and 4X. Our method can also pinpoint the most likely population of origin for eDNA or forensic samples. This work provides a tool for the application of human eDNA in evolutionary and forensic studies and an implementation is available here: https://github.com/Jan-van-Waaij/Charon

## 1 Introduction

Environmental DNA (eDNA) has rapidly evolved into a transformative tool for ecological monitoring, enabling the detection of species in various environments without the need for physical capture or direct observation [2]. By collecting and analyzing DNA fragments shed by organisms into their surroundings—whether it be in water, soil, or air—researchers can identify the species present in an area, offering a non-invasive means of studying biodiversity [8, 11]. For example, airborne eDNA has been effectively utilized to track seasonal changes in plant communities and to assess the impact of human activities on biodiversity at a landscape scale [16, 1]. Moreover, the ability to capture eDNA from various environments, including air and water, has opened new avenues for environmental monitoring that can respond rapidly to ecological shifts, such as those induced by climate change or human encroachment [12, 20].

The applications of eDNA extend beyond ecological studies into the realm of forensic science, where it is being increasingly recognized as a valuable method for the non-invasive collection of human DNA. In forensic contexts, eDNA has demonstrated significant potential in scenarios where traditional DNA sampling methods may be challenging, such as in aquatic environments or across large areas [5, 12, 18, 30, 32]. Some studies have indeed shown that eDNA can be used to recover human DNA from water, offering new avenues for locating human remains or tracing human presence in forensic investigations [9, 10]. Additionally, the use of airborne eDNA in forensic investigations could provide critical evidence in crime scene analyses, especially in situations where physical evidence is scarce [13, 24]. As these methods continue to be refined, they hold promise for enhancing the capabilities of forensic science, providing innovative and less invasive approaches to evidence collection and analysis [4, 31].

An issue facing both the forensics DNA community as well as the ancient DNA (aDNA) community is the presence of multiple individuals in the mix [21, 25]. The present-day human individuals involved in either excavation or library preparation can mix their DNA with the DNA from the ancient bones and bespoke computational methods had to be developed to quantify as contamination [23, 28]. Racimo and Renaud *et al*. developed a method to jointly estimate present-day contamination as well as demographic parameters [26]. This model revolved around computing the posterior probability of the genotype and divergent bases were either due to i) contamination error ii) heterozygous sites or iii) sequencing errors. The demographic parameters that were inferred were drift times which provided the prior on the genotype. Later, Schraiber *et al*. generalize this model to multiple samples [29].

Inferring drift times between populations is crucial for understanding population structure and history, as it allows us to reconstruct much of this history from the genomes of contemporary individuals. For human eDNA or forensic DNA, drift times to other populations would be informative to provide the ethnicity of the individuals in the mix. Another key piece of information for forensics DNA and human eDNA is the number of individuals that contributed to the blend.

To address the challenges of analyzing eDNA and forensic DNA with multiple contributing individuals, we introduce CHARON, a novel statistical method that jointly estimates both the number of individuals and the drift times between an ancient or forensic sample and a modern reference population (see Fig. 1). Our approach leverages allele frequencies from known populations to infer these parameters, providing a more accurate understanding of the genetic makeup of the samples.

**Figure 1.**
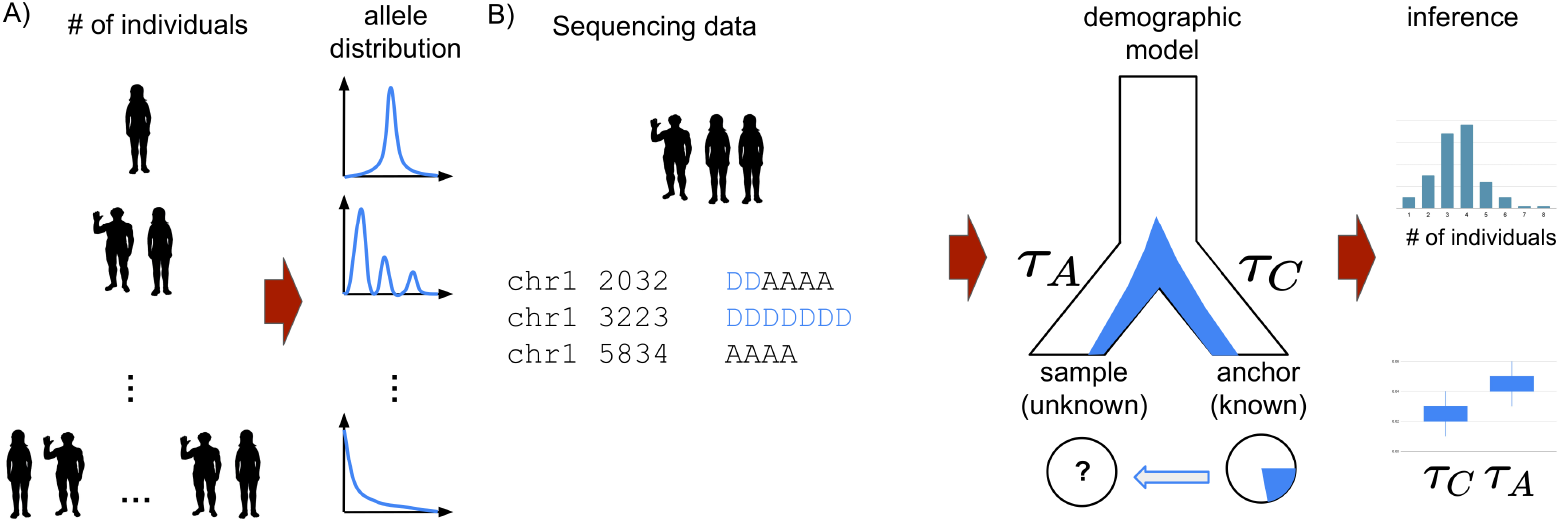
The general workflow for estimating the number of individuals. A) For a single individual with 2 chromosomes, the distribution of the bases around polymorphic sites will follow a binomial distribution around 0.5. For two individuals thus carrying 4 chromosomes, the distribution will be skewed around 0.25, 0.5 and 0.75 when 1 allele is present, 2 and 3, respectively. As the number of individuals increases, this distribution takes the shape of the site frequency spectrum. B) Our method relies on taking base counts from samples of numerous individuals for which we know the allele frequency in an anchor population i.e. a population that is relatively close with well-resolved allele frequencies. Drift parameters are estimated along with an estimate of the number of individuals.

We make the following two assumptions about the data i) they are all from the same population in the genetics sense ii) they have all had more or less the same contribution to the sampled data. By employing simulations and synthetic empirical data, we demonstrate that our method is robust across varying levels of coverage, reliably estimating up to eight individuals with moderate coverage and accurately pinpointing the population of origin. We provide an implementation of our methods written in the Julia programming language here https://github.com/Jan-van-Waaij/Charon.

## 2 Materials and Methods

### 2.1 Test data generation

#### 2.1.1 Simulated data

We generated two types of simulated data. i) a very simple demographics scenario with 2 populations and known drift parameters to determine if our approach could accurately estimate them ii) a more complex demographic model based on human history to see if our method could pinpoint the population of origin. To simulate genetic data, we used msprime [17], a coalescent-based simulation tool that can generate genetic data for large sample sizes.

For the simple model, we used msprime to simulate the population history in Fig. 3, with *τ*_*A*_ = *τ*_*C*_ = 0.25, and the effective population size is 2000 for both populations. See the code in Section S6.1 in the supplementary material. We use the anchor population to calculate the frequencies of the alleles, and we randomly pick *n* = 1, 2, .., 8 individuals from the ancient population and use them to simulate reads with constant coverage 100. We used downsampling to get lower coverages. We have set the error rate equal to 3 ‰. We have a sequence of length 10^7^ and 155’776 SNPs, which have frequencies strictly between 0.0 and 1.0 in the anchor population. The results are found in Section 3.1.1

**Figure 2.**
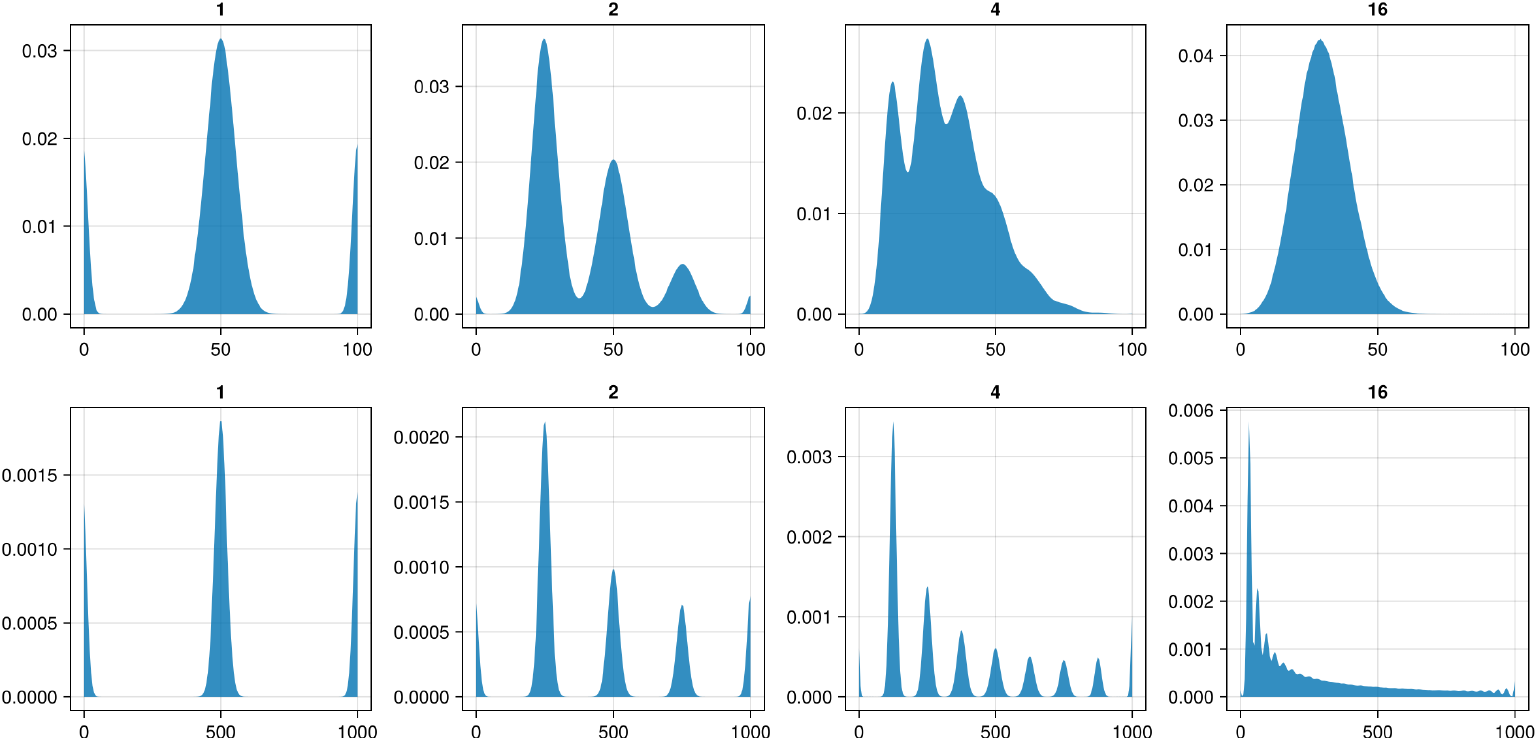
Top) Histograms simulating the read distributions for an allele frequency of 0.3 and 10M replicates. The read distribution from binom(100, *k/*(2*n*)), where *k* in turn is simulated from binom(2*n*, 0.3), for 1, 2, 4 and 16 individuals. Bottom) Same as above but for all sites in the genomes where the derived read frequencies are from a site frequency spectrum. For a single individual, the distribution is a binomial centered around 0.5 however as the number of individuals increases, this distribution converges to the site frequency spectrum.

**Figure 3.**
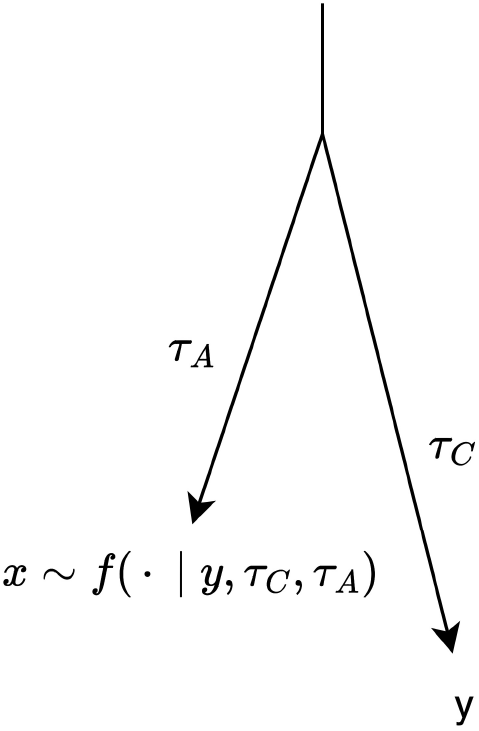
*τ*_*C*_ and *τ*_*A*_, model parameters that represent the drift times of the anchor population and sample population, respectively. We seek *x*, the derived allele frequency in the sample population, with distribution *f* which itself depends on *τ*_*C*_ and *τ*_*A*_ and *y*.

We then sought to simulate a more realistic demographic scenario with multiple populations. For this simulation, input demographic data was provided in the Demes format [14]. Demes offers a way to describe demographic models involving multiple interacting populations, migration events, and growth rates via a YAML-based format. To specify the demographic model, we used a pre-defined model provided by the Demes specification, accessible via their official website [7]. In particular, the starting point for the model we employed is the Gutenkunst out-of-Africa (OOA) model that describes the demographic history of African, European, and Asian populations [15]. This model was then adapted to include the demographic history Indigenous peoples of the Americas [19].

In our model, we implemented population splits occurring 75 years (3 generations) ago. At this time, each of the four populations were duplicated, creating an identical copy of itself, and all of these new populations were isolated from their original counterparts. These new, isolated populations were referred to as “splits”. Each split maintained the same effective population size as the original population at the moment of the split. With these models finalized, simulations were performed, leading to the generation of VCF files. Details about the demographic model as well as the results are found in the Results in Section 3.1.2. The source code can be found in Section S6.2 of the supplementary material. The results are found in Section 3.1.2.

#### 2.1.2 Synthetic empirical data

As we cannot get empirical data with a known number of contributing sources, we can however combine whole genome sequencing files together to create a synthetic empirical data set of known origin and number of contributing individuals. We mixed several individuals with data from the 1000 Genomes Project [6], to create a realistic environmental DNA sample. Briefly, we took BAM files from 8 individuals pertaining to the the Iberian population (IBS). We downsampled to create files of relatively equal coverage and merged the files in equal proportions to create a realistic synthetic empirical BAM file. We simulated 1, 2, 4 and 8 individuals. The final coverage for all BAM files ranged from 10X for the BAM file with 1 individuals, 7X for 2, 12X for 4 and 24X for the BAM with 8 individuals (estimated using mosdepth [22]), which were further downsampled to test the robustness of the method. The results are found in Section 3.2.

### 2.2 General idea of our method

In this paper, we rely on the model described by Schraiber [29], who calculates probabilities of having a total of *k* derived alleles at a certain locus among *n* diploid individuals and with given drift times. We use this by putting an additional prior on *n* and the drift parameters, to estimate *n* and the drift parameters simultaneously, and provide credible intervals as well.

To provide an insight as to how we can estimate the number of individuals, *n*, we will show that the distributions for different *n* differ from each other. For a certain locus, assume the derived allele in the population the sample pertains to has frequency *x* ∈ [0, 1]. Under the Hardy-Weinberg equilibrium, the total number of derived alleles, *k*, among the *n* diploid individuals is binomially distributed with parameters 2*n* and *x*. Given *k* ∈{0, …, 2*n*} derived alleles, the number of derived reads is binomially distributed with parameters *R* and *k/*(2*n*), where *R* is the coverage. Thus, the derived read count follows a mixture of 2*n* + 1 binomials. Note that we do not require knowing which read comes from which individual.

For large *n*, the distribution of the reads converges more and more to a binomial distribution with parameters *R* and *x* (Lemma S4 and Section S1 in the supplementary material). As *n* is small, the different modes are easily distinguishable, which makes it easy to estimate *n*. For larger *n*, it is harder. To illustrate this principle, we simulate *k* derived alleles with frequency 0.3, and given *k* derived alleles, we simulate derived reads with probability *k/*(2*n*) and coverage 100. The densities for the different *n*, based on 10M simulation are shown in the top row of Fig. 2. We do the same where the frequency is distributed with the site frequency spectrum (instead of being fixed at 0.3). This gives the bottom row of Fig. 2. For larger *n* and *m*, the distributions become less distinguishable. Consequently, it is also harder to estimate *n* and *m* exactly, however, if we are only interested in estimating the drift parameters, then for larger *n*, the exact value becomes less important.

### 2.3 The demographic model

In this section, we describe our statistical method for correctly estimating the number of individuals, the drift parameters, and the error rate.

Let us consider a fixed locus with a derived allele frequency *y* ∈ (0, 1) in a related population for which we have high-resolution allele frequencies. We call this population the anchor population. We assume that the ancient and anchor populations have a common ancestor. Since the common ancestor *t*_*C*_ generations have passed until the arrival of the anchor population, and *t*_*A*_ generations have passed until the ancient population, from which we have our eDNA or forensic sample. Let 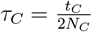 and 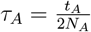 be the corresponding drift times, where *N*_*C*_ is the effective population size of the anchor population and *N*_*A*_ is the effective population size of the ancient population.

We assume that the frequency of the derived allele in the ancient population has a distribution *f*, which depends on *τ*_*C*_, *τ*_*A*_ and *y*. We denote this dependency by *f* (·) = *f* (·|*y, τ*_*A*_, *τ*_*C*_). We consider *f* later.

So, at this locus, in the ancient population, the frequency *x* of the derived allele is a realization of *f*. Given *x*, under the Hardy-Weinberg equilibrium, the 2*n* alleles of the *n* individuals have each, independently, a probability *x* of being derived, so the total number of derived alleles *k* is binomially distributed with parameters 2*n* and *x*, so the probability of *k* derived alleles is

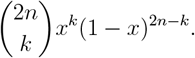

So with *k* derived alleles, a read has a chance of 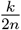 of being derived. However, through deamination or read errors, a derived read or allele might become (apparently) ancestral or vice versa, so the probability of a derived read with error rate *ε* becomes

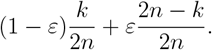

So the probability of *d* derived reads, conditioned on *k* derived alleles and error rate *ε*, with coverage *R* is binomially distributed with parameters *R* and 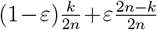. In Section S3.1 in the supplementary material, we argue that a separate estimation of the deamination and read error, under this model, is impossible. In addition, to avoid identifiability issues, we assume *ε*∈ [0, 1*/*2) (see Section S3.2 in the supplementary material). So, in a scheme, we have

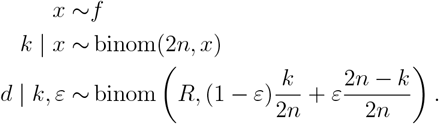

Schraiber [29] calculates the formula for the integral 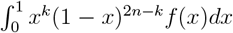, for all *n* and *k*. He found that the probability of *k* derived alleles out of 2*n*, given frequency *y* in the anchor population and drift parameters *τ*_*C*_ and *τ*_*A*_, to be

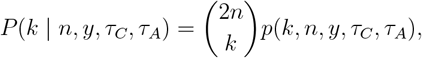

where

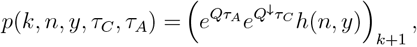

and *h*(*n, y*) is a real vector of length 2*n* + 1, defined as

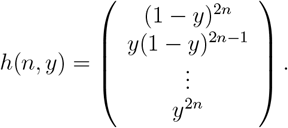

and *Q* and *Q*^*↓*^ are (2*n* + 1) ×(2*n* + 1) tri-diagonal matrices defined (for *k, 𝓁* ∈ {0, …, 2*n*}) by

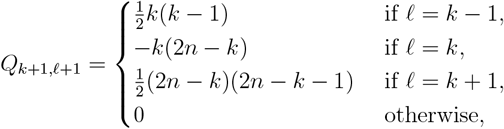

and

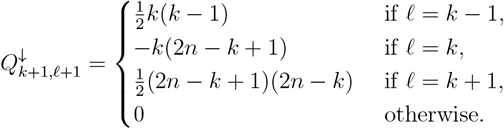

(Note that in mathematics, matrices and vectors traditionally start with index 1.) So for a given locus, the probability of *d* derived reads out of *R*, with *n* individuals, frequency *y* in the anchor population, *τ*_*C*_ and *τ*_*A*_ and error rate *ε*, is

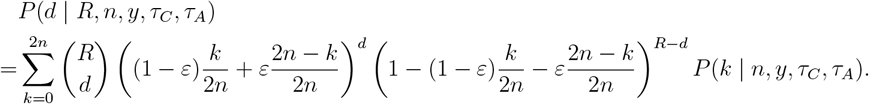

Schraiber [29] argues in the appendix of his article that loci with exactly 1X coverage do not contribute information about *τ*_*A*_. In addition, we argue in Section S3.3 that loci with 1X coverage also do not provide information about the number of individuals. Loci with coverage one only help to estimate *τ*_*C*_ and the error rate *ε*. These facts limit estimating *τ*_*A*_ and *n* with low coverage data, under this model.

### 2.4 The Metropolis sampler

As we pointed out at the start of this section, correct estimation of the number of individuals is crucial, as conditioning on different *n* leads to different distributions of the reads, even with the same frequency.

Early tests showed that incorrect estimation of *n* can lead to erroneous estimates of the drift parameters. When we set a uniform distribution on the drift parameters, the posterior concentrates around significantly different values of *τ*_*C*_ and *τ*_*A*_ for different *n*; this makes the correct estimation of *n* essential and simultaneously very challenging. Suppose at step *m* − 1 we are at position (*n*_*m−*1_, *τ*_*C,m−*1_, *τ*_*A,m−*1_), then if we were to propose a different *n*_*prop*_ with proposed *τ*_*C*_ and *τ*_*A*_ values close to *τ*_*C,m−*1_ and *τ*_*A,m−*1_ (as is, with high probability, done in random walk MCMC), then this proposal is almost certainly rejected, because the posterior conditioned on *n*_*prop*_ individuals concentrates around other *τ*_*C*_ and *τ*_*A*_ values. However, simulations show that conditioned on *n*, the posterior is unimodal. In conclusion, a naive symmetric random-walk Metropolis sampler leads to samples that do not mix and often estimate the wrong number of individuals. Typically, different chains get stuck at different *n*. To remedy this, we crafted an new MCMC sampler that can handle this mixture model.

The idea is as follows. For every *n*, we run a separate Metropolis sampler of the posterior distribution conditioned on having *n* individuals. To make this practical, we should assume that the number of individuals is bounded by some known *n*_max_. Then we have an extra sampler (*n*_*m*_)_*m≥*1_ for the posterior distribution of the number of individuals, where at step *m*, the proposal is accepted depending on the conditional drift values in the *n*_max_ other chains at step *m*. Finally, from these *n*_max_ + 1 chains we construct one unconditioned posterior sample, by taking *n*_*m*_ and the drift parameters and error estimate corresponding to conditioning on *n*_*m*_. So, if the conditioned chains are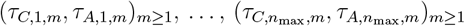, then we take 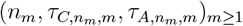 as the sample from our unconditioned posterior. The theoretical justification can be found in Theorem S5 in the supplementary material. The sampler is described in detail in Section S2.1 in the supplementary material.

#### 2.4.1 Software implementation

We implemented the model and the MCMC sampler in CHARON in the Julia programming language [3]. Installation, test data and usage can be found in the GitHub repository: https://github.com/Jan-van-Waaij/Charon. BAM files can be easily transformed in our native format using glactools [27] and scripts provided with the software.

It is possible to sample several chains in parallel. For this, one has to set the `--threads` flag of Julia (which requires Julia 1.5 or higher; for older versions of Julia, one has to set the `JULIA_NUM_THREADS` environment variable). For example, Julia could start with julia --threads=4 to run four chains in parallel.

For efficient calculation of the matrix exponent in the likelihood, we make use of the fact that one can factorize *Q* and *Q*^*↓*^ as *M* = *PDP* ^*−*1^, where *P* is an invertible matrix and *D* = diag(*d*_1_, …, *d*_*n*_) is a diagonal matrix with entries *d*_1_, …, *d*_2*n*+1_. Then *e*^*τM*^ = *PEP* ^*−*1^, where 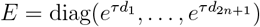 is a diagonal matrix with entries 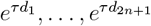).

## 3 Results

### 3.1 Simulations

To show the effectiveness of our method, we first tested it with simulations. We want to show that our method i) can estimate the number of individuals accurately. ii) can estimate the population parameters and the error rate accurately. iii) Pinpoint the most likely population that contributed to the sample. To achieve goals 1 and 2, we first simulate a simple population model with known drift parameters, see Section 3.1.1. For goal 3, we simulate a population history, see Section 3.1.2. For goals 1 and 3, we also constructed synthetic empirical data, see Section 3.2.

#### 3.1.1 Simple 2-pop simulations

In order to show the effectiveness of our method to estimate the correct number of individuals and the population parameters, we simulated a two-population history where the parameters are exactly known (previously described in Section 2.1.1). Using the simulations, we subsample the reads to simulate lower coverage by using sampling without replacement. Thus the distribution of the reads remains the same (see Lemma S2 in the supplementary material).

We apply our MCMC sampler and obtain a chain of 100,000 samples. We use the last 50,000, which are then thinned by using only every 10-th element of the sample, resulting in 5,000 draws from the posterior.

We calculate 95% credible intervals, by taking the 2.5% and 97.5% quantiles of the posterior sample. The results are shown in Figs. S2 to S5 in in the supplementary material. Each plot has credible intervals for coverage 1, 2, … 100. The black dot is the posterior median, and the horizontal coloured lines are the 95% credible intervals. The vertical blue line indicates the true simulated value.

For the estimation of *τ*_*C*_, our estimates seem robust down to 1X. The intervals tend to become slightly smaller for a larger number of individuals. For illustration, we only show here in Fig. 4 the plots for one and eight individuals. The other plots are shown in the supplementary material Fig. S2.

**Figure 4.**
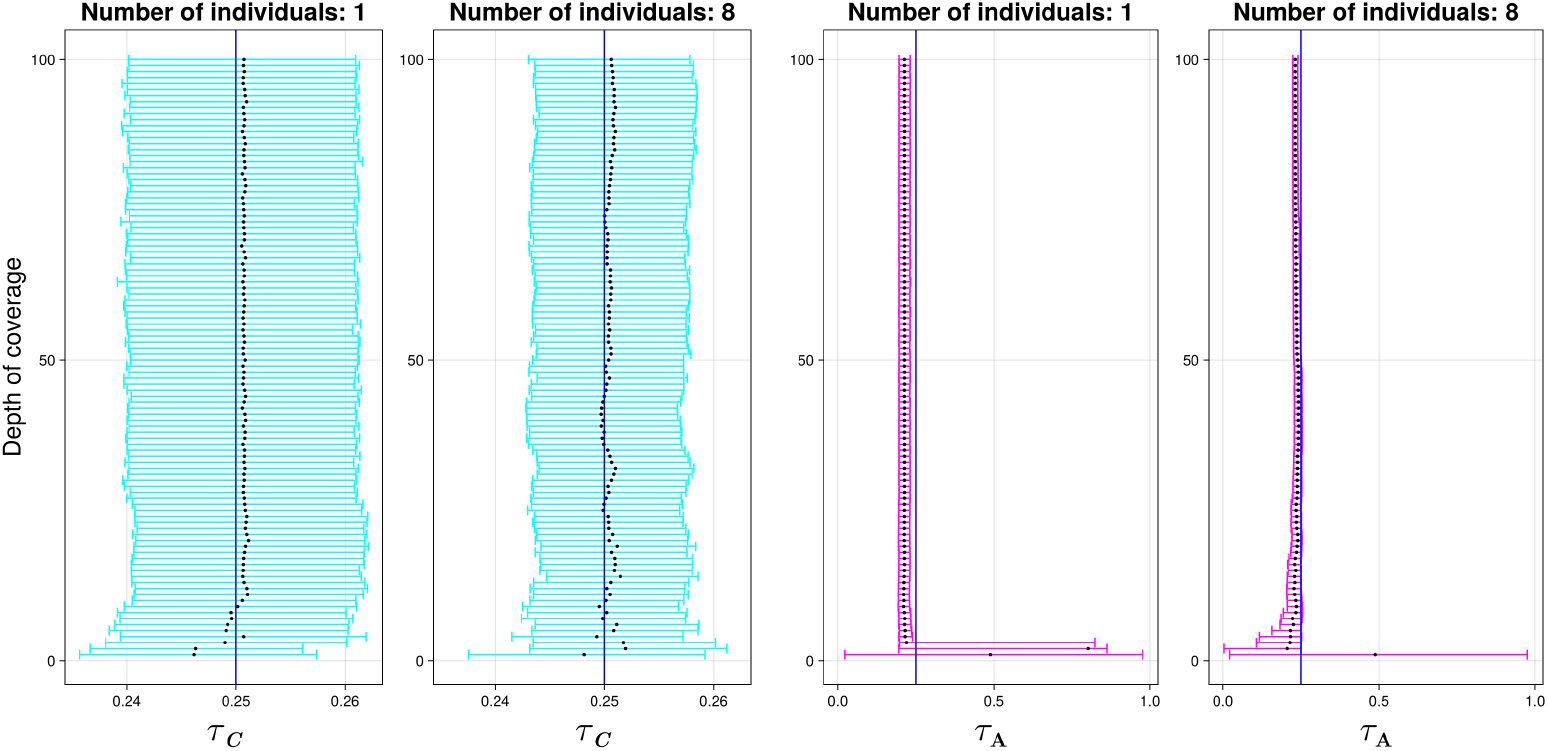
Credible intervals for *τ*_*C*_ (cyan) and *τ*_*A*_ (magenta), for 1 and 8 individuals, coverages 1,…,100. The blue line indicates the true simulated value. The black dot indicates the median.

For the estimation of *τ*_*A*_ the credible intervals get narrower as the depth of coverage increases. The number of individuals does not seem to affect the estimate. For illustration, we only show in Fig. 4 the case with one and eight individuals. In the supplement, all plots are shown, see Fig. S3.

The estimation of the number of individuals *n* converges to the right answer as coverage increases. For a small number of individuals (e.g. 1,2), we need depth of coverage of 2X-3X, for a larger number of individuals (e.g. 8), the uncertainty is higher and we require more than 50X for a perfect point estimate but roughly 10X to get a confidence interval of plus or minus 2 individuals. For the most difficult case, 8 individuals with 1X coverage, our estimate of the number of individuals is between 1 and 10, meaning that we cannot predict with certainty the number of individuals. The credible interval gets narrower and narrower around the correct value as coverage increases. For illustration, we only show here in Fig. 5 the plots for one and eight individuals. In the supplementary material Fig. S4, all plots are shown. Note that the credible intervals might be asymmetric and skewed because the posterior puts almost all mass on one value, and a little bit on either the smaller or higher values, and almost no mass on the remaining values, resulting in asymmetrical, skewed intervals.

**Figure 5.**
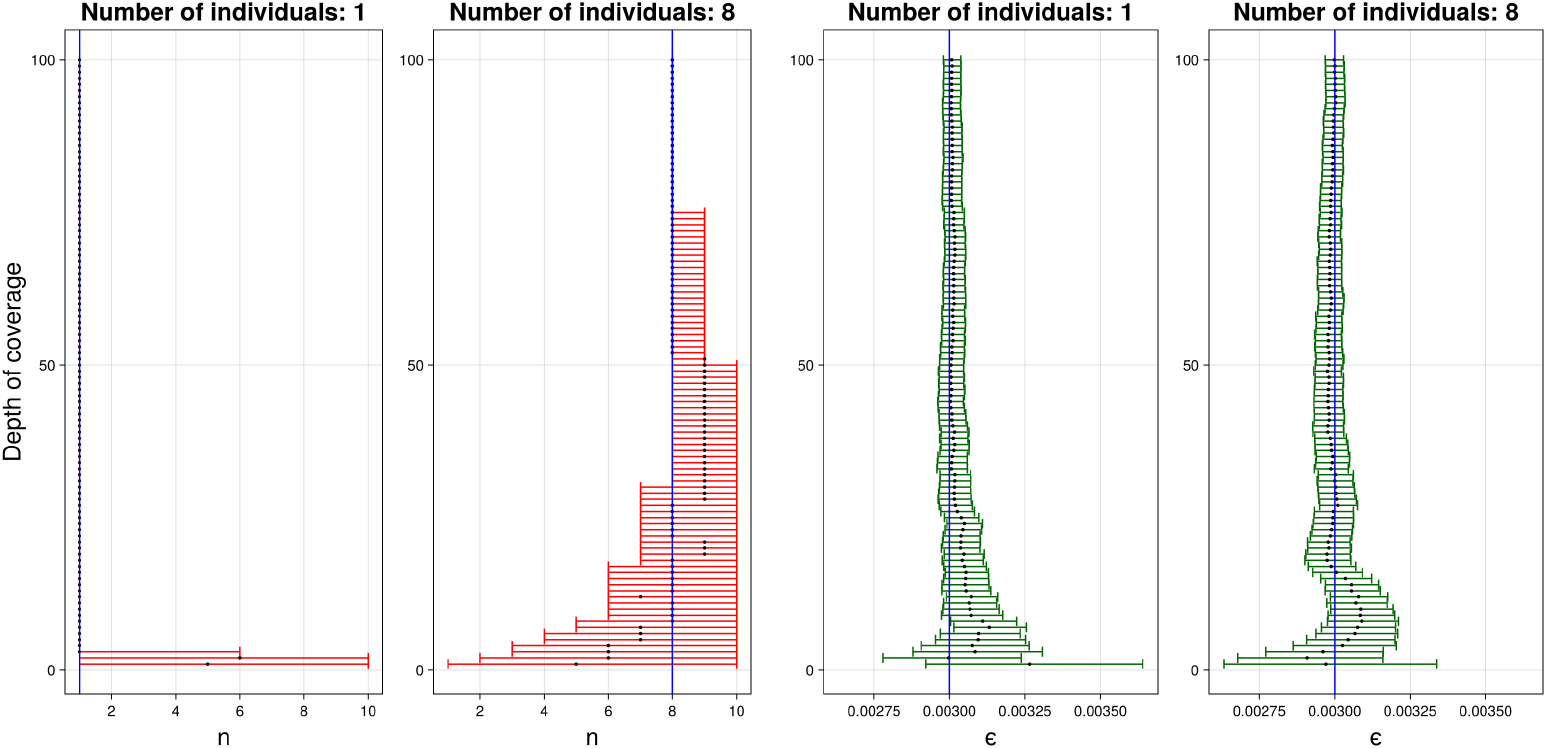
Credible intervals for *n* (red) and *ε* (green), for 1 and 8 individuals, coverages 1,…,100. The blue line indicates the true simulated value. The black dot indicates the median. For higher depth of coverage, the 95% credible intervals of *n* consist of only one point.

For the estimation of the error rate *ε*, there is no visual difference between the number of individuals. The estimates become narrower as the depth of coverage increases and converge around the correct value. For illustration, we only show here in Fig. 5 the plots for one and eight individuals. In the supplementary material Fig. S5 all plots are shown.

#### 3.1.2 Simulated multiple populations

Here we will show that our method is able to pinpoint the right population. We simulated a human population history with Demes and msprime as described in Section 2.1.1 and show that our method can accurately pinpoint the population and also estimate the correct number of individuals.

The human population history being simulated follows an out-of-Africa split, followed by a subsequent one in Eurasia and the settling of the Americas (see Fig. 6). We use the following abbreviations: **AMH**: Anatomically modern humans, **OOA**: Bottleneck out-of-Africa population, **YRI**: Yoruba in Ibadan, Nigeria, **YRIS**: Yoruba in Ibadan, Nigeria - Split, **CEU**: Utah residents with northern and western European ancestry, **CEUS**: Utah residents with northern and western European ancestry - split, **CHB**: Han Chinese in Beijing, China, **CHBS**: Han Chinese in Beijing, China - split, **KAR**: Karitiana native Brazilians, **KARS**: Karitiana native Brazilians - split.

**Figure 6.**
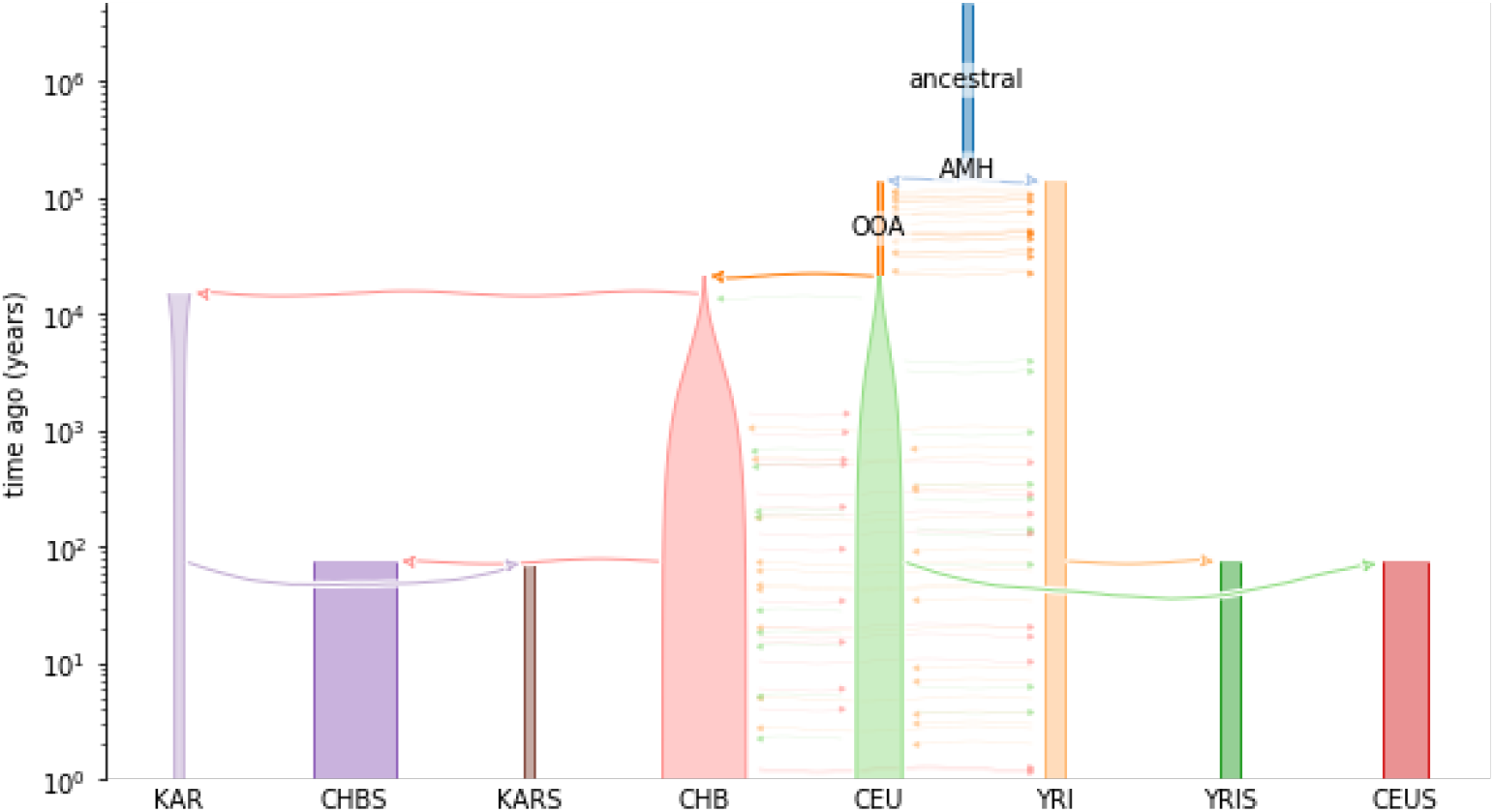
From the ancestral human population in Africa, we have different migrations and splits in new parts of the world. This is visualized here as the YRI staying in Africa, OOA being the emigration leading to the CEU, CHB and KAR populations. For this study, each population also has a split population used to generate the individual samples. We used a split of 75 years (3 generations) ago for each sample. This is done to simulate the passage of time between the split from the population in the sample versus the anchor population.

We took the populations KAR, CHB, CEU, and YRI as anchor populations, and we simulated eDNA from the populations CHBS, KARS, YRIS and CEUS, which split from their respective populations 75 years (3 generations) ago. We estimated the confidence interval for each of the combinations of populations with coverage 5, 10, …, 100. For the number of individuals, we took 1, 2, …, 8. The simulated error rate is 0.003. The plots can be found in Figs. S10 to S73 in the supplementary material.

The number of individuals and the error rate are correctly estimated. The credible intervals for the number of individuals can be found in Figs. S13, S17, S21, S25, S29, S33, S37, S41, S45, S49, S53, S57, S61, S65, S69 and S73 in the supplementary materialand the plots for the credible intervals of the error rate can be found in Figs. S12, S16, S20, S24, S28, S32, S36, S40, S44, S48, S52, S56, S60, S64, S68 and S72 in the supplementary material. In the data with three individuals and coverage 20X, the posterior correctly places more than 95% of its mass on *n* = 3. The credible intervals for the estimation error are given in Fig. 7. We see from Fig. 7 that our method accurately infers the short distance both in terms of *τ*_*A*_ and *τ*_*C*_ between CEU and CEUS. The plots for the other sample populations can be found in Figs. S6 to S9. For *τ*_*A*_, we are mostly dependent on the split time hence the higher value for YRI. For *τ*_*C*_, we are both dependent on the split time and effective population size in the anchor population. Due to the lower effective population size in the KAR, we see the highest value for this parameter followed by CHB and YRI. This indicates that we can use our method to determine the population that is the genetically closest one to the eDNA sample.

**Figure 7.**
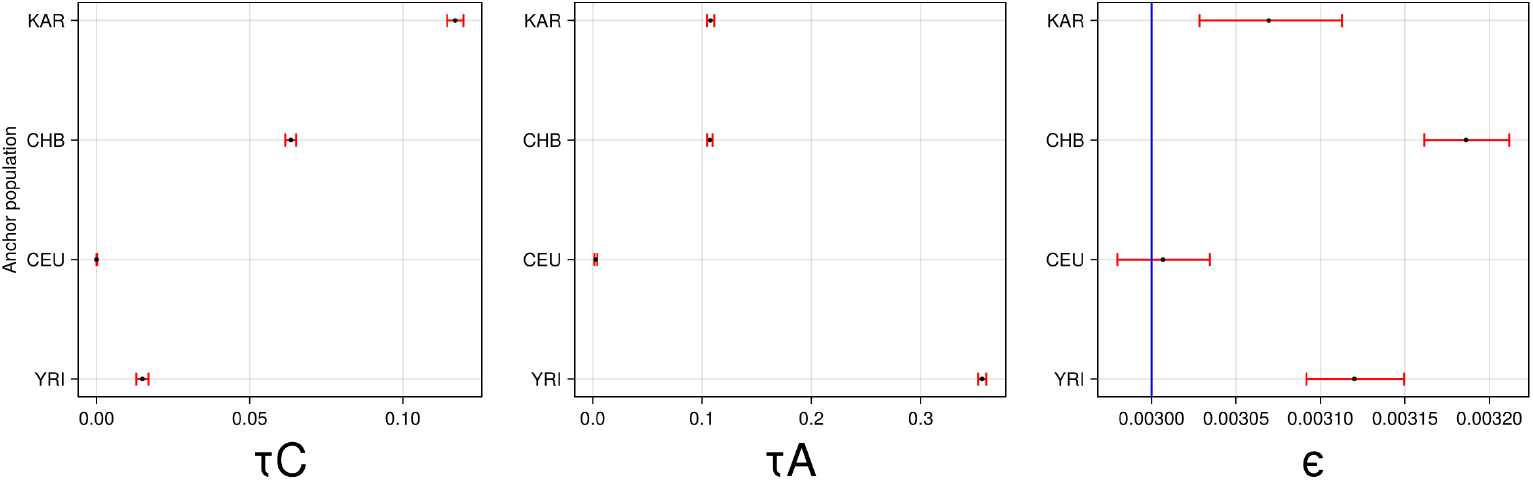
Credible intervals for the drift in the branch leading to the sample *τ*_*A*_ (left) and drift in the branch of the anchor population *τ*_*C*_ (middle), with *n* = 3 individuals and a coverage of 20X. To the right is the credible interval for *ε*. The blue line is the simulated true error level.

### 3.2 Synthetic empirical data

Using the synthetic empirical data described in Section 2.1.2, we sought to determine if our method could identify the number of individuals as well as the source population. For the latter, we use different statistics to determine the shortest distance to the sediment population from several anchor populations, that is, the median of *τ*_*C*_, *τ*_*A*_, max {*τ*_*A*_, *τ*_*C*_} and *τ*_*A*_ + *τ*_*C*_.

The results of the different population drift statistics are in Fig. S75 in the supplementary material. Here, in Fig. 8, we only show the results for *τ*_*C*_ + *τ*_*A*_. In general, we see that when considering *τ*_*C*_ + *τ*_*A*_ hence the total drift between the sample and anchor population, the population with the lowest value is IBS followed by either Tuscans (TSI) or Northern European (CEU) whereas the highest values are for the Mende (MSL) and Esan (ESN). These results are expected due to the demographic history. Regardless of the number of individuals, IBS was always the population with the lowest drift values.

**Figure 8.**
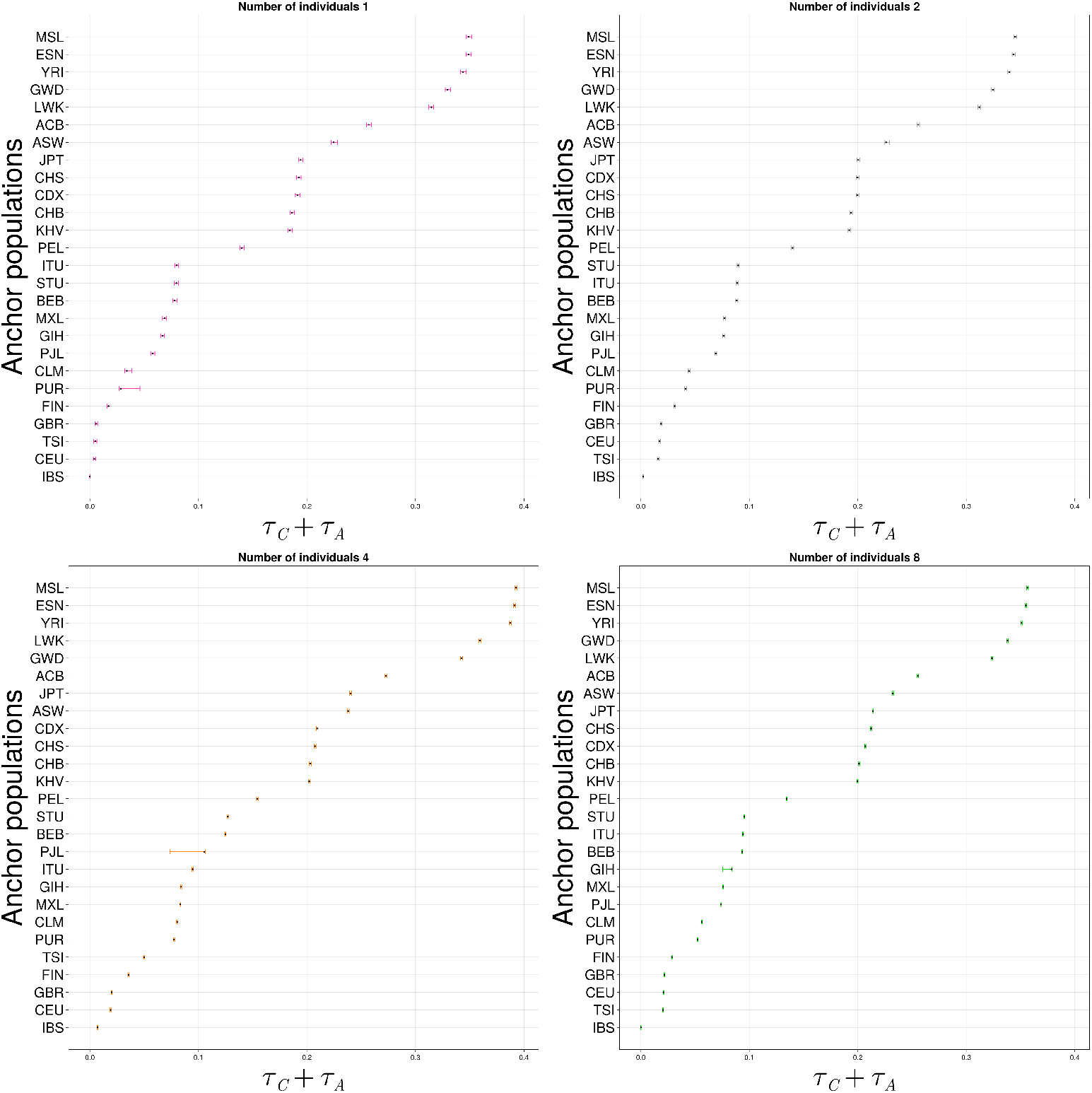
Populations ordered by *τ*_*C*_ + *τ*_*A*_ for the samples simulated with IBS as source population, respectively, along with 95% credible intervals. The black dot indicates the posterior median.

We also estimate the number of individuals (see Fig. 9). We see that our estimates are correct for the number of individuals for *n* = 1, 2 regardless of the anchor population. At *n* = 4, our estimate is mostly correct, especially for populations with lower drift values (e.g. IBS, TSI, CEU). These estimates are off by one for the population with higher drift values (e.g. ESN, MSL), probably due to the increasing lack of allele frequency resolution at high drift. At *n* = 8, we often overestimate by 1 or 2 for most populations but using IBS as anchor gives us the correct estimate.

**Figure 9.**
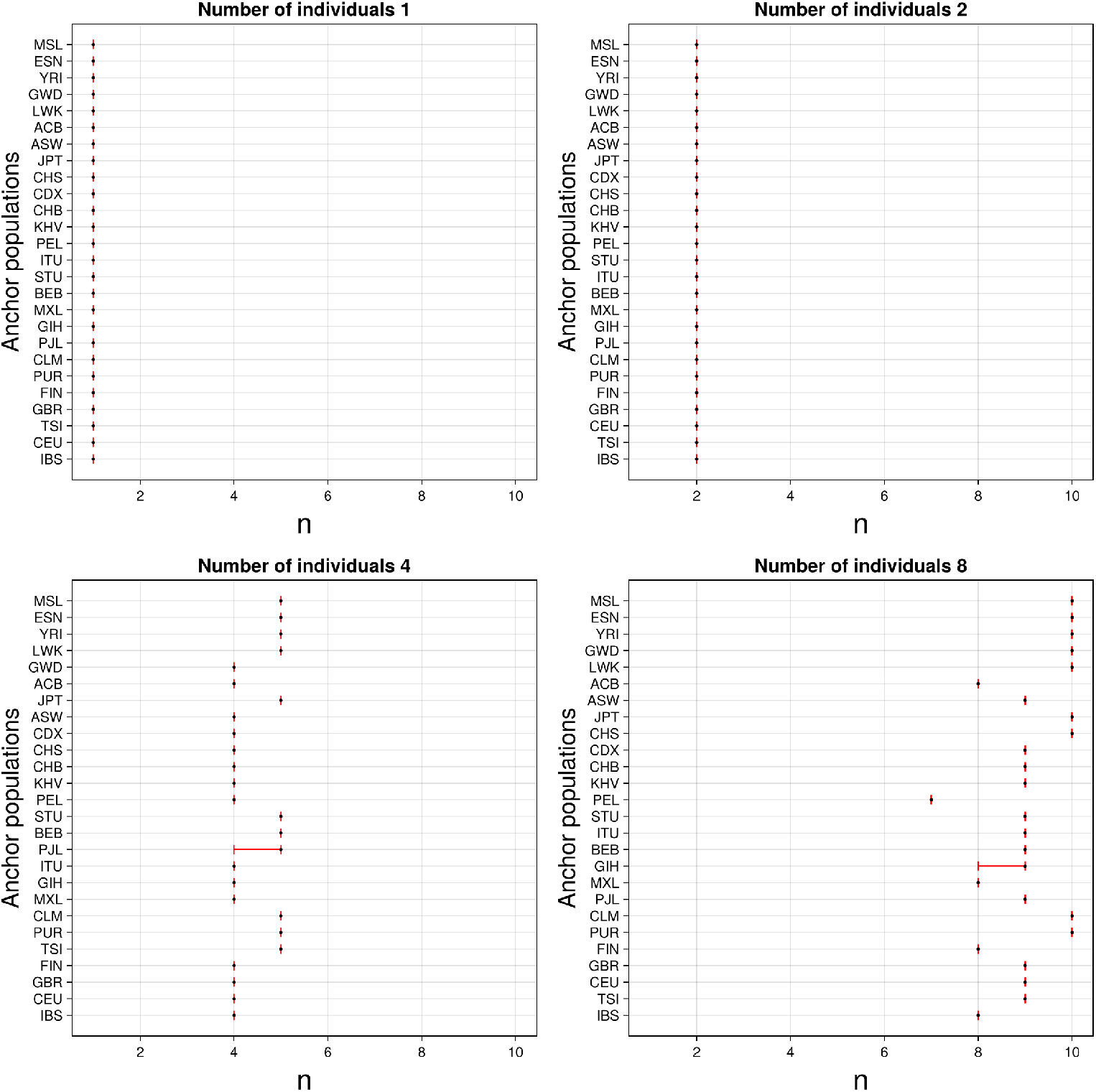
Credible intervals for the number of individuals, except for a few cases, the posterior puts almost all its mass in one point. The black dot indicates the posterior median.

Thus, for small values of *n*, our method accurately recovers the number of individuals. Moreover, it consistently identifies the correct population structure, with drift patterns aligning with known demographic histories. The credible intervals for the error rates *ε* are in Fig. S74 in the supplementary material.

## 4 Discussion

In this study, we introduced CHARON, a novel statistical method designed to address two significant challenges in environmental and forensic DNA analyses: accurately estimating the number of contributing diploid individuals and inferring genetic drift times between a sampled population and known reference populations. Using allele frequency distributions from known populations, CHARON provides a statistical framework for jointly estimating these parameters.

Our simulation studies demonstrated that CHARON accurately estimates the number of individuals contributing to DNA mixtures at mid-coverage (above ∼3X for a 1-2 of individuals, above 8-10X for 8 individuals), distinguishing up to eight diploid individuals. Beyond this, CHARON runs into numerical and statistical issues. Despite this, as the coverage increases, the credible intervals for estimates of drift parameters (*τ*_*C*_ and *τ*_*A*_), the error rate, and the number of individuals narrow, confirming the robustness of the model.

When applied to simulated population data, CHARON effectively pinpointed the correct population of origin. It accurately inferred smaller drift times between closely related populations (e.g., CEU and its split CEUS) while providing wider drift intervals for more genetically distant populations. Using synthetic empirical data constructed from the 1000 Genomes Project, CHARON correctly identified the population of origin (IBS) and accurately estimated the number of individuals across different simulated scenarios. The reasonable depth of coverage inherent to these synthetic mixtures (8X-10X), showed that our method is applicable to empirical human eDNA and forensic contexts. It is plausible that even ancient sediments will yield enough human DNA to allow quantification of the population of origin and estimate the number of individuals.

However, CHARON assumes equal contributions from all individuals and a homogeneous genetic background within the sampled population. These assumptions, while practical, could limit applicability in scenarios involving highly skewed individual contributions or heterogeneous samples. Future work might explore extensions of CHARON to account for unequal DNA contributions or mixtures from genetically diverse populations.

## Supporting information

Supplementary figures and explanation

## 5 Acknowledgements

First, we would like to thank Fernando Racimo and Joshua Schraiber for their help and comments. We would also like to thank Graham Gower for his assistance in simulating the populations. This research was funded by the Novo Nordisk Data Science Investigator grant number NNF20OC0062491 (JvW, MVH, GR). Additional funding for computational resources was provided by the Department of Health Technology at the Technical University of Denmark (DTU). The funders had no role in study design, data collection and analysis, the decision to publish, or preparation of the manuscript. GR acknowledges using OpenAI/ChatGPT for polishing the text of this manuscript.

## Notes

### Competing Interest Statement

The authors have declared no competing interest.

## References

[1] J. S. Allwood, N. Fierer, and R. R. Dunn. “The future of environmental DNA in forensic science”. In: Applied and Environmental Microbiology 86.2 (2020), e01504–19.

[2] K. C. Beng and R. T. Corlett. “Applications of environmental DNA (eDNA) in ecology and conservation: opportunities, challenges and prospects”. In: Biodiversity and conservation 29.7 (2020), pp. 2089–2121.

[3] J. Bezanson et al. “Julia: A fresh approach to numerical computing”. In: SIAM review 59.1 (2017), pp. 65–98. url: 10.1137/141000671.

[4] J. M. Butler. “The future of forensic DNA analysis”. In: Philosophical transactions of the royal society B: biological sciences 370.1674 (2015), p. 20140252.

[5] B. Chapman et al. “An environmental DNA approach to the isolation of human intra- and extra-cellular DNA from large volumes of water from crime scenes”. In: Forensic Genomics 3.3 (2023), pp. 94–102.

[6] 1000 Genomes Project Consortium et al. “A global reference for human genetic variation”. In: Nature 526.7571 (2015), p. 68.

[7] Popsim Consortium. Demes specification documentation: Pre-made models. Accessed: 2024-10-22. 2022. url: https://popsim-consortium.github.io/demes-spec-docs/main/introduction.html.

[8] M. E. Cristescu and P. D. N. Hebert. “Uses and misuses of environmental DNA in biodiversity science and conservation”. In: Annual Review of Ecology, Evolution, and Systematics 49.1 (2018), pp. 209–230.

[9] M. A. Dass et al. “Assessing eDNA capture method from aquatic environment to optimise recovery of human mt-eDNA”. In: Forensic Science International (2024), p. 112085.

[10] M. A. Dass et al. “Assessing the use of environmental DNA (eDNA) as a tool in the detection of human DNA in water”. In: Journal of Forensic Sciences 67.6 (2022), pp. 2299–2307.

[11] K. Deiner, H. Yamanaka, and L. Bernatchez. “The future of biodiversity monitoring and conservation utilizing environmental DNA”. In: Environmental DNA 3.1 (2021), pp. 3–7.

[12] C. Fantinato, P. Gill, and A. E. Fonneløp. “Investigative use of human environmental DNA in forensic genetics”. In: Forensic Science International: Genetics 70 (2024), p. 103021.

[13] M. Goray et al. “Emerging use of air eDNA and its application to forensic investigations–a review”. In: Electrophoresis 45.9-10 (2024), pp. 916–932.

[14] G. Gower et al. “Demes: A standard format for demographic models”. In: Genetics 222.3 (2022).

[15] R. N. Gutenkunst et al. “Inferring the Joint Demographic History of Multiple Populations from Multidimensional SNP Frequency Data”. In: PLOS Genetics 5.10 (2009), e1000695. doi: 10.1371/journal.pgen.1000695.

[16] M. D. Johnson et al. “Airborne eDNA reflects human activity and seasonal changes on a landscape scale”. In: Frontiers in Environmental Science 8 (2021), p. 563431.

[17] J. Kelleher et al. “Efficient ancestry and mutation simulation with msprime 1.0”. In: Genetics 220.3 (2020).

[18] M. Lewis et al. “The forensic potential of environmental DNA (eDNA) in freshwater wildlife crime investigations: From research to application”. In: Science & Justice (2024).

[19] J. Lindo et al. “A time transect of exomes from a Native American population before and after European contact”. In: Nature Communications 7 (2016), p. 13175. doi: 10.1038/ncomms13175.

[20] C. Neves and M. Zieger. “Total Human DNA Sampling–Forensic DNA profiles from large areas”. In: Forensic Science International: Genetics 67 (2023), p. 102939.

[21] L. Orlando et al. “Ancient DNA analysis”. In: Nature reviews methods primers 1.1 (2021), p. 14.

[22] B. S. Pedersen and A. R. Quinlan. “Mosdepth: quick coverage calculation for genomes and exomes”. In: Bioinformatics 34.5 (2018), pp. 867–868.

[23] S. Peyrégne and B. M. Peter. “AuthentiCT: a model of ancient DNA damage to estimate the proportion of present-day DNA contamination”. In: Genome biology 21.1 (2020), p. 246.

[24] D. Primorac and M. Schanfield. Forensic DNA applications: An interdisciplinary perspective. CRC press, 2023.

[25] K. Prüfer et al. “Computational challenges in the analysis of ancient DNA”. In: Genome biology 11 (2010), pp. 1–15.

[26] F. Racimo, G. Renaud, and M. Slatkin. “Joint Estimation of Contamination, Error and Demography for Nuclear DNA from Ancient Humans”. In: PLOS Genetics 12.4 (2016), pp. 1–27. doi: 10.1371/journal.pgen.1005972.

[27] G. Renaud. “glactools: a command-line toolset for the management of genotype likelihoods and allele counts”. In: Bioinformatics 34.8 (2018), pp. 1398–1400.

[28] G. Renaud et al. “Schmutzi: estimation of contamination and endogenous mitochondrial consensus calling for ancient DNA”. In: Genome Biology 16 (2015). doi: 10.1186/s13059-015-0776-0.

[29] J. G. Schraiber. “Assessing the Relationship of Ancient and Modern Populations”. In: Genetics 208.1 (2018), pp. 383–398. issn: 1943-2631. doi: 10.1534/genetics.117.300448.

[30] M. Shahzad et al. “Evaluation of storage conditions and the effect on DNA from forensic evidence objects retrieved from lake water”. In: Genes 15.3 (2024), p. 279.

[31] J. D. Wells and V. Skaro. “Application of DNA-based methods in forensic entomology”. In: Forensic DNA Applications (2023), pp. 285–304.

[32] K. E. Williams, K. P. Huyvaert, and A. J. Piaggio. “Clearing muddied waters: Capture of environmental DNA from turbid waters”. In: PLoS ONE 12.7 (2017), e0179282.

