## Supplementary figures and explanation for "CHARON: Estimating the drift time and the number of individuals in environmental DNA with diploid individuals"

#### S1 The distribution of the reads

Given a frequency $x$ of a derived allele at a particular locus and $n$ individuals, the number of derived alleles $k$ is, under Hardy-Weinberg equilibrium, binomially distributed with parameters $2n$ and $x$. With coverage $R$ and $k$ derived alleles, the number of derived reads $d$ is (ignoring errors) binomially distributed with parameters $R$ and $\frac{k}{2n}$. So one might wonder whether the unconditional distribution of $d$ (so $d$ is only conditioned on $x$, $n$, and $R$, but not on $k$) simplifies to a binomial distribution. However, that is not the case, as the following lemma shows.

**Lemma S1.** *Let $n$ and $r$ be positive integers with $r \geq 2$, and let $p \in (0,1)$. Let $X$ be binomially distributed with parameters $n$ and $p$. Let $Y$ be a random variable, so that $Y \mid X$ is binomially distributed with parameters $r$ and $\frac{X}{n}$. Then, $Y$ is not binomially distributed (for any pair of parameters).*

*Proof.* We argue by contradiction that $Y$ is not binomially distributed. Let us assume that $Y$ is binomially distributed. Note that for every $x$, $Y \mid X = x$ takes values in (a subset of) $\{0, \dots, r\}$. So $Y$ takes values in $\{0, \dots, r\}$. Moreover, for $m \in \{0, \dots, r\}$,

$$\begin{aligned} P(Y = m) \geq & P(Y = m \mid X = 1) P(X = 1) \\ = & \left(\frac{1}{n}\right)^m \binom{n}{1} p (1-p)^{n-1} > 0, \end{aligned}$$

as $0 < p < 1$. So $Y$ has support $\{1, \dots, r\}$. So, the number-of-experiments-parameter of the binomial distribution of $Y$ is $r$.

Note that $\mathbb{E}\, Y = \mathbb{E}[\mathbb{E}[Y \mid X]] = \mathbb{E}\left[r \cdot \frac{X}{n}\right] = \frac{r}{n}\, \mathbb{E}\, X = \frac{r}{n} np = rp$. It follows that the probability-of-succes-parameter of $Y$ is $p$. However,

$$\begin{aligned} \operatorname{var}(Y) = & \mathbb{E}[\operatorname{var}(Y \mid X)] + \operatorname{var}(\mathbb{E}[Y \mid X]) \\ = & \mathbb{E}\left[r \cdot \frac{X}{n} \cdot \left(1 - \frac{X}{n}\right)\right] + \operatorname{var}\left(r \cdot \frac{X}{n}\right) \\ = & \frac{r}{n}\, \mathbb{E}[X] - \frac{r}{n^2}\, \mathbb{E}[X^2] + \frac{r^2}{n^2} np(1-p) \\ = & \frac{r}{n} np - \frac{r}{n^2}\left(np(1-p) + (np)^2\right) + \frac{r^2}{n} p(1-p) \\ = & rp - \frac{1}{n} rp(1-p) - rp^2 + \frac{r^2}{n} p(1-p) \\ = & rp(1-p) + \frac{r(r-1)}{n} p(1-p) \\ > & rp(1-p), \end{aligned}$$

as $r > 1$ and $p \notin \{0, 1\}$. However, $\operatorname{var}(Y) > rp(1-p)$ is in contradiction with the fact that the binomial distribution with parameters $r$ and $p$ has variance $rp(1-p)$. So, $Y$ is not binomially distributed. □

**Lemma S2.** *Let $R$ be a positive integer. Let $f$ be a probability density on the interval $[0, 1]$. Given $f$, and $n \in \mathbb{N}$, consider the following distribution*

$$\begin{aligned} x \sim & f, \\ k \mid x \sim & \mathit{binom}(2n, x), \\ r_1, \dots, r_R \sim & \mathit{Bernoulli}(k/(2n)). \end{aligned} \tag{1}$$

*Define*

$$S_R = \sum_{i=1}^{R} r_i.$$

*Let $R_1, R_2$ be positive integers so that $R_1 < R_2$. Let $r_1, \dots, r_{R_2}$ be generated according to Eq. (1). Sample $R_1$ items $i_1, \dots, i_{R_1}$ from $\{1, \dots, R_2\}$ without replacement. Define*

$$T = \sum_{j=1}^{R_1} r_{i_j}.$$

*Then $T$ has the same distribution as $S_{R_1}$.*

*Proof.* Note that given $k$, $r_1, \dots, r_{R_2}$ are independent random variables. So, if you pick randomly $R_1$ of them, this has the same distribution as if you sampled $R_1$ in one go. So $S_{R_1}$ and $T$ have the same distribution. □

Let's introduce the notion of convergence in total variation, which implies the more familiar convergence in distribution [5, section 2.9].

**Definition S3.** A sequence of random variables $(X_n)$ converges in total variation to a random variable $X$ when

$$\sup_B |P(X_n \in B) - P(X \in B)| \to 0, \text{ as } n \to \infty,$$

where the supremum is taken over all measurable sets $B$.

**Lemma S4.** *Let $r \in \mathbb{N}$ and $0 \leq p \leq 1$. Let $d_n$ be defined as follows*

$$k \sim binom\,(n, p)$$
$$d_n \mid k \sim binom\left(r, \frac{k}{n}\right).$$

*Then $d_n$ converges in total variation to the binom$(r, p)$-distribution, as $n \to \infty$.*

*Proof.* Let $f_n$ be the distribution of $d_n$ and $f_n(\cdot \mid k)$ the distribution of $d_n$ conditioned on $k$. Let $d \in \{0, \dots, r\}$. Then

$$f_n(d \mid k) = \binom{r}{d} \left(\frac{k}{n}\right)^d \left(1 - \frac{k}{n}\right)^{r-d}.$$

So

$$f_n(d) = \binom{r}{d} \sum_{k=0}^{2n} \left(\frac{k}{n}\right)^d \left(1 - \frac{k}{n}\right)^{r-d} \binom{n}{k} p^k (1-p)^{n-k}$$

Define $g(y) = \left|y^d(1-y)^{r-d}\right| \wedge 1$ (for real numbers $a, b$, $a \wedge b$ is the minimum of $a$ and $b$). Then $g : \mathbb{R} \to \mathbb{R}$ is a Lipschitz function and $f_n(d) = \binom{r}{d} \mathbb{E}\, g(X_n/n)$ where $X_n$ is binomially distributed with parameters $n$ and $p$. By the law of large numbers, $X_n/n$ converges almost certainly to $p$. As $g$ is bounded and continuous, $\mathbb{E}\, g(X_n/n)$ converges, according to Portmanteau's lemma [5, Lemma 2.2], to $g(p)$.

So $f_n(d)$ converges to

$$\binom{r}{d} p^d (1-p)^{r-d}.$$

Hence, by Scheffé's theorem [5, corollary 2.30], $d_n$ converges in total variation to the binomial distribution with parameters $r$ and $p$, as $n \to \infty$. □

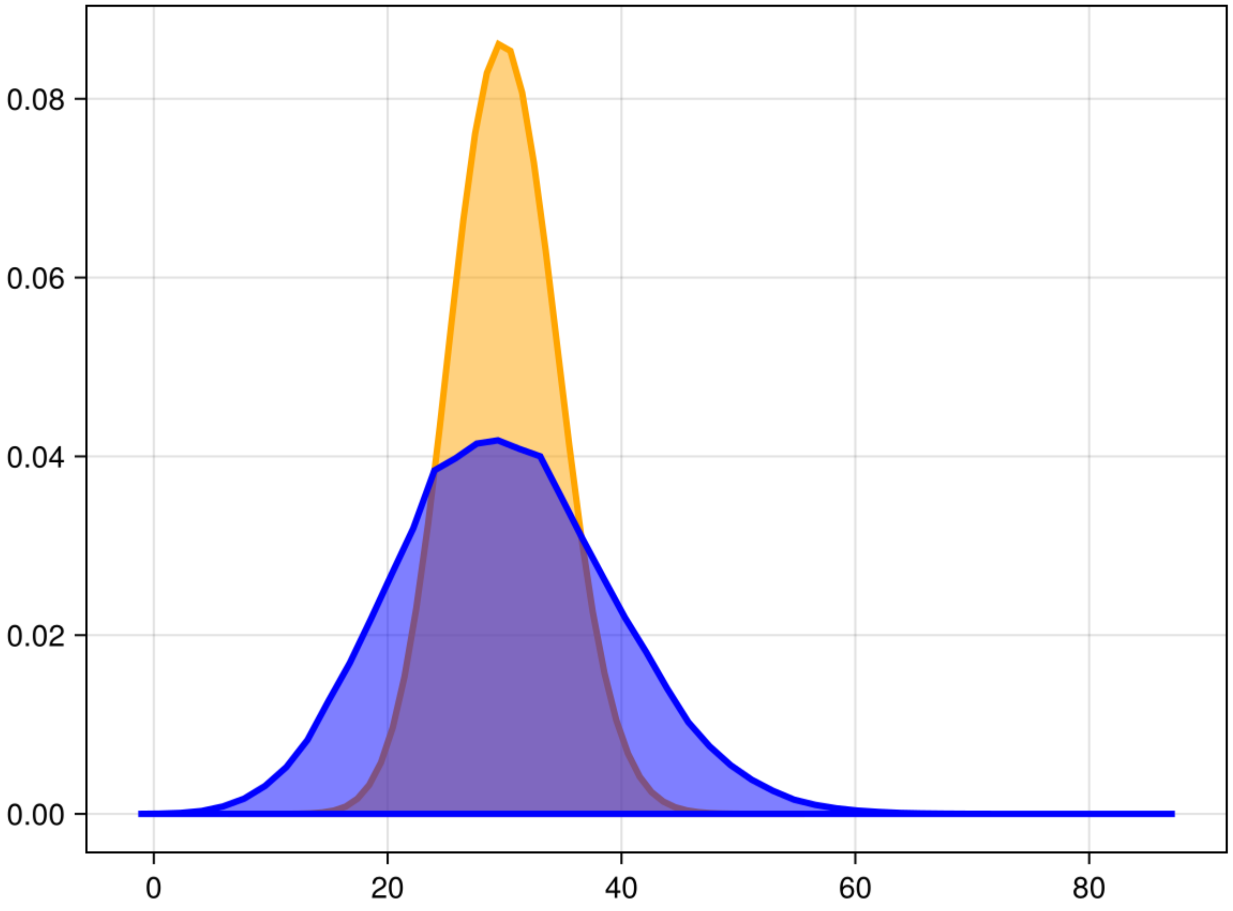

Figure S1: Distribution of the reads with a probability of derived allele 0.3, 16 individuals, and coverage 100 (blue graph) vs. the distribution of a binom$(100, 0.3)$-distribution. So the distribution of the reads has heavier tails than the binomial distribution.

#### S2 The sampler

In the following theorem, we prove that our MCMC sampler converges to the posterior distribution.

**Theorem S5.** *Let $n_{\max} \in \mathbb{N}$. Let $f(n, \theta)$ be a density on $\{1, \ldots, n_{\max}\} \times \mathbb{R}^d$. Let $f(\theta \mid n)$ be the distribution of $f$ conditioned on $n$. Let $(n_m, \theta_{1,m}, \ldots, \theta_{n_{\max},m})_m$ be a Markov chain so that for each $n \in \{1, \ldots, n_{\max}\}$, $(\theta_{n,m})_m$ is an ordinary Metropolis-Hastings Markov chain with stationary distribution $f(\theta \mid n)$. For given $n_{m-1}$, sample $n_m$ from a proposal distribution $q(n_m \mid n_{m-1})$, and accept $n_m$ with probability*

$$\min\left(\frac{f(n_m, \theta_{n_m,m})}{f(n_{m-1}, \theta_{n_{m-1},m})} \frac{q(n_{m-1} \mid n_m)}{q(n_m \mid n_{m-1})}, 1\right)$$

*Then $f$ is the invariant distribution of $(n_m, \theta_{n_m,m})$; in other words, if $(n_{m-1}, \theta_{n_{m-1},m-1})$ is distributed according to $f$, then $(n_m, \theta_{n_m,m})$ is also distributed according to $f$.*

*Proof.* We follow section 7.3 in [2] for the proof.

Note that support of the proposal distribution includes the support of the posterior.

Note that for each $n$, $(\theta_{n,m})_m$ is an ordinary Metropolis-Hastings Markov chain and thus converges to the invariant distribution $f(\cdot \mid n)$, and for each $n$, the transition density $K_n$ satisfies the detailed balance equation $K_n(\theta, \theta')f(\theta \mid n) = K_n(\theta', \theta)f(\theta' \mid n)$, for all $\theta, \theta' \in \mathbb{R}^d$.

The transition probability of $(n_m)_m$ depends on $(\theta_{1,m}, \dots, \theta_{n_{\max},m})$ and for $n' \neq n$ the transition probability of going from $n$ to $n'$ is given by

$$K(n, n' \mid \theta'_1, \dots, \theta'_n) = \min\left(\frac{f(n', \theta'_{n'})}{f(n, \theta'_n)}\frac{q(n \mid n')}{q(n' \mid n)}, 1\right) q(n' \mid n).$$

First consider the case $\frac{f(n',\theta'_{n'})}{f(n,\theta'_n)}\frac{q(n|n')}{q(n'|n)} \leq 1$, then $K(n', n \mid \theta'_1, \dots, \theta'_n) = q(n \mid n')$. So

$$K(n, n' \mid \theta'_1, \dots, \theta'_n)f(n, \theta'_n) = f(n', \theta'_{n'})q(n \mid n') = K(n', n \mid \theta'_1, \dots, \theta'_n)f(n', \theta'_{n'}).$$

When $\frac{f(n',\theta'_{n'})}{f(n,\theta'_n)}\frac{q(n|n')}{q(n'|n)} > 1$, then $K(n, n' \mid \theta'_1, \dots, \theta'_n) = q(n' \mid n)$ and

$$K(n', n \mid \theta'_1, \dots, \theta'_n) = \frac{f(n, \theta'_n)}{f(n', \theta'_{n'})}q(n' \mid n),$$

So

$$\begin{aligned}
&K(n, n' \mid \theta'_1, \dots, \theta'_n)f(n, \theta'_n) \\
=&f(n, \theta'_n)q(n' \mid n) \\
=&\frac{f(n, \theta'_n)}{f(n', \theta'_{n'})}q(n' \mid n)f(n', \theta'_{n'}) \\
=&K(n', n \mid \theta'_1, \dots, \theta'_n)f(n', \theta'_{n'}).
\end{aligned}$$

When $n' = n$, then

$$K(n, n' \mid \theta'_1, \dots, \theta'_n)f(n, \theta'_n) = K(n', n \mid \theta'_1, \dots, \theta'_n)f(n', \theta'_{n'})$$

holds trivially. Summarising, for all $n, n'$, and $\theta'$,

$$K(n, n' \mid \theta'_1, \dots, \theta'_n)f(n, \theta'_n) = K(n', n \mid \theta'_1, \dots, \theta'_n)f(n', \theta'_{n'})$$

So we have

$$\begin{aligned}
&K(n, n' \mid \theta'_1, \dots, \theta'_n)K_n(\theta_n, \theta'_n)f(n, \theta_n) \\
=&K(n, n' \mid \theta'_1, \dots, \theta'_n)K_n(\theta_n, \theta'_n)f(\theta_n \mid n)f_n(n) \\
=&K(n, n' \mid \theta'_1, \dots, \theta'_n)K_n(\theta'_n, \theta_n)f(\theta'_n \mid n)f_n(n) \\
=&K(n, n' \mid \theta'_1, \dots, \theta'_n)K_n(\theta'_n, \theta_n)f(n, \theta'_n) \\
=&K(n', n \mid \theta'_1, \dots, \theta'_n)K_n(\theta'_n, \theta_n)f(n', \theta'_{n'}).
\end{aligned}$$

Suppose $(n_{m-1}, \theta_{n_{m-1},m-1}) \sim f$. Let $n' \in \{1, \dots, n_{\max}\}$. Let $B \subseteq \mathbb{R}^n$ be a measurable set. Then

$$\begin{aligned}
&P((n_m, \theta_{n_m,m}) \in \{n'\} \times B) \\
=&\int_{\theta' \in B} \sum_{n=1}^{n_{\max}} \int_{\theta \in \mathbb{R}^d} K(n, n' \mid \theta'_1, \dots, \theta'_n)K_n(\theta, \theta')f(n, \theta)d\theta d\theta' \\
=&\int_{\theta' \in B} \sum_{n=1}^{n_{\max}} \int_{\theta \in \mathbb{R}^d} K(n', n \mid \theta'_1, \dots, \theta'_n)K_n(\theta', \theta)f(n', \theta')d\theta d\theta' \\
=&\int_{\theta' \in B} \sum_{n=1}^{n_{\max}} K(n', n \mid \theta'_1, \dots, \theta'_n)f(n', \theta')d\theta' \\
=&\int_{\theta' \in B} f(n', \theta')d\theta'.
\end{aligned}$$

So $f$ is the invariant measure of $(n_m, \theta_{n_m,m})$. □

#### S2.1 A description of the sampler

Let us describe the sampler in detail. We do Bayesian inference for the parameters $n, \tau_C, \tau_A$ and the error rate $\varepsilon$. We equip $n$, $\tau_C, \tau_A$ and $\varepsilon$ with priors. For our implementation, we need the prior on $n$ to have finite support; that is, there is an $n_{\max}$ so that $P(n \leq n_{\max}) = 1$. We allow for correlation between $\tau_C$ and $\tau_A$, so take a prior on the pair $(\tau_C, \tau_A)$.

Let, at step $m-1$, the $n_{\max}+1$ chains be at states

$$n_{m-1}, (\tau_{C,1,m-1}, \tau_{A,1,m-1}), \ldots, (\tau_{C,n_{\max},m-1}, \tau_{A,n_{\max},m-1}).$$

We take, for $n = 1, \ldots, n_{\max}$ the $n+1$-th chain to be the adaptive Metropolis-Hasting sampler [3]. Our sampler will asymptotically sample from the posterior distribution conditioned on $n$. We start our sampler in the maximum likelihood estimator for $(\tau_C, \tau_A, \varepsilon)$ estimated over a course grid of the domain of the prior in order to speed up convergence. For $m \geq 2$, suppose we are at state $(\tau_{C,n,m-1}, \tau_{A,n,m-1}, \varepsilon_{n,m-1})$. Then we simulate a proposal for $\tau_C, \tau_A$ and $\varepsilon$, from independent normal distributions with means $\tau_{C,n,m-1}$, $\tau_{A,n,m-1}$, and $\varepsilon_{n,m-1}$, respectively and variances $c\sigma_x$, where $\sigma_x = \hat{\sigma}_x + \delta$, where $x = \tau_C, \tau_A, \varepsilon$, respectively, and $\sigma_x$ is fixed for the first 5'000 samples and then the empirical variance $\hat{\sigma}_x$ is calculated from samples $3000, 3020, 3040$, until sample $\lfloor (m-1)/1000 \rfloor \cdot 1000$ (so updated every 1000 steps). $\delta$ is a small constant to prevent $\sigma_x$ from becoming zero. $c$ is a scaling constant, which is initially set to $2.38^2/3$, as suggested in [3], but then from step 10'000 until at most a quarter of the total number of steps; it is every 1000 steps updated so that the acceptance rate is between 0.1 and 0.15. If the acceptance rate is too low, we set $c \to 3c/4$; if the acceptance rate is too large, we set $c \to 4c/3$.

So, at step $m-1$, for chain $n+1$, a new proposal $(\tau_{C,n,prop}, \tau_{A,n,prop}, \varepsilon_{n,prop})$ is accepted with probability

$$\min\left(\frac{P(d \mid n, \tau_{C,n,prop}, \tau_{A,n,prop}, \varepsilon_{n,prop})\pi(\tau_{C,n,prop}, \tau_{A,n,prop})\pi(\varepsilon_{n,prop})}{P(d \mid n, \tau_{C,n,m-1}, \tau_{A,n,m-1}, \varepsilon_{n,m-1})\pi(\tau_{C,n,m-1}, \tau_{A,n,m-1})\pi(\varepsilon_{n,m-1})}, 1\right).$$

It is possible that some proposals for some of the $n_{\max}$ chains get accepted, and some might not be accepted in the same grand step $m$.

After sampling the $n_{\max}$ conditional probabilities, we sample $n$ from the proposal function, which is $n = n_{m-1}$ with probability 0.8 or one of the other values $1, \ldots, n_{m-1}-1, n_{m-1}+1, \ldots, n_{\max}$ with probability $0.2/(n_{\max}-1)$ each. When $n_{\text{prop}} \neq n_{m-1}$, we set $n_m = n_{\text{prop}}$ with probability

$$\min\left(\frac{P(d \mid n_{\text{prop}}, \tau_{C,n_{\text{prop}},m}, \tau_{A,n_{\text{prop}},m}, \varepsilon_{n_{\text{prop}},m})\pi(n_{\text{prop}})\pi(\tau_{C,n_{\text{prop}},m}, \tau_{A,n_{\text{prop}},m})\pi(\varepsilon_{n_{\text{prop}},m})}{P(d \mid n_{m-1}, \tau_{C,n_{m-1},m}, \tau_{A,n_{m-1},m}, \varepsilon_{n_{m-1},m})\pi(n_{m-1})\pi(\tau_{C,n_{m-1},m}, \tau_{A,n_{m-1},m})\pi(\varepsilon_{n_{m-1},m})}, 1\right),$$

otherwise, we set we set $n_m$ equal to $n_{m-1}$.

Next we take the $\tau_C$ and $\tau_A$ value belonging to the $n_m+1$-th chain, so we take $(n_m, \tau_{C,n_m,m}, \tau_{A,n_m,m})$ as our posterior sample, which, according to Theorem S5, converges to the posterior distribution.

### S3 Some identifiability issues

#### S3.1 Deamination and misread errors are inseparable

Suppose at death, at a specific locus, there are $k$ derived alleles and $2n-k$ ancestral alleles. At the time of sampling, some of the derived alleles might deaminate into ancestral alleles, each allele with probability $\varepsilon_{\text{deam}}$, and an allele ancestral at death might deaminate into a derived allele with probability $\varepsilon_{\text{deam}}$. Assume that the sample has $k'$ derived alleles, some of which might be deaminated.

The probability of a derived read, given $k'$ derived alleles in the sample, is the probability of choosing a derived allele and having no misread or choosing an ancestral allele and having a misread. The probability of these events is

$$(1-\varepsilon_{\text{misr}})\frac{k'}{2n}+\varepsilon_{\text{misr}}\left(1-\frac{k'}{2n}\right).$$

Let $P(k' \mid k)$ denote the probability of having $k'$ derived alleles in the sample, given $k$ derived alleles at death.

So, the probability of a derived read given $k$ derived alleles at death is

$$\begin{aligned}
&\sum_{k'=0}^{2n}\left((1-\varepsilon_{\text{misr}})\frac{k'}{2n}+\varepsilon_{\text{misr}}\left(1-\frac{k'}{2n}\right)\right)P(k' \mid k)\\
=&(1-\varepsilon_{\text{misr}})\sum_{k'=0}^{2n}\frac{k'}{2n}P(k' \mid k)+\varepsilon_{\text{misr}}\left(1-\sum_{k'=0}^{2n}\frac{k'}{2n}P(k' \mid k)\right).
\end{aligned}$$

So now we need to know the expectation of $k'$ given $k$. Note that there are $k$ derived alleles at death, whom each independently remained derived with probability $1-\varepsilon_{\text{deam}}$ and $2n-k$ ancestral alleles at death, whom each independently deaminated with probability $\varepsilon_{\text{deam}}$. So the expectation of $k'$ is

$$(1-\varepsilon_{\text{deam}})k+\varepsilon_{\text{deam}}(2n-k).$$

So, the probability of a derived read is

$$\begin{aligned}
&(1-\varepsilon_{\text{misr}})\frac{(1-\varepsilon_{\text{deam}})k+\varepsilon_{\text{deam}}(2n-k)}{2n}+\varepsilon_{\text{misr}}\left(1-\frac{(1-\varepsilon_{\text{deam}})k+\varepsilon_{\text{deam}}(2n-k)}{2n}\right)\\
=&(1-\varepsilon_{\text{misr}})(1-\varepsilon_{\text{deam}})\frac{k}{2n}+(1-\varepsilon_{\text{misr}})\varepsilon_{\text{deam}}\frac{2n-k}{2n}\\
&+\varepsilon_{\text{misr}}\left(\frac{k+2n-k}{2n}-\frac{(1-\varepsilon_{\text{deam}})k+\varepsilon_{\text{deam}}(2n-k)}{2n}\right)\\
=&(1-\varepsilon_{\text{misr}})(1-\varepsilon_{\text{deam}})\frac{k}{2n}+(1-\varepsilon_{\text{misr}})\varepsilon_{\text{deam}}\frac{2n-k}{2n}\\
&+\varepsilon_{\text{misr}}\left(\varepsilon_{\text{deam}}\frac{k}{2n}+(1-\varepsilon_{\text{deam}})\frac{2n-k}{2n}\right)\\
=&(1-\varepsilon_{\text{misr}}-\varepsilon_{\text{deam}}+2\varepsilon_{\text{misr}}\varepsilon_{\text{deam}})\frac{k}{2n}+(\varepsilon_{\text{misr}}+\varepsilon_{\text{deam}}-2\varepsilon_{\text{misr}}\varepsilon_{\text{deam}})\frac{2n-k}{2n}.
\end{aligned}$$

Now, let $\varepsilon=\varepsilon(\varepsilon_{\text{misr}},\varepsilon_{\text{deam}})=\varepsilon_{\text{misr}}+\varepsilon_{\text{deam}}-2\varepsilon_{\text{misr}}\varepsilon_{\text{deam}}$, then the probability of a derived allele is

$$(1-\varepsilon)\frac{k}{2n}+\varepsilon\frac{2n-k}{2n}.$$

So the number of derived reads depends only on the errors via $\varepsilon=\varepsilon_{\text{misr}}+\varepsilon_{\text{deam}}-2\varepsilon_{\text{misr}}\varepsilon_{\text{deam}}$. Under this model, it is impossible to estimate $\varepsilon_{\text{misr}}$ and $\varepsilon_{\text{deam}}$ separately, only the quantity $\varepsilon_{\text{misr}}+\varepsilon_{\text{deam}}-2\varepsilon_{\text{misr}}\varepsilon_{\text{deam}}$ at once. Several different values of $\varepsilon_{\text{deam}}$ and $\varepsilon_{\text{misr}}$ might lead to the same value of $\varepsilon$, for example $\varepsilon_{\text{deam}}=0$ and $\varepsilon_{\text{misr}}=\varepsilon$ or $\varepsilon_{\text{deam}}=\varepsilon$ and $\varepsilon_{\text{misr}}=0$. [1] confirmed this with simulations. They found no benefit of using separate parameters for the deamination and read error. Note that for small $\varepsilon_{\text{deam}}$ and $\varepsilon_{\text{misr}}$, $\varepsilon\approx\varepsilon_{\text{misr}}+\varepsilon_{\text{deam}}$, so, one can think of $\varepsilon$ as the sum of the two errors. Simple algebra shows that $\varepsilon(\varepsilon_{\text{misr}},\varepsilon_{\text{deam}})$ takes values in $[0,1]$ for all $\varepsilon_{\text{misr}},\varepsilon_{\text{deam}}\in[0,1]$, moreover $\varepsilon(\varepsilon_{\text{misr}},\varepsilon_{\text{deam}})=\varepsilon(1-\varepsilon_{\text{misr}},1-\varepsilon_{\text{deam}})$.

#### S3.2 Identifiability of the error rate

We have to make a point about identifiability. The probability of $d$ derived reads with coverage $R$ and $k$ derived reads and error rate $\varepsilon$ is given by

$$P(d \mid k, \varepsilon, R) = \binom{R}{d}\left((1-\varepsilon)\cdot\frac{k}{2n} + \varepsilon\cdot\frac{2n-k}{2n}\right)^{d}\left(1-(1-\varepsilon)\cdot\frac{k}{2n} - \varepsilon\cdot\frac{2n-k}{2n}\right)^{R-d}.$$

Note that $P(d \mid k, \varepsilon, R)$ satisfies

$$P(d \mid k, \varepsilon, R) = P(d \mid 2n-k, 1-\varepsilon, R),$$

for every $d \in \{0, \ldots, R\}$. So, the likelihood cannot distinguish between $k$ derived alleles with error rate $\varepsilon$ or $2n-k$ derived alleles with error rate $1-\varepsilon$. Therefore, we assume that the error rate is strictly smaller than 0.5, which solves the identifiability issue.

#### S3.3 Mathematical limitations for coverage 1

Schraiber [4] argues in the appendix that loci with coverage one do not contribute information about $\tau_A$.

In addition, it is easy to see that for $R = 1$, the likelihood equals, for $d = 1$,

$$\begin{aligned}
&\sum_{k=0}^{2n}\left((1-\varepsilon)\frac{k}{2n} + \varepsilon\frac{2n-k}{2n}\right)\binom{2n}{k}p_{n,k}(\tau_C, \tau_A) \\
=&\frac{1}{2n}\sum_{k=0}^{2n}((1-\varepsilon)k + \varepsilon(2n-k))\binom{2n}{k}\int_0^1 x^k(1-x)^{2n-k}f(x \mid y, \tau_C, \tau_A)dx \\
=&\frac{1}{2n}\int_0^1\sum_{k=0}^{2n}((1-\varepsilon)k + \varepsilon(2n-k))\binom{2n}{k}x^k(1-x)^{2n-k}f(x \mid y, \tau_C, \tau_A)dx \\
=&\frac{1}{2n}\int_0^1(2(1-\varepsilon)nx + 2\varepsilon n(1-x))f(x \mid y, \tau_C, \tau_A)dx \\
=&\int_0^1((1-\varepsilon)x + \varepsilon(1-x))f(x \mid y, \tau_C, \tau_A)dx,
\end{aligned}$$

which does not depend on $n$. Clearly, for $d = 0$,

$$P(0 \mid 1, n, \tau_C, \tau_A) = 1 - \int_0^1 xf(x)dx$$

does also not depend on $n$. So loci with coverage one do not contribute information about the number of individuals $n$.

### S4 Identifiers for the 1000 Genomes individuals

These are the identifiers used for the synthetic empirical data are found in Supplementary Table 1.

| **Sample IDs from the 1000G** |
| --- |
| HG01680 |
| HG01504 |
| HG02219 |
| HG01777 |
| HG01710 |
| HG02223 |
| HG01607 |
| HG01679 |

Table 1: List of Sample IDs

### S5 Results

#### S5.1 Simple 2-pop simulations

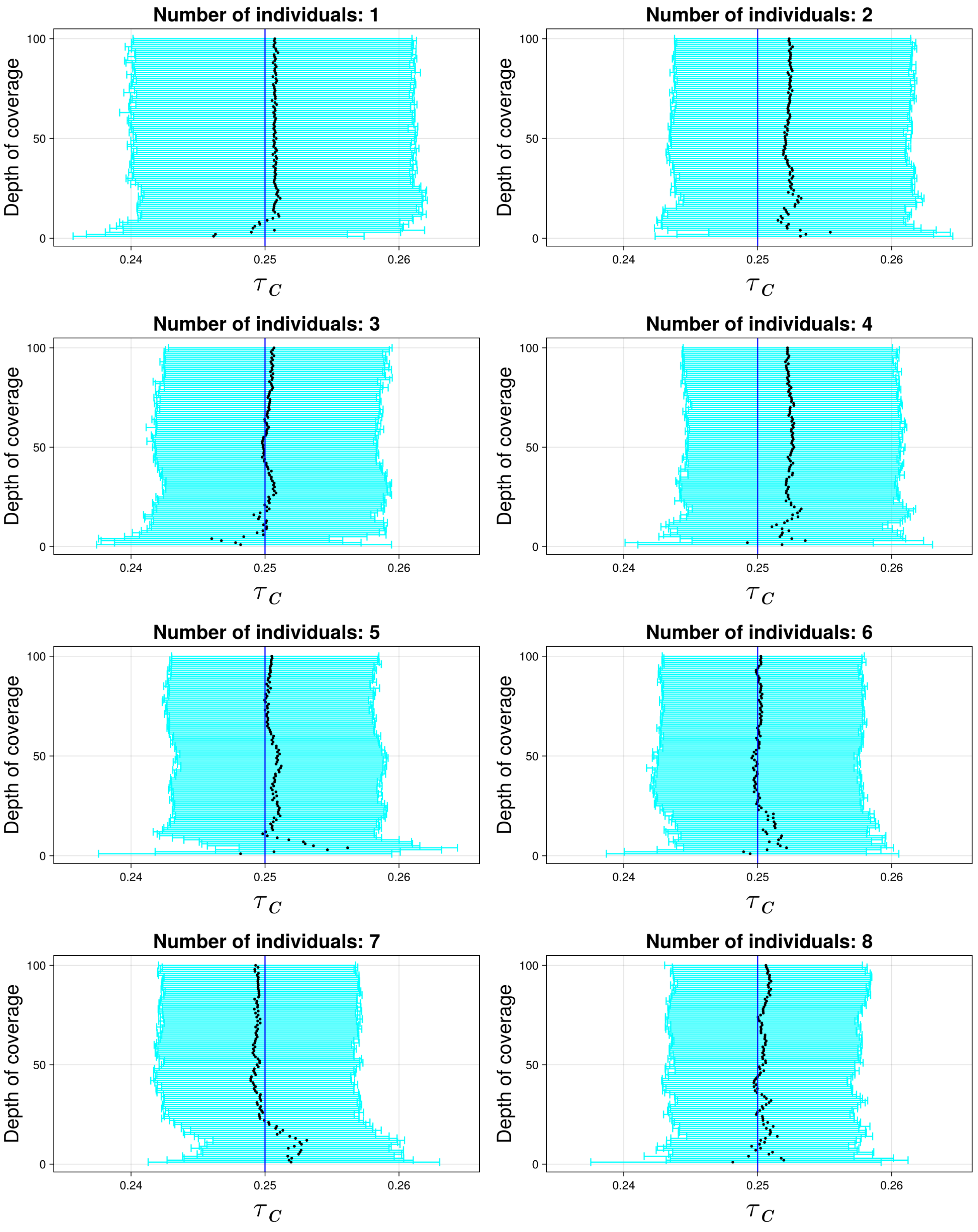

Figure S2: Credible intervals for $\tau_C$, coverages 1,...,100. The blue line indicates the true simulated value. The black dot indicates the median.

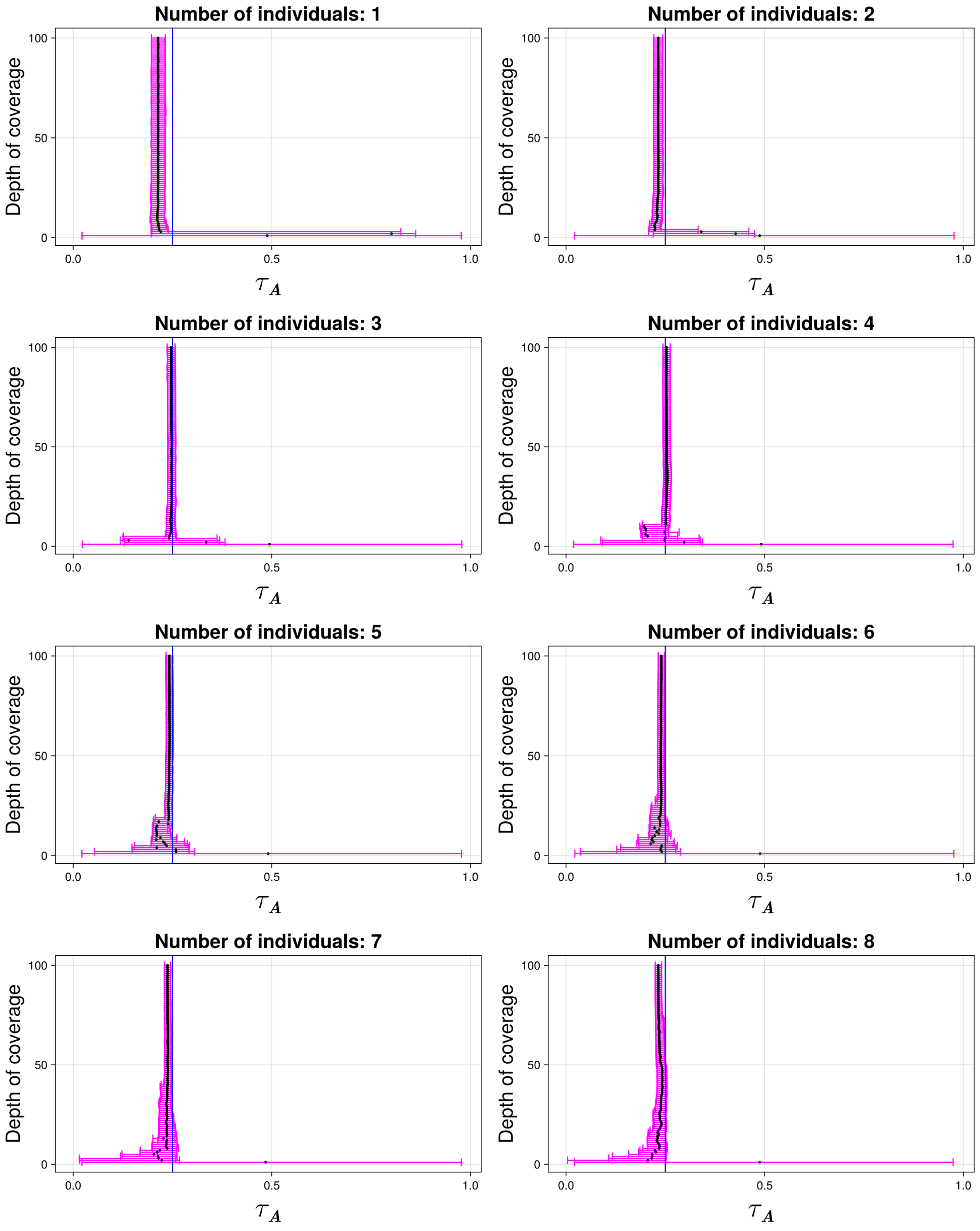

Figure S3: Credible intervals for $\tau_A$, coverages 1,...,100. The blue line indicates the true simulated value. The black dot indicates the median.

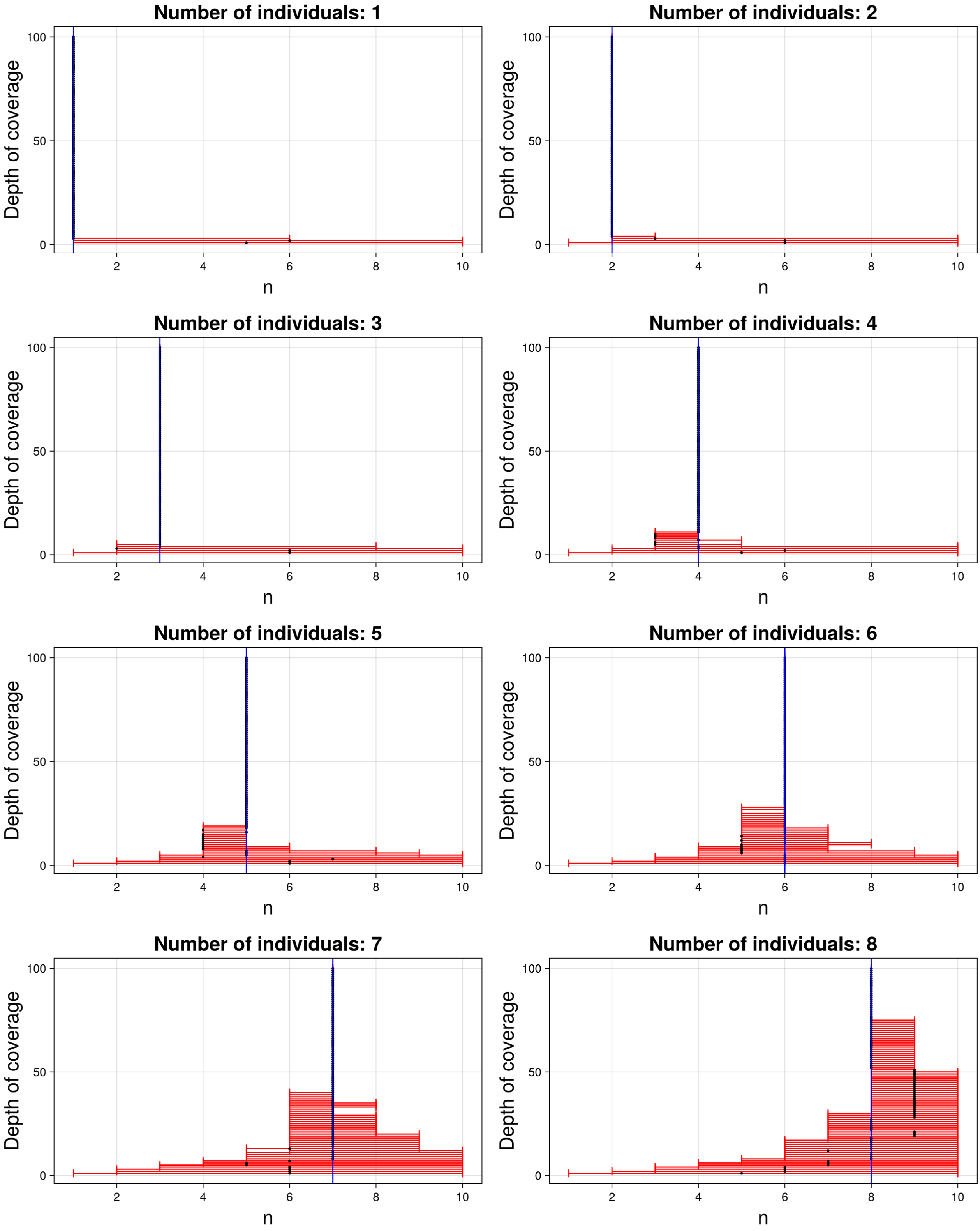

Figure S4: Credible intervals for $n$, coverages 1,...,100. The blue line indicates the true simulated value. The black dot indicates the median. For larger depth of coverage, the credible set consist of only one point.

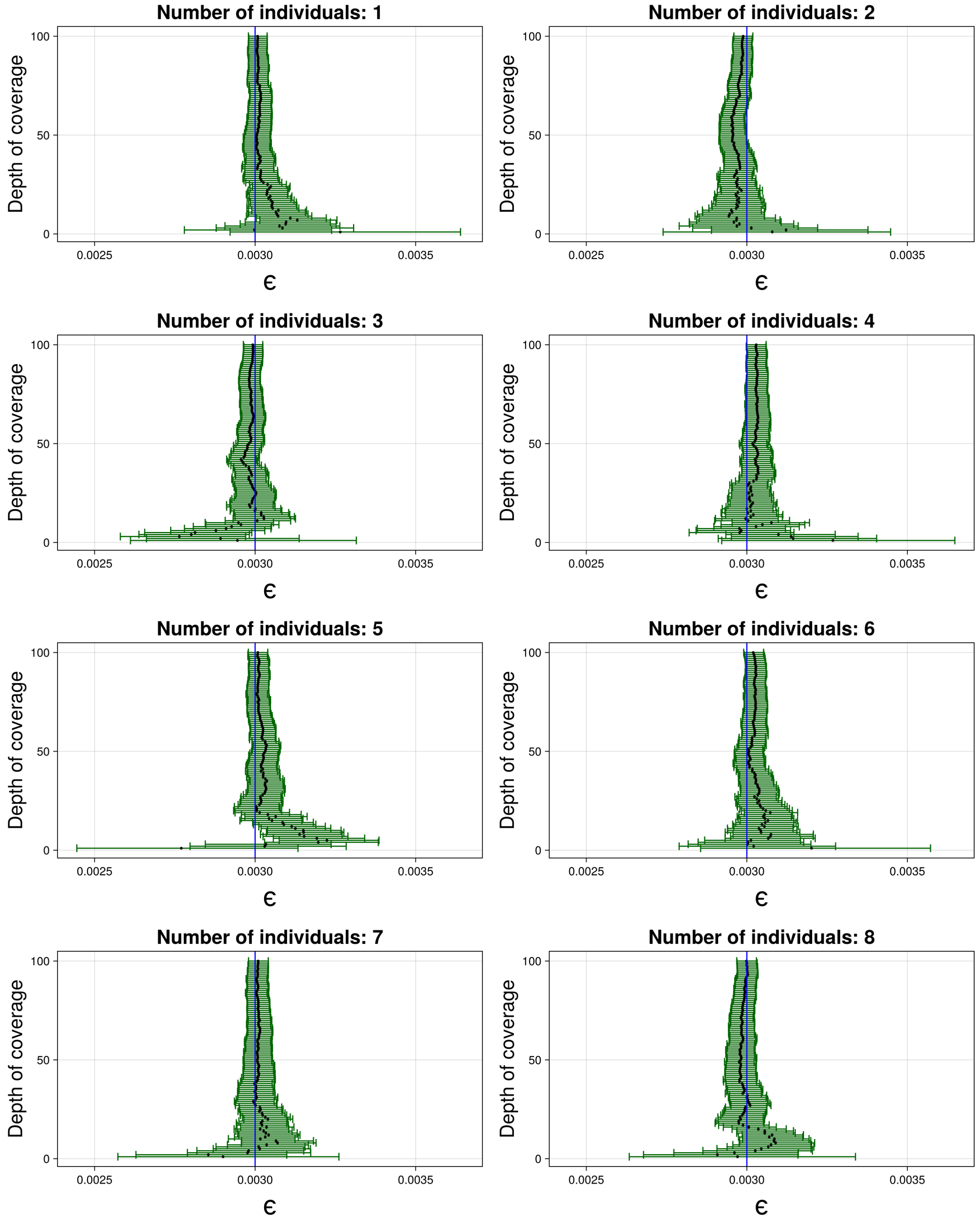

Figure S5: Credible intervals for $\varepsilon$, coverages 1,...,100. The blue line indicates the true simulated value. The black dot indicates the median.

#### S5.2 Plots

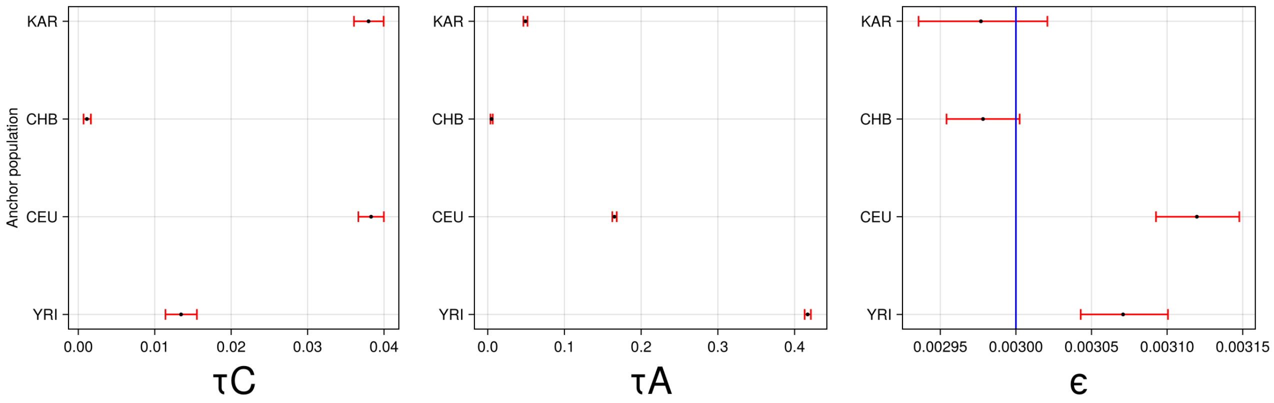

Figure S6: Population of the sample: CHBS. Credible intervals for the drift in the branch leading to the sample $\tau_C$ (left) and drift in the branch of the anchor population $\tau_A$ (middle), with $n = 3$ individuals and a coverage of 20X. To the right is the credible interval for $\varepsilon$. The blue line is the simulated true error level.

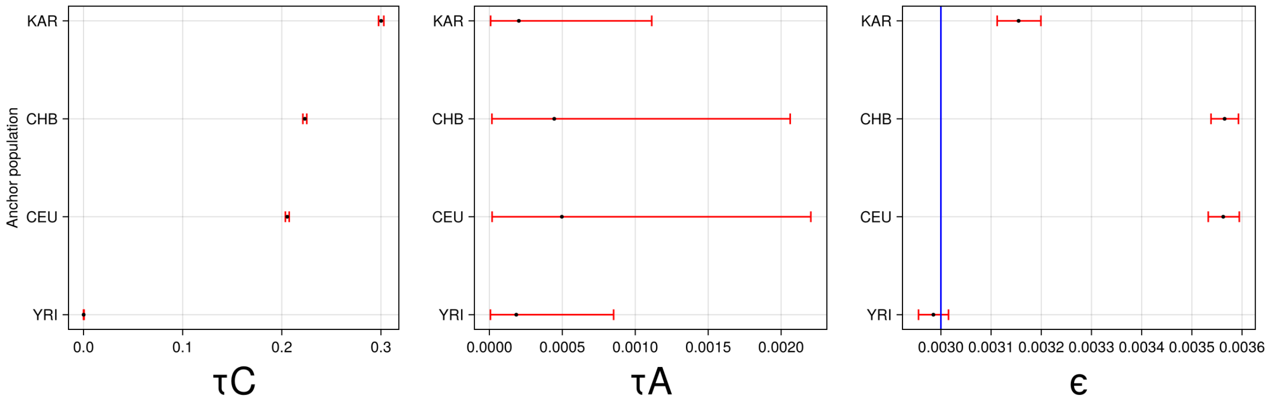

Figure S7: Population of the sample: YRIS. Credible intervals for the drift in the branch leading to the sample $\tau_C$ (left) and drift in the branch of the anchor population $\tau_A$ (middle), with $n = 3$ individuals and a coverage of 20X. To the right is the credible interval for $\varepsilon$. The blue line is the simulated true error level.

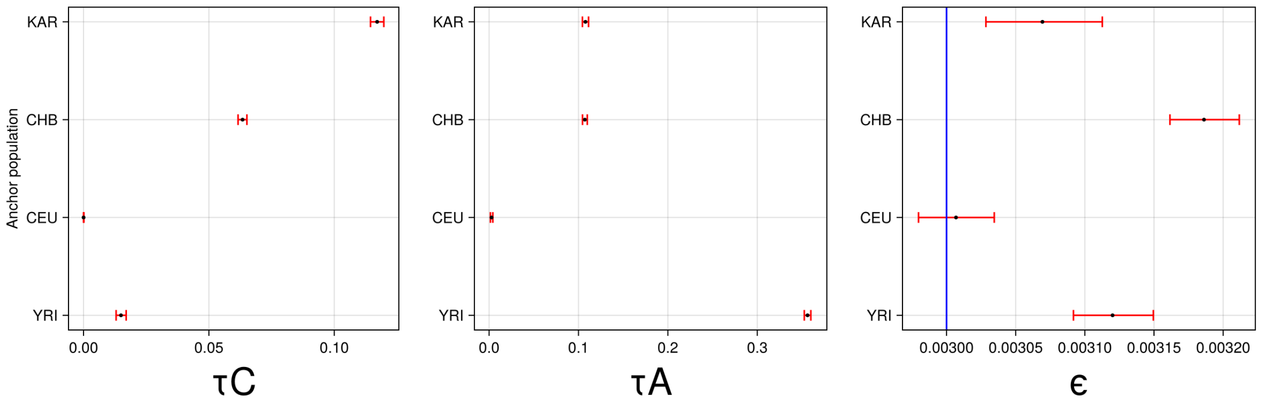

Figure S8: Population of the sample: CEUS. Credible intervals for the drift in the branch leading to the sample $\tau_C$ (left) and drift in the branch of the anchor population $\tau_A$ (middle), with $n = 3$ individuals and a coverage of 20X. To the right is the credible interval for $\varepsilon$. The blue line is the simulated true error level.

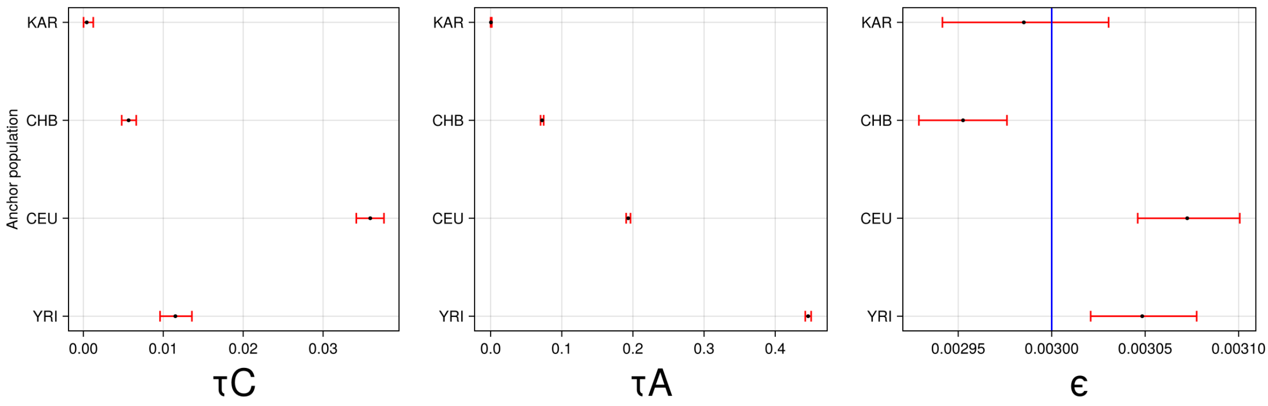

Figure S9: Population of the sample: KARS. Credible intervals for the drift in the branch leading to the sample $\tau_C$ (left) and drift in the branch of the anchor population $\tau_A$ (middle), with $n = 3$ individuals and a coverage of 20X. To the right is the credible interval for $\varepsilon$. The blue line is the simulated true error level.

#### S5.3 Plots for simulated multiple populations

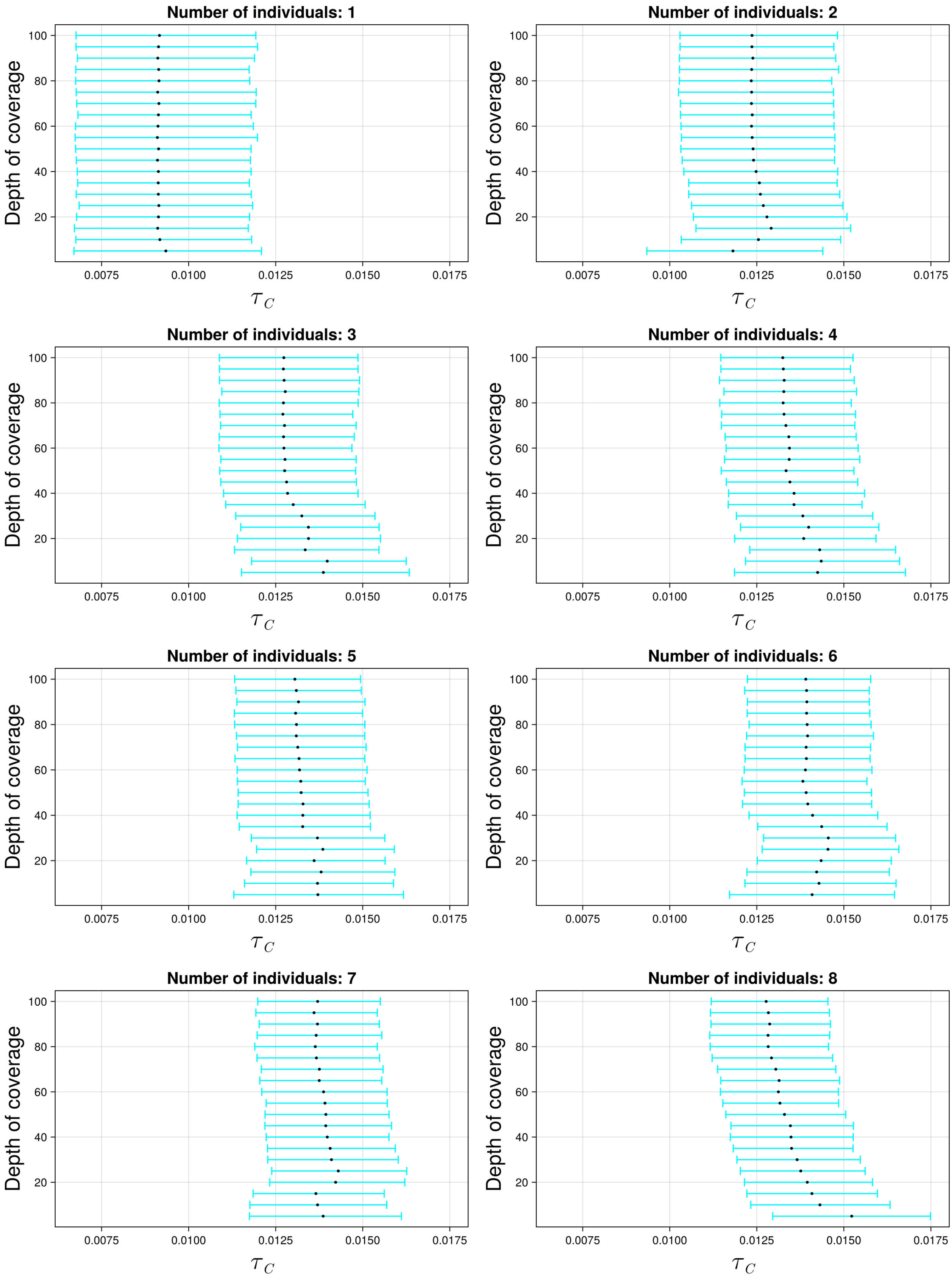

Figure S10: The 95% credible intervals for $\tau_C$ with ancient population CHBS and anchor population YRI. The black dot indicates the median.

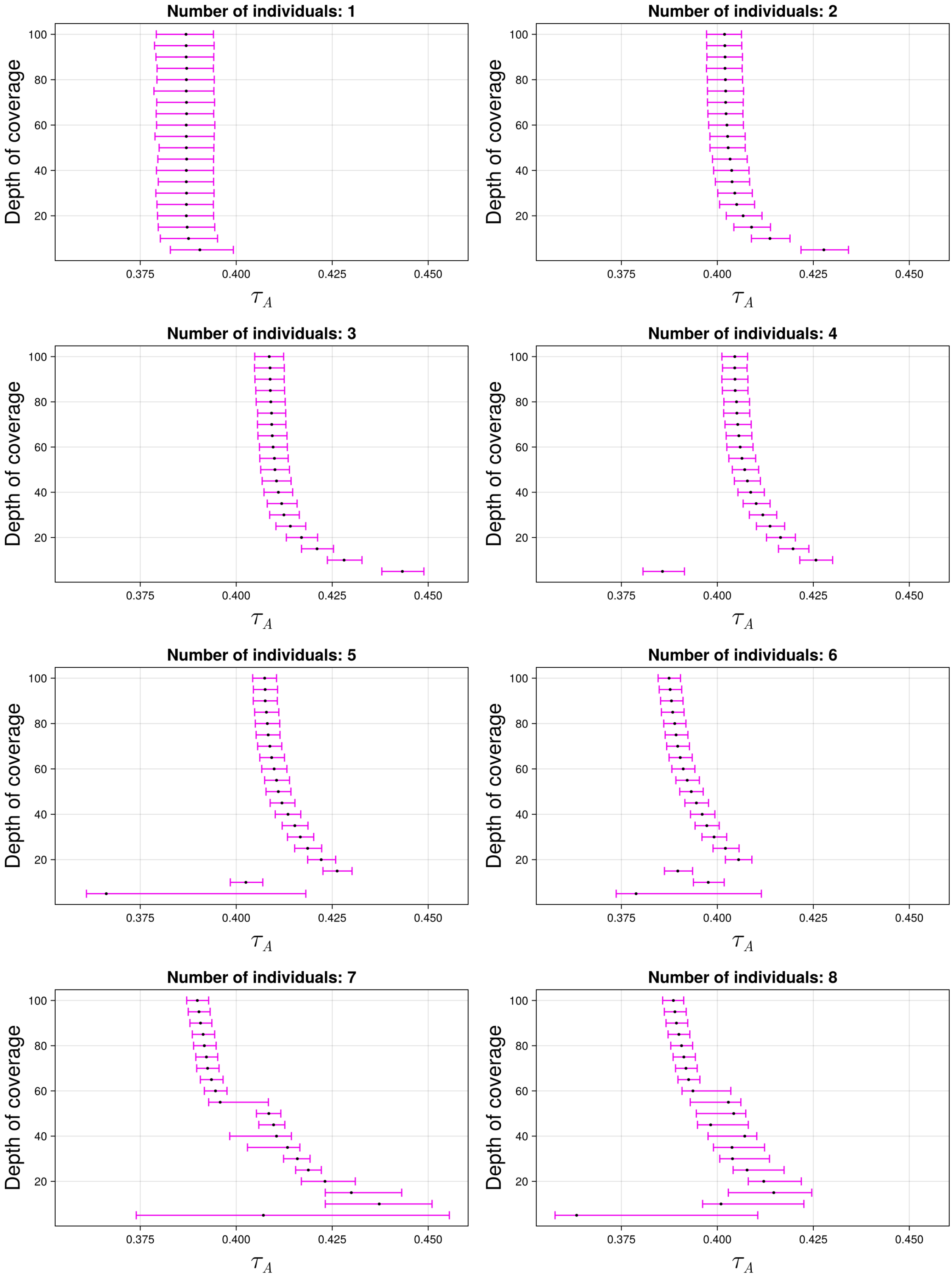

Figure S11: The 95% credible intervals for $\tau_A$ with ancient population CHBS and anchor population YRI. The black dot indicates the median.

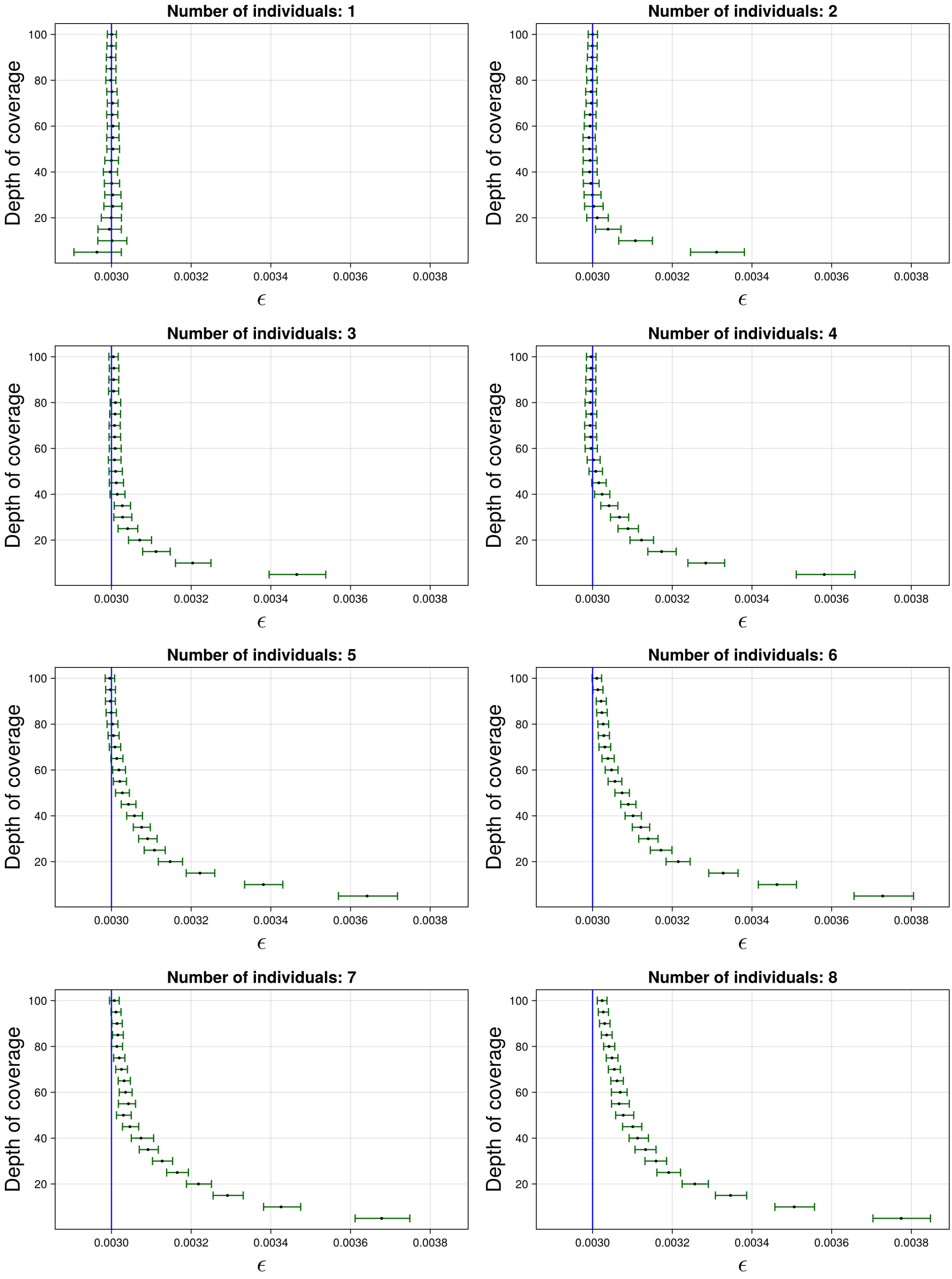

Figure S12: The 95% credible intervals for $\varepsilon$ with ancient population CHBS and anchor population YRI. The black dot indicates the median. The blue line indicates the true simulated value.

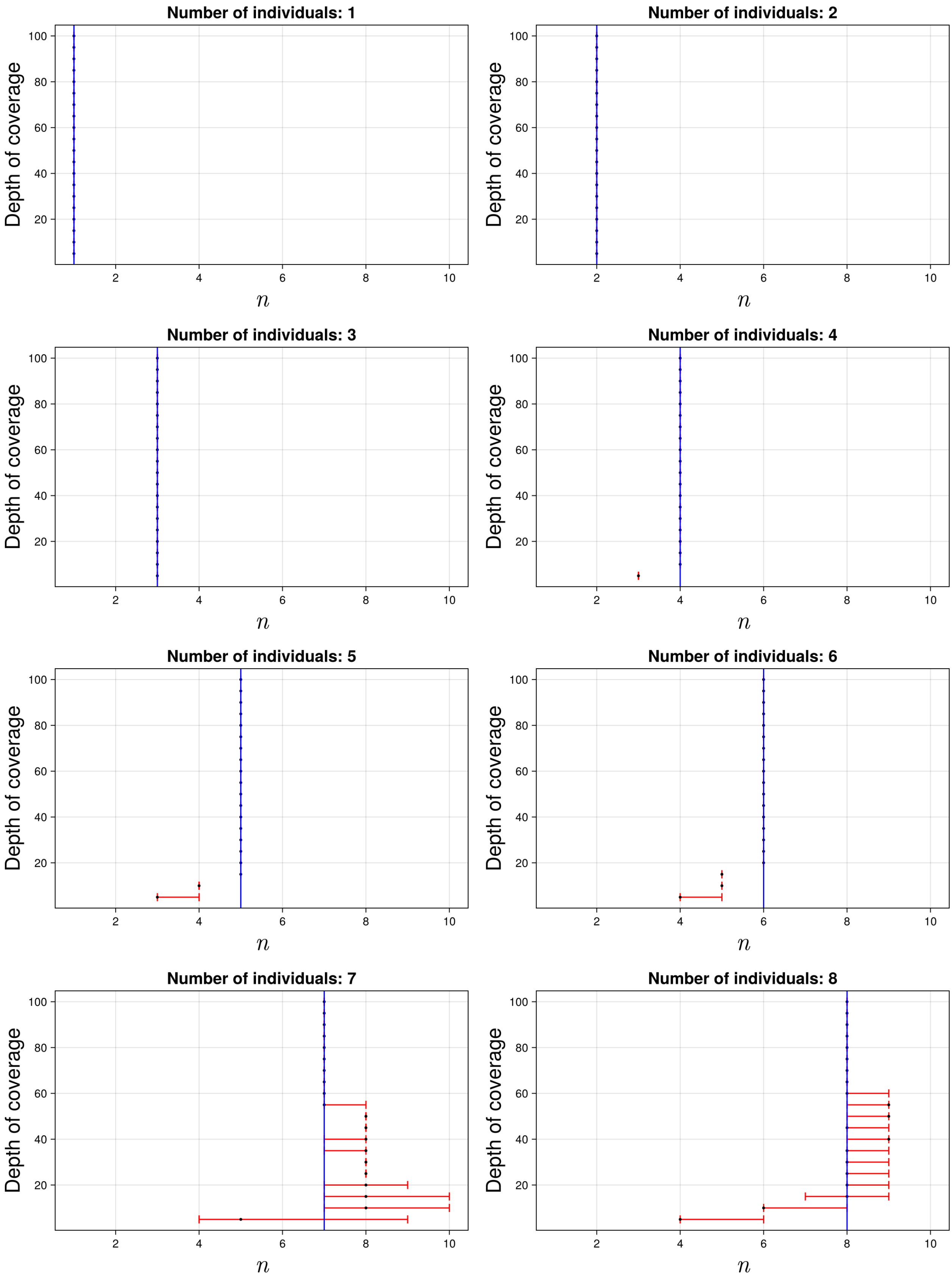

Figure S13: The 95% credible intervals for $n$ with ancient population CHBS and anchor population YRI. The black dot indicates the median. The blue line indicates the true simulated value.

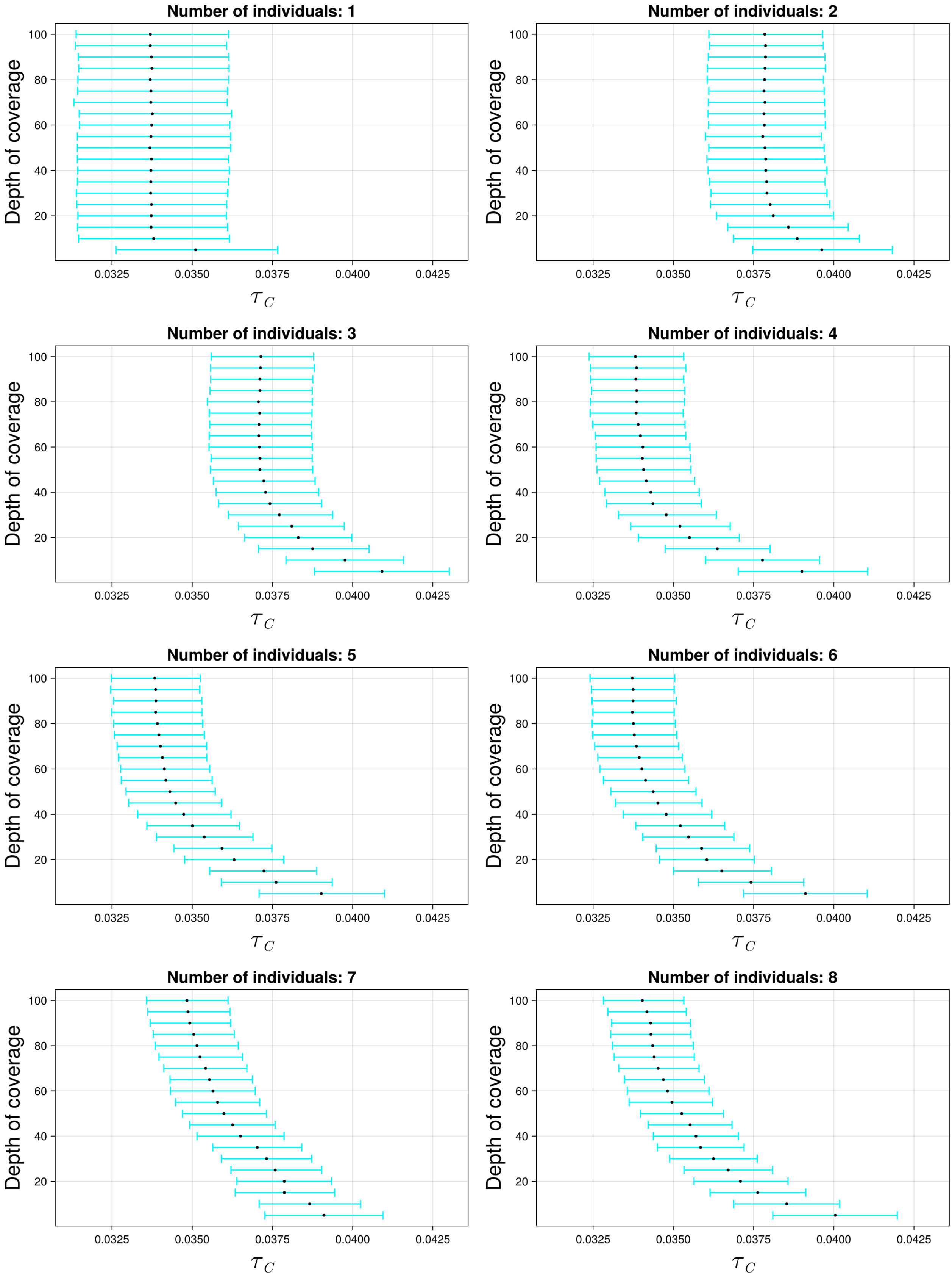

Figure S14: The 95% credible intervals for $\tau_C$ with ancient population CHBS and anchor population CEU. The black dot indicates the median.

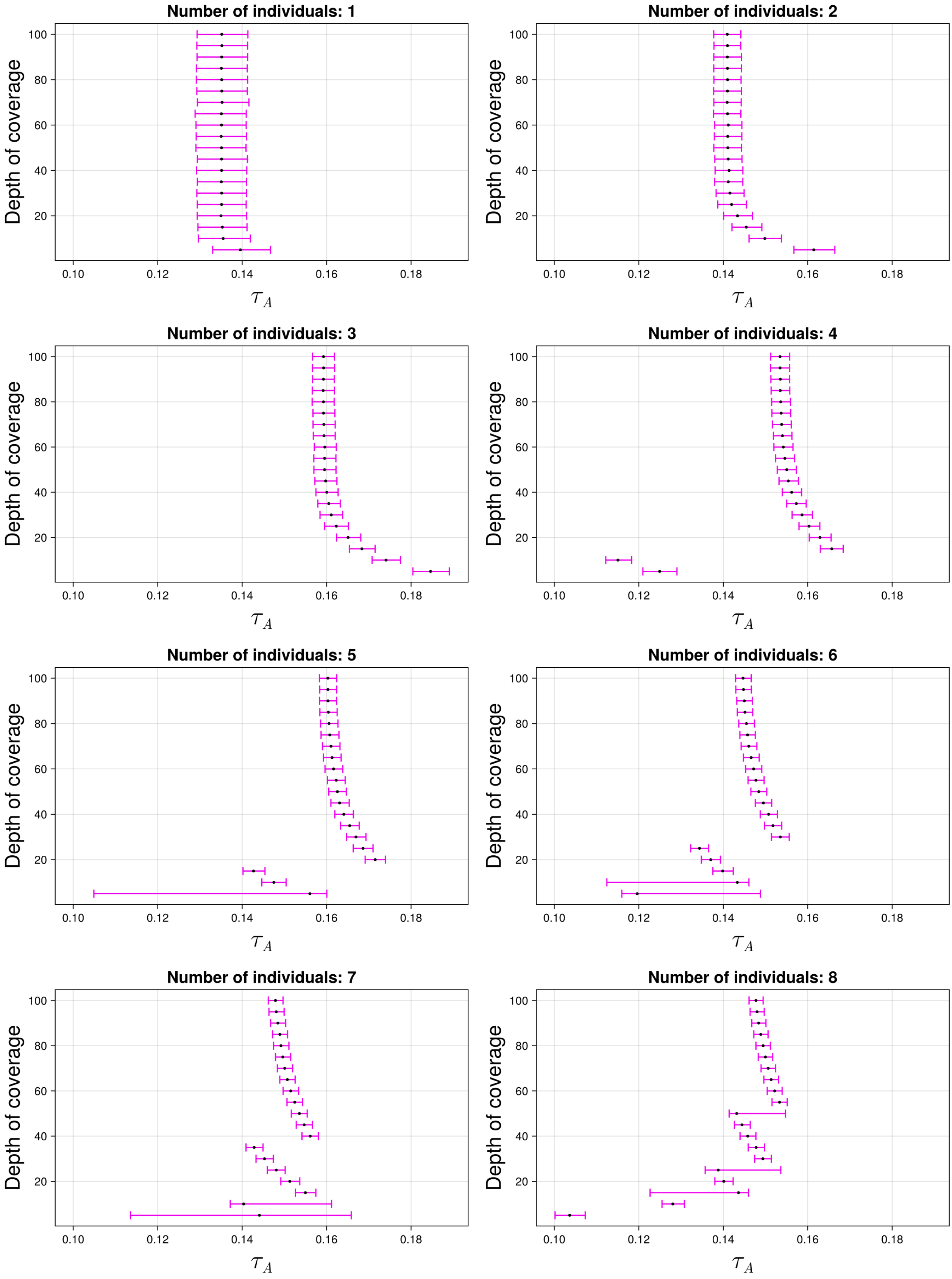

Figure S15: The 95% credible intervals for $\tau_A$ with ancient population CHBS and anchor population CEU. The black dot indicates the median.

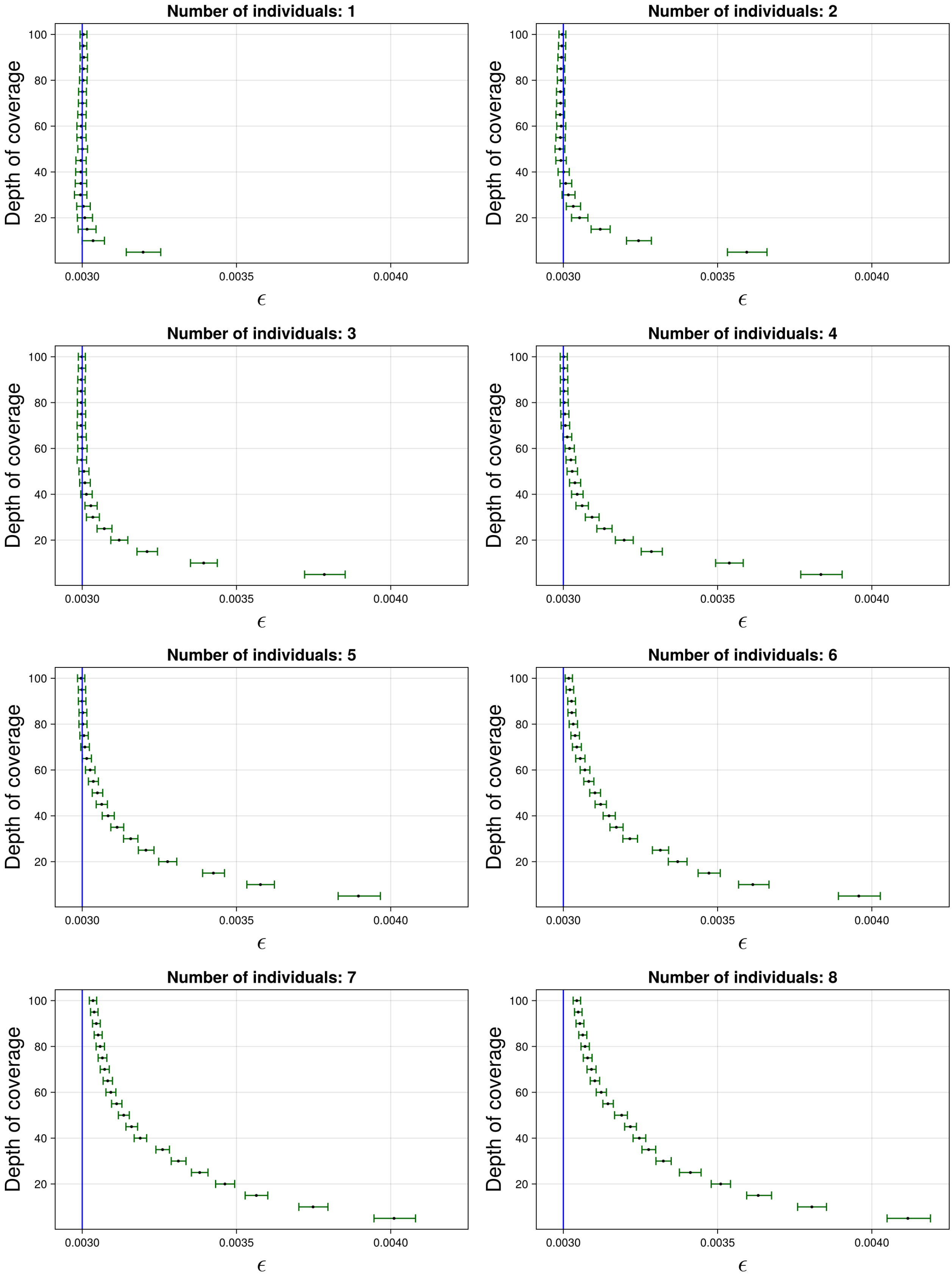

Figure S16: The 95% credible intervals for $\varepsilon$ with ancient population CHBS and anchor population CEU. The black dot indicates the median. The blue line indicates the true simulated value.

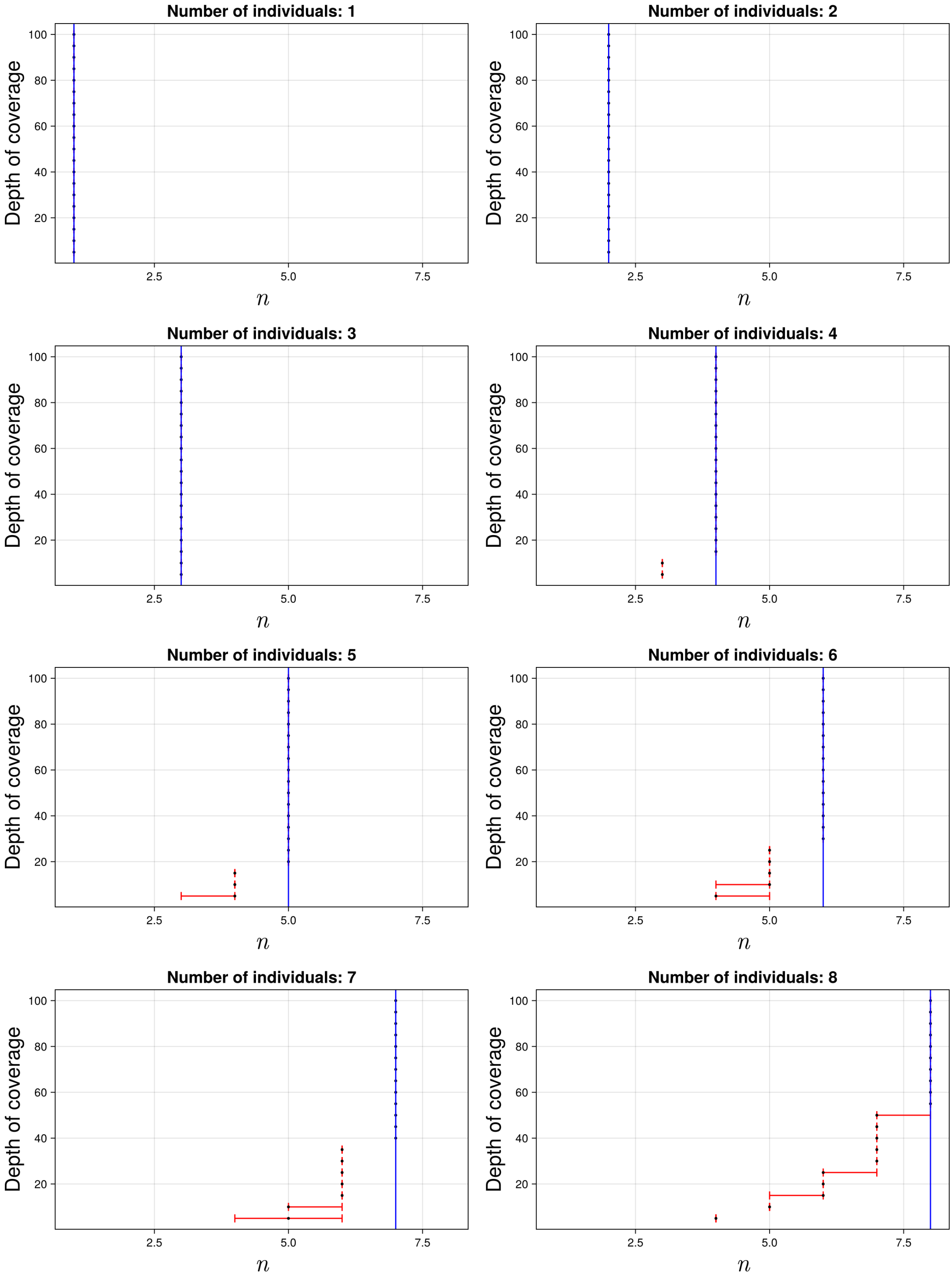

Figure S17: The 95% credible intervals for $n$ with ancient population CHBS and anchor population CEU. The black dot indicates the median. The blue line indicates the true simulated value.

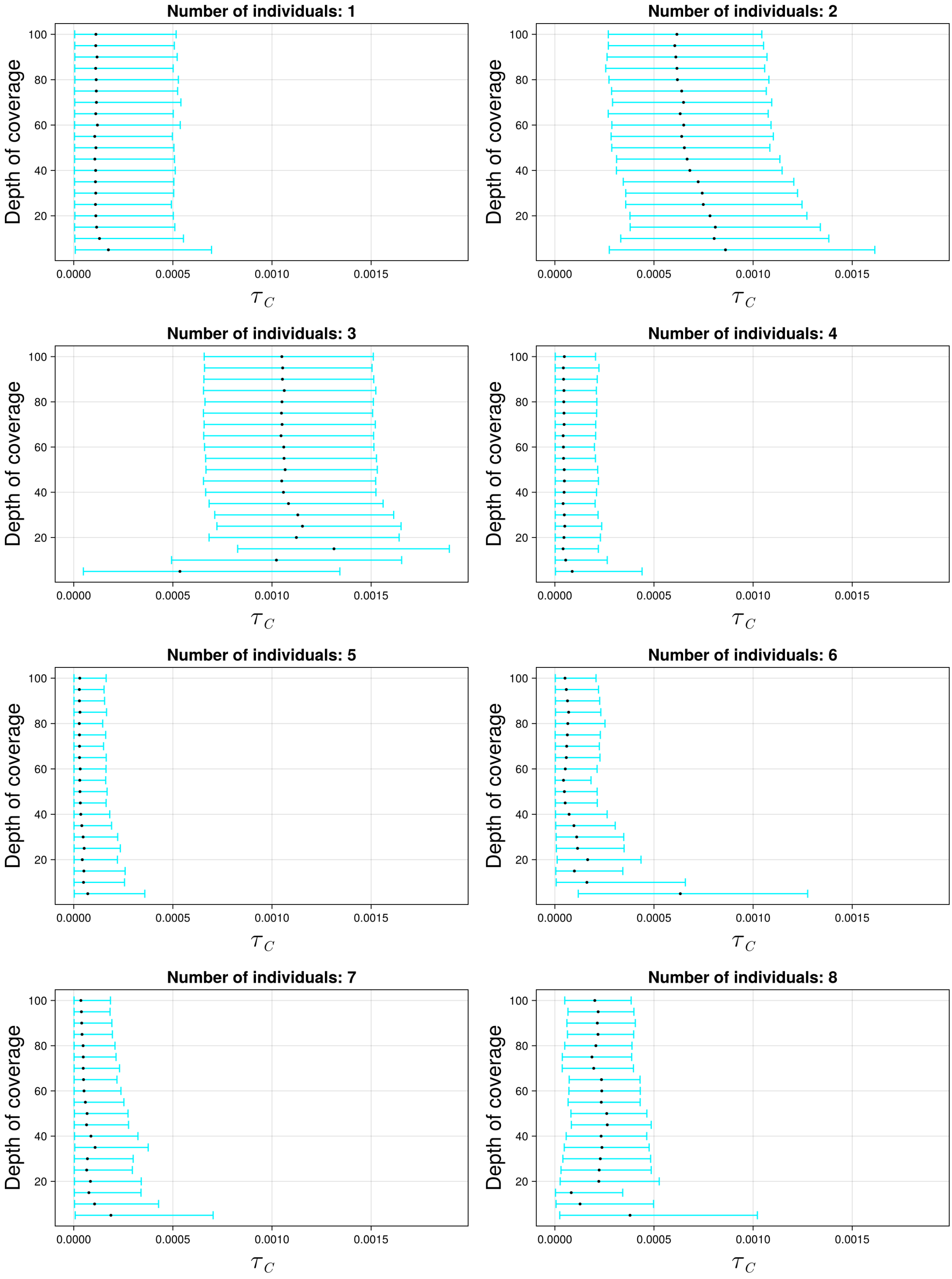

Figure S18: The 95% credible intervals for $\tau_C$ with ancient population CHBS and anchor population CHB. The black dot indicates the median.

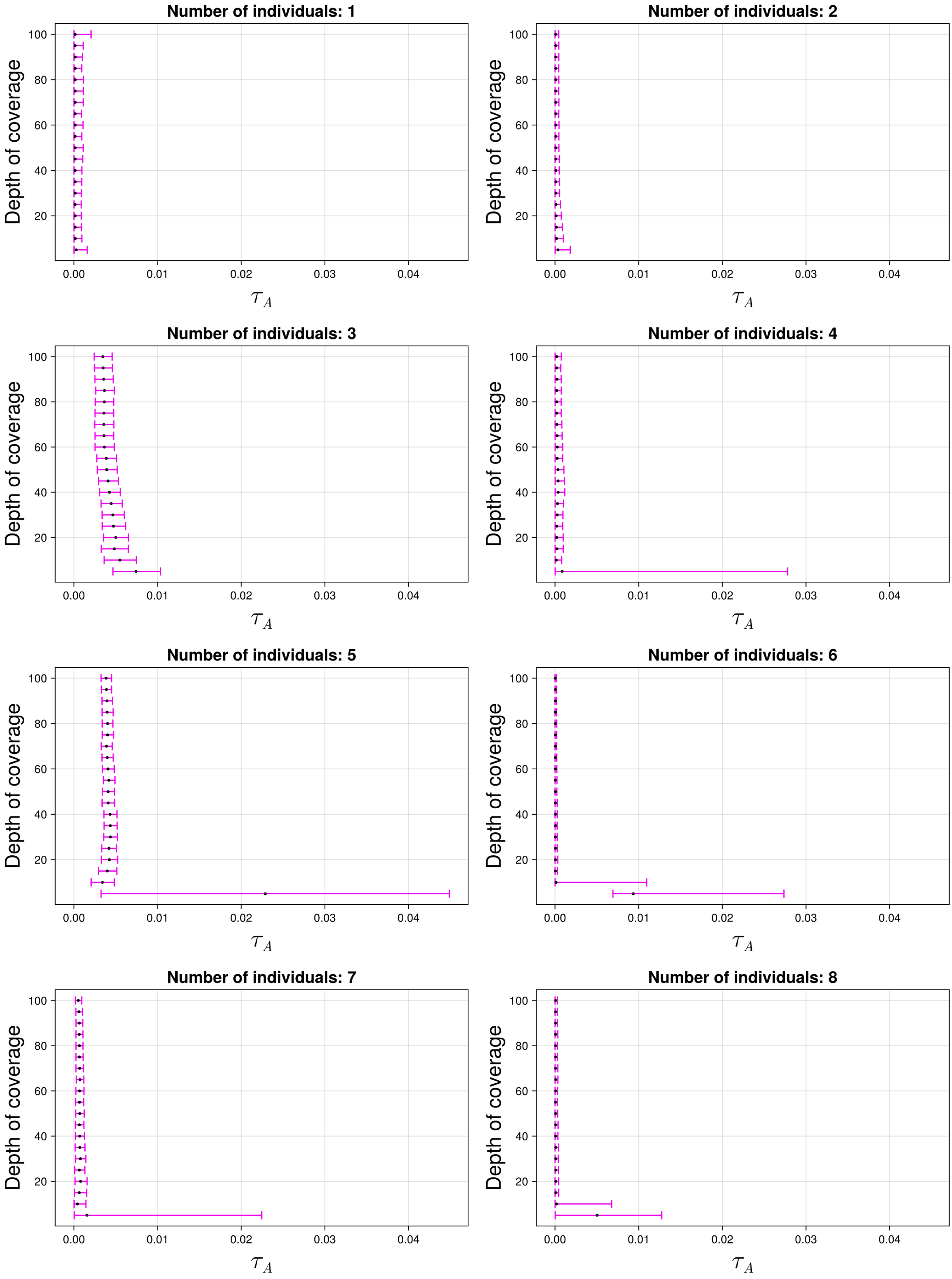

Figure S19: The 95% credible intervals for $\tau_A$ with ancient population CHBS and anchor population CHB. The black dot indicates the median.

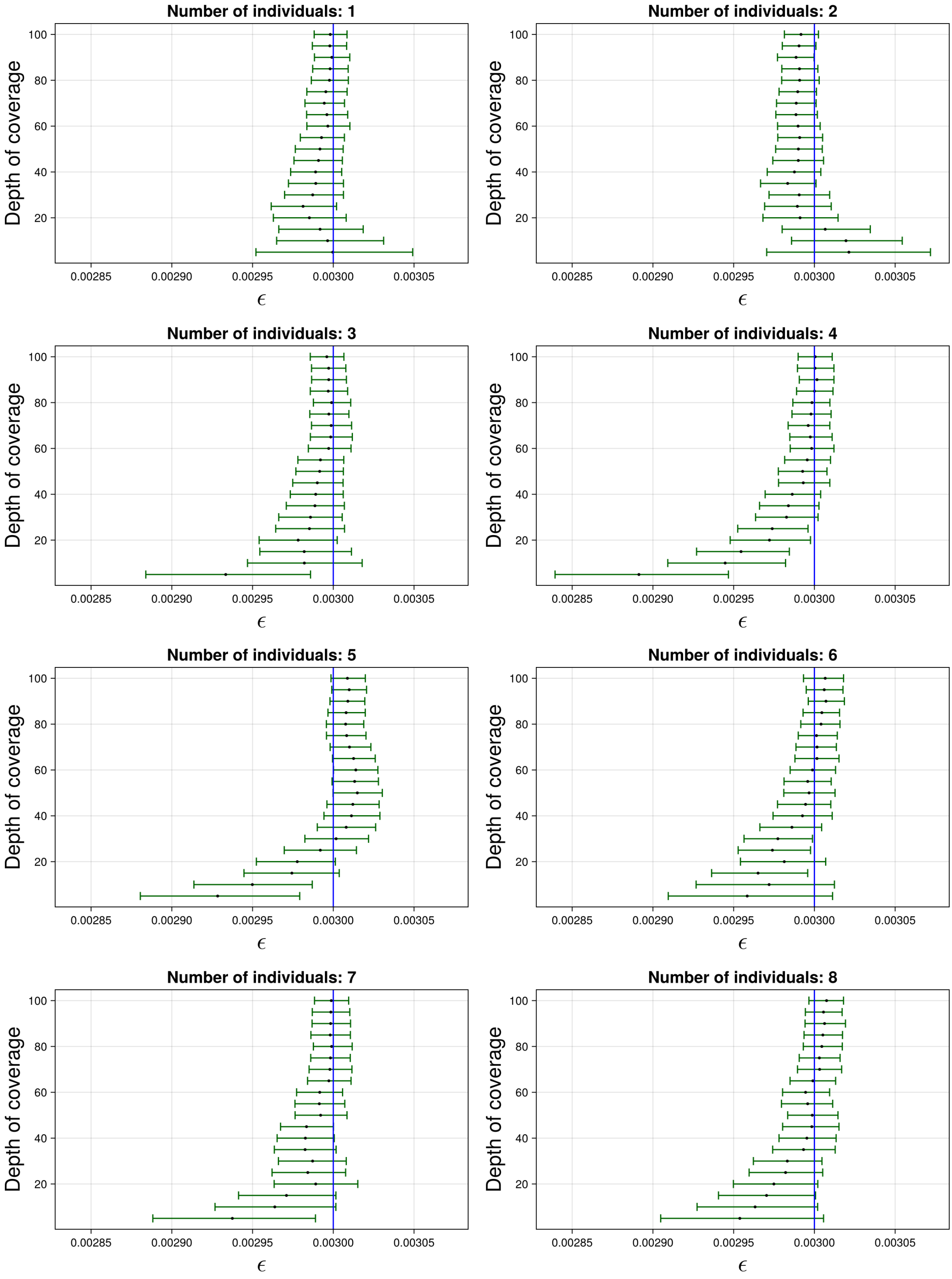

Figure S20: The 95% credible intervals for $\varepsilon$ with ancient population CHBS and anchor population CHB. The black dot indicates the median. The blue line indicates the true simulated value.

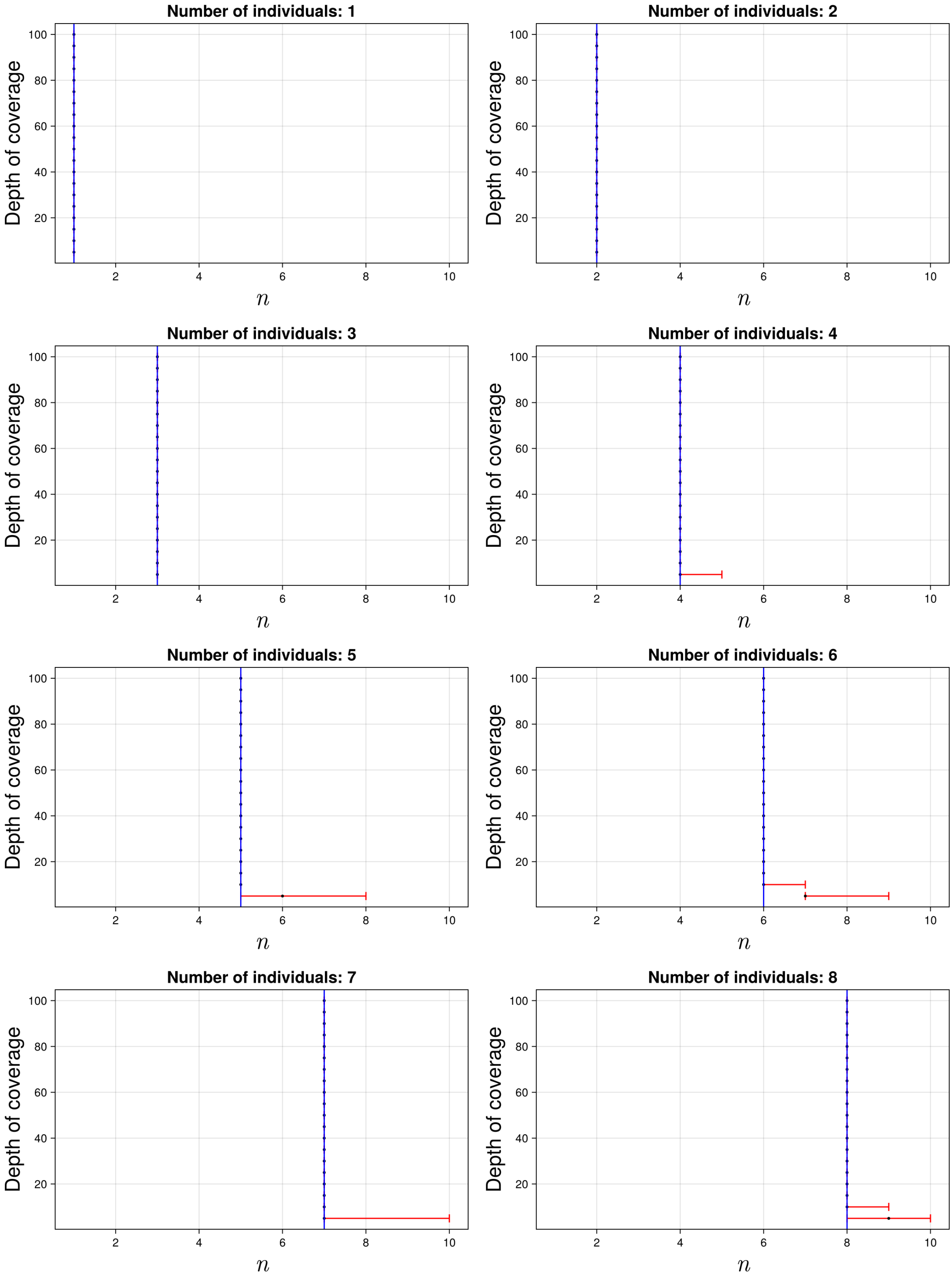

Figure S21: The 95% credible intervals for $n$ with ancient population CHBS and anchor population CHB. The black dot indicates the median. The blue line indicates the true simulated value.

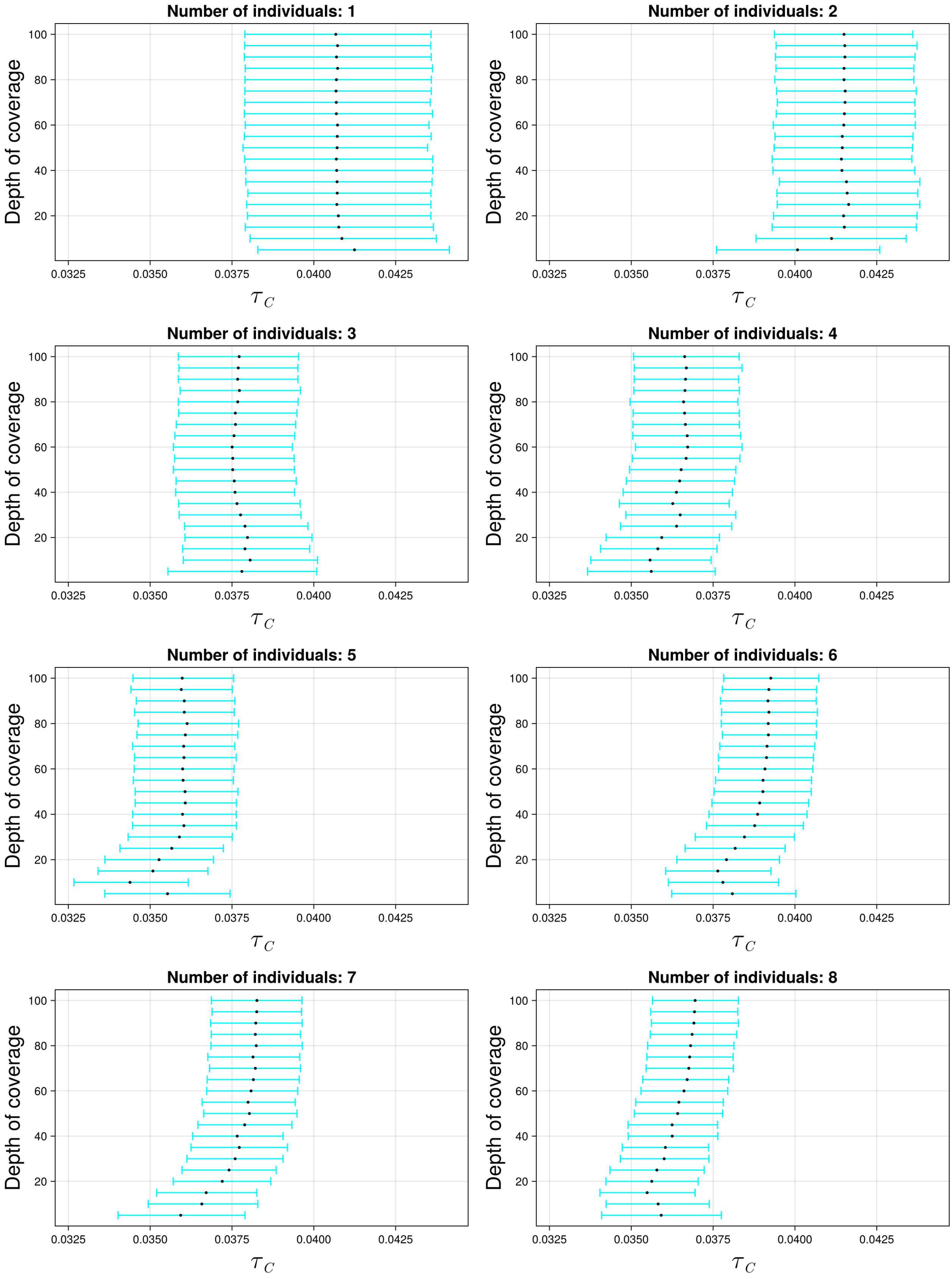

Figure S22: The 95% credible intervals for $\tau_C$ with ancient population CHBS and anchor population KAR. The black dot indicates the median.

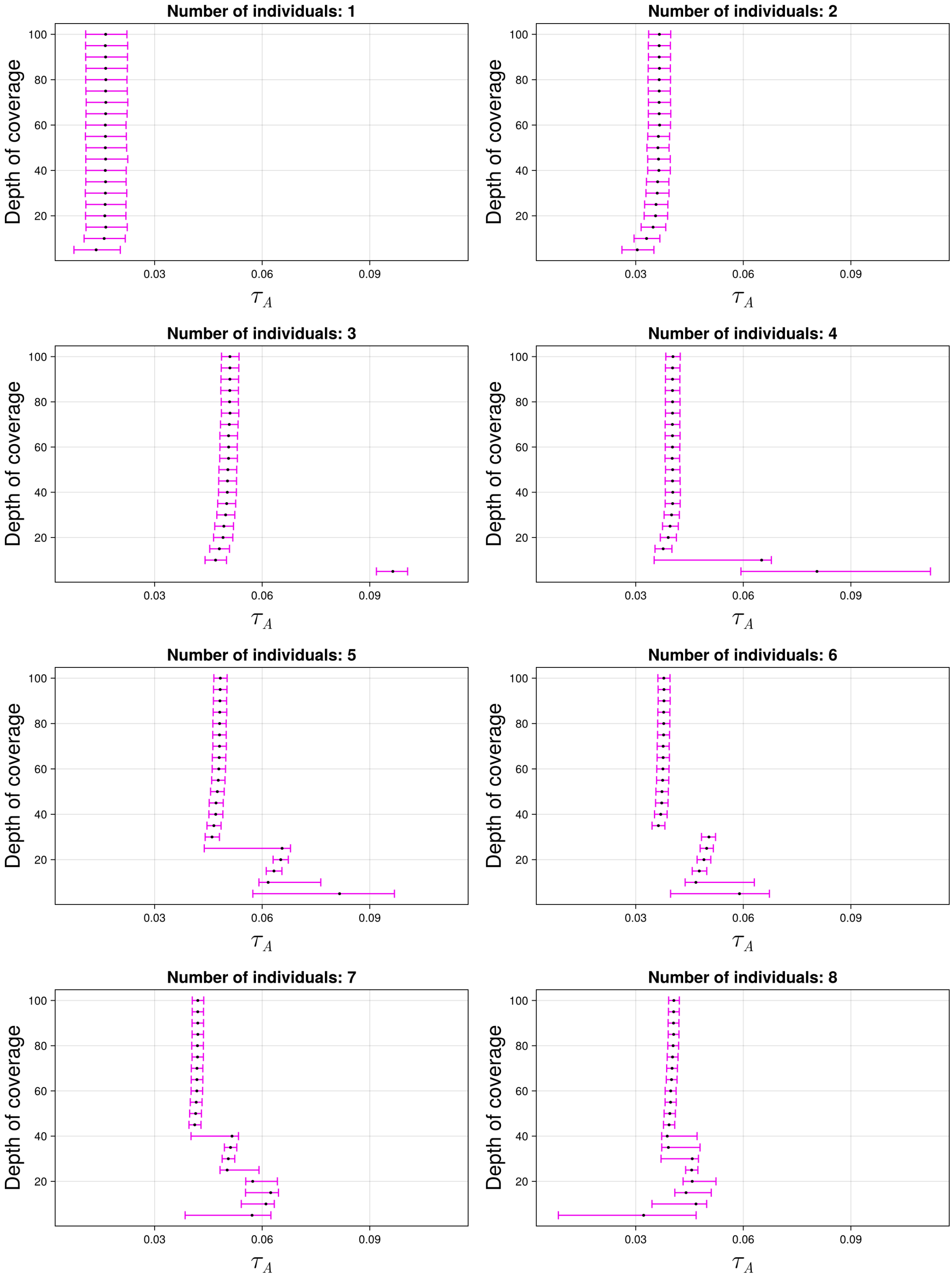

Figure S23: The 95% credible intervals for $\tau_A$ with ancient population CHBS and anchor population KAR. The black dot indicates the median.

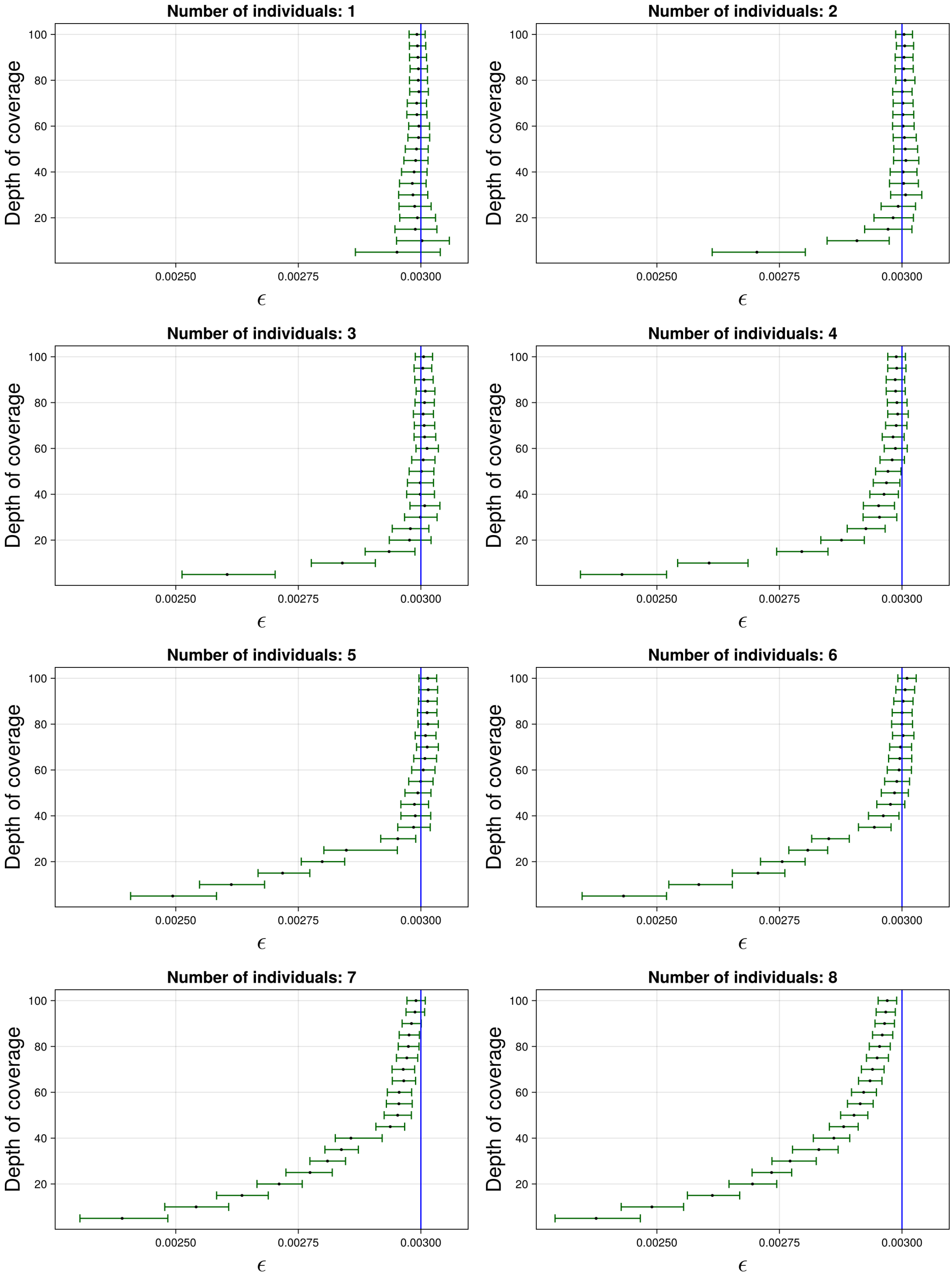

Figure S24: The 95% credible intervals for $\varepsilon$ with ancient population CHBS and anchor population KAR. The black dot indicates the median. The blue line indicates the true simulated value.

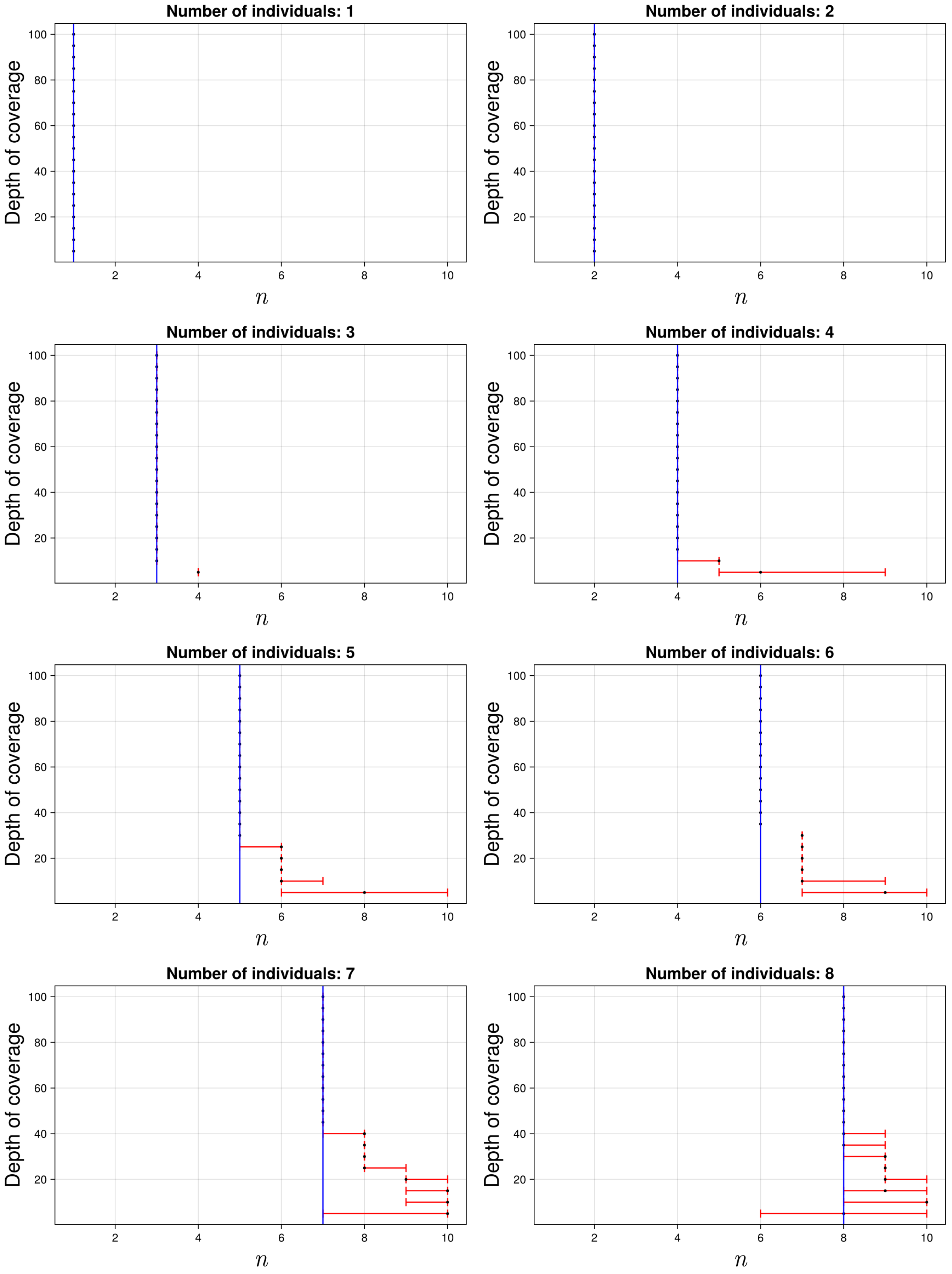

Figure S25: The 95% credible intervals for $n$ with ancient population CHBS and anchor population KAR. The black dot indicates the median. The blue line indicates the true simulated value.

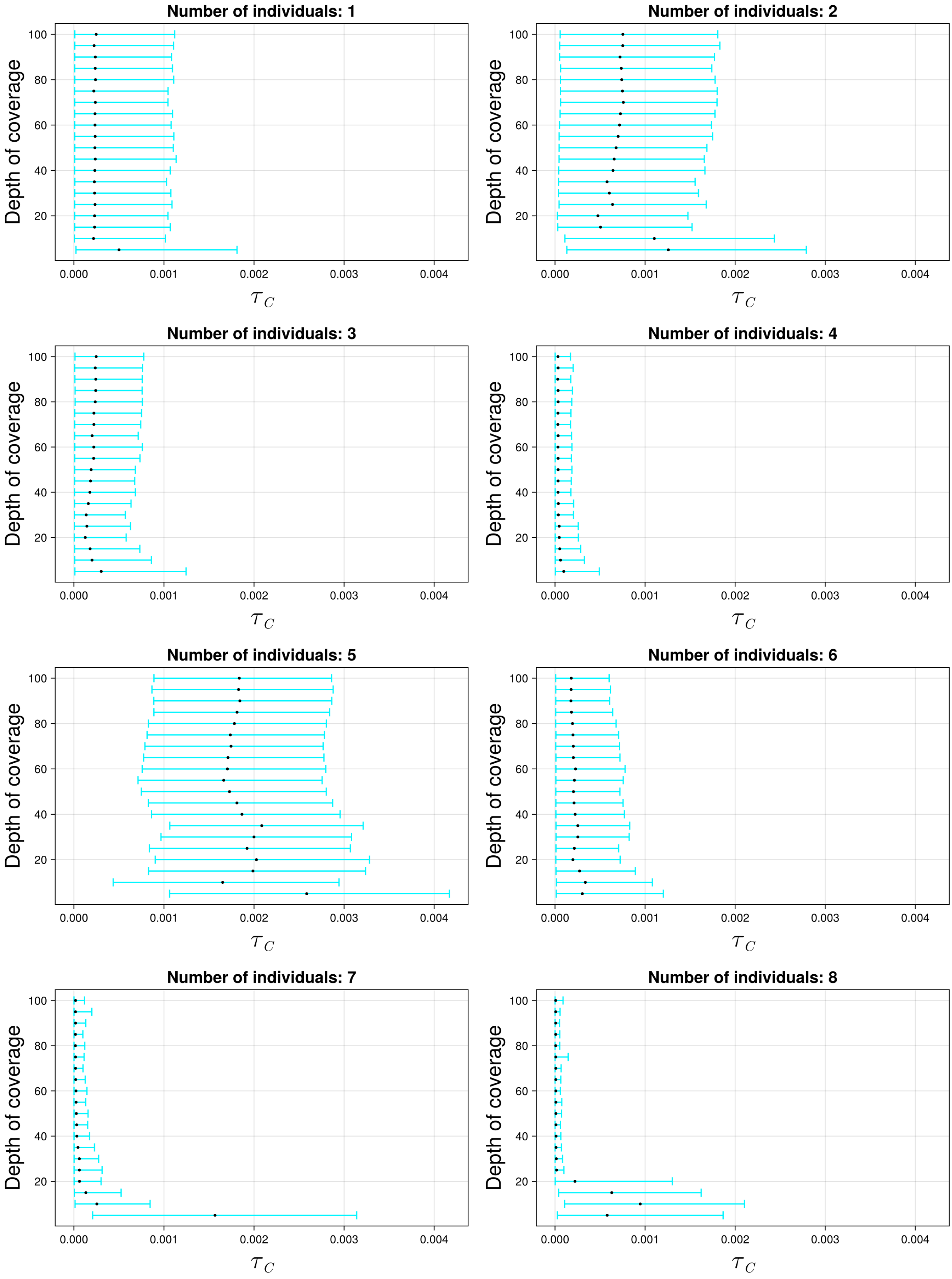

Figure S26: The 95% credible intervals for $\tau_C$ with ancient population YRIS and anchor population YRI. The black dot indicates the median.

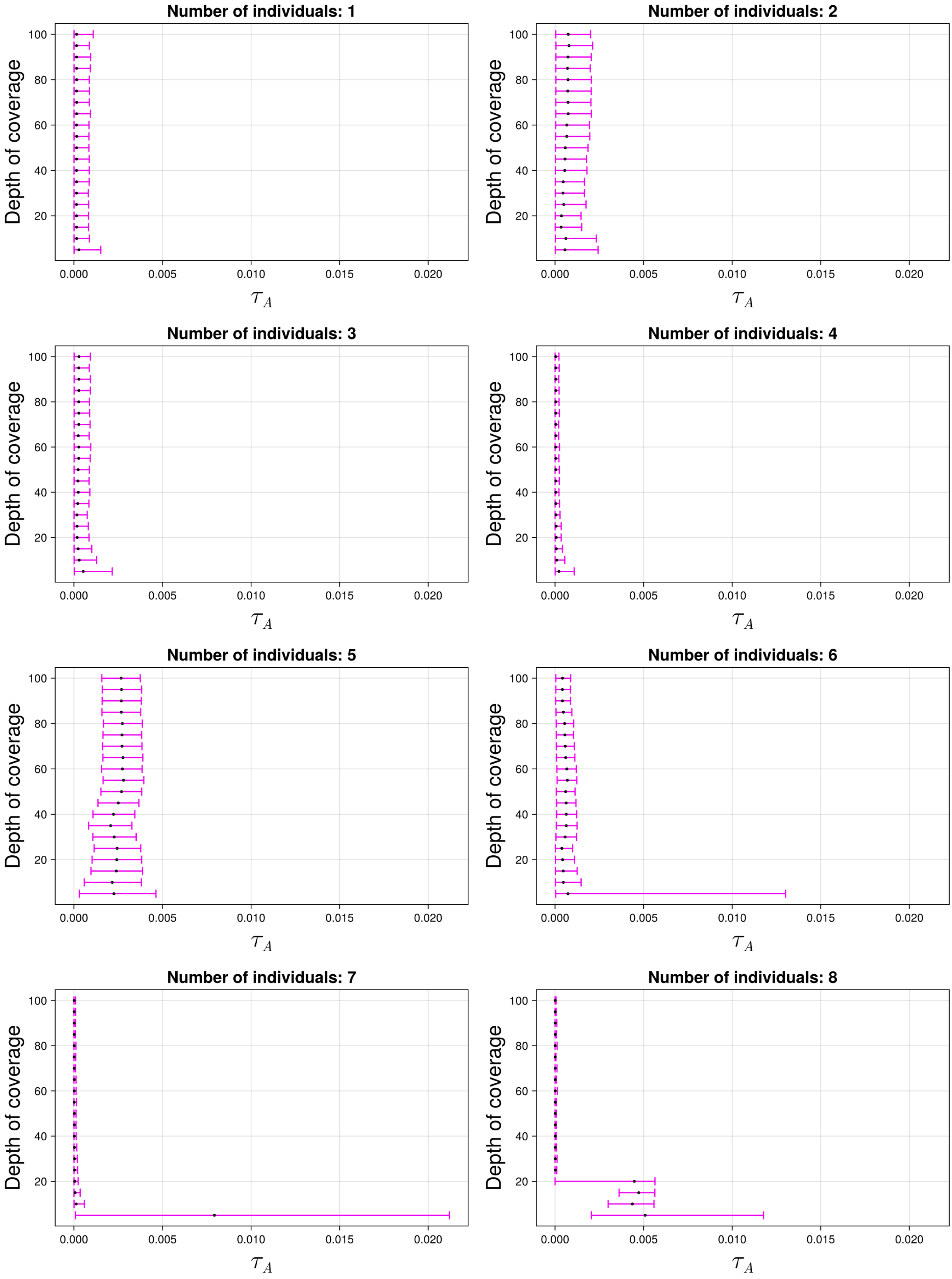

Figure S27: The 95% credible intervals for $\tau_A$ with ancient population YRIS and anchor population YRI. The black dot indicates the median.

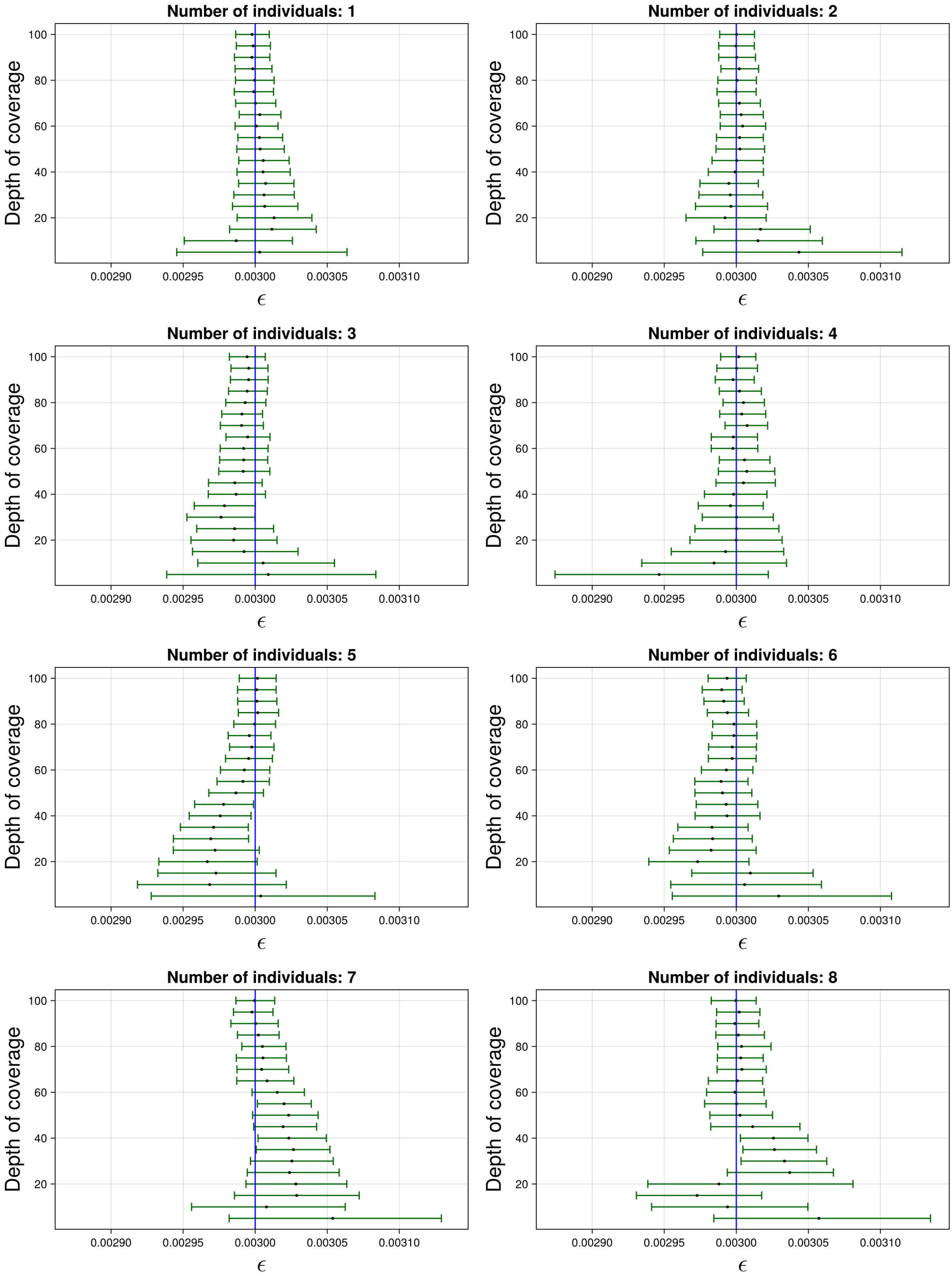

Figure S28: The 95% credible intervals for $\varepsilon$ with ancient population YRIS and anchor population YRI. The black dot indicates the median. The blue line indicates the true simulated value.

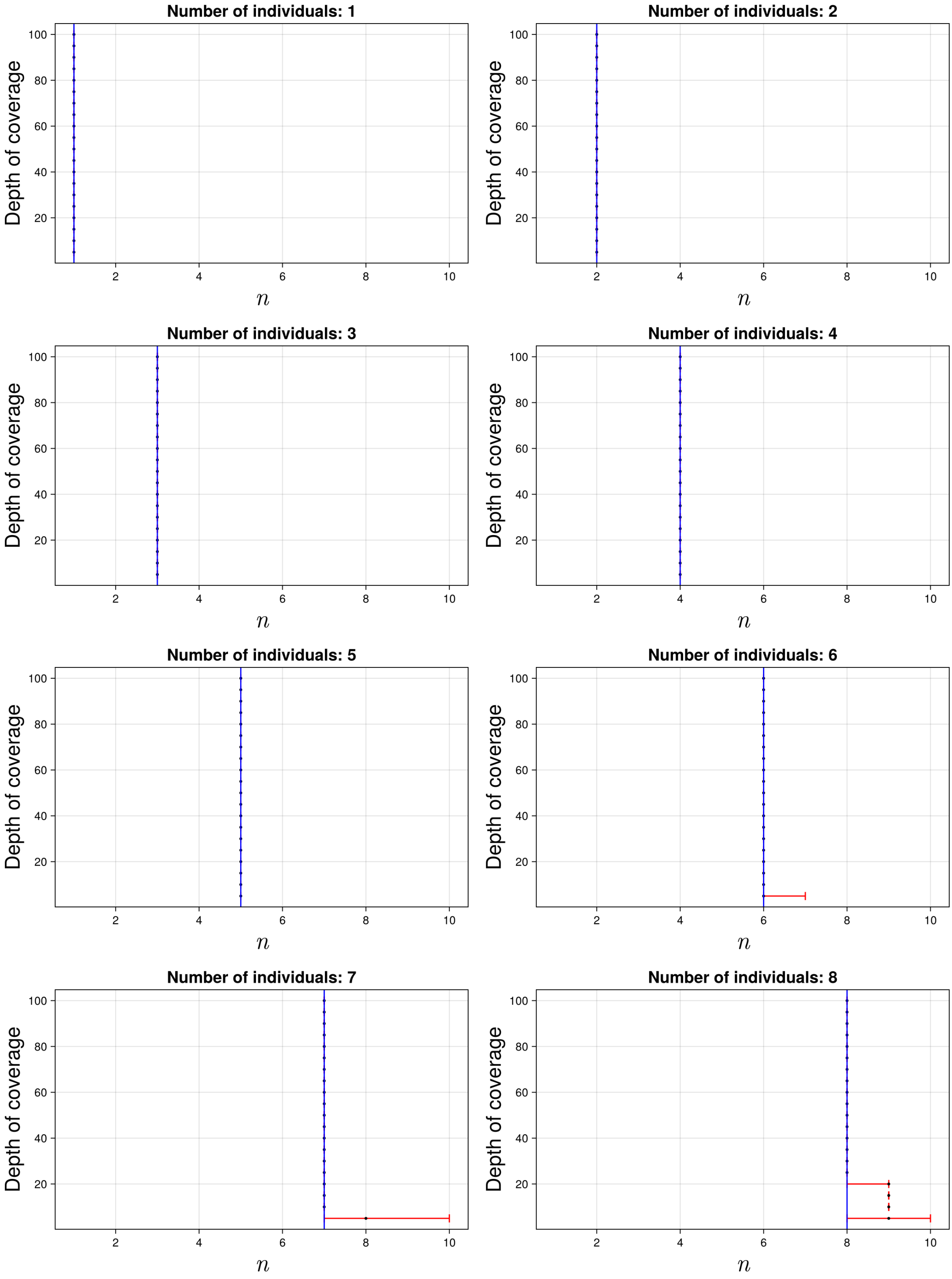

Figure S29: The 95% credible intervals for $n$ with ancient population YRIS and anchor population YRI. The black dot indicates the median. The blue line indicates the true simulated value.

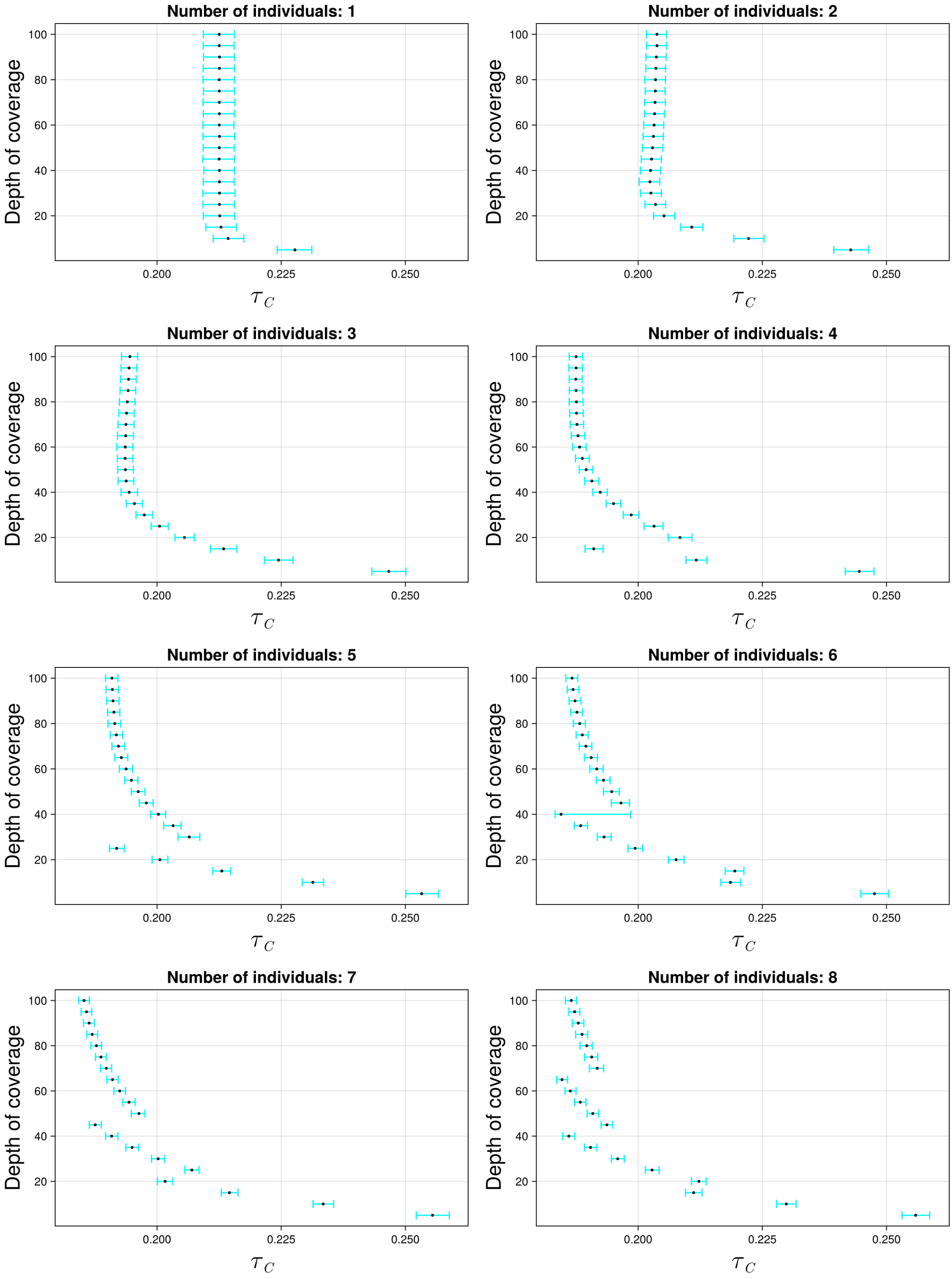

Figure S30: The 95% credible intervals for $\tau_C$ with ancient population YRIS and anchor population CEU. The black dot indicates the median.

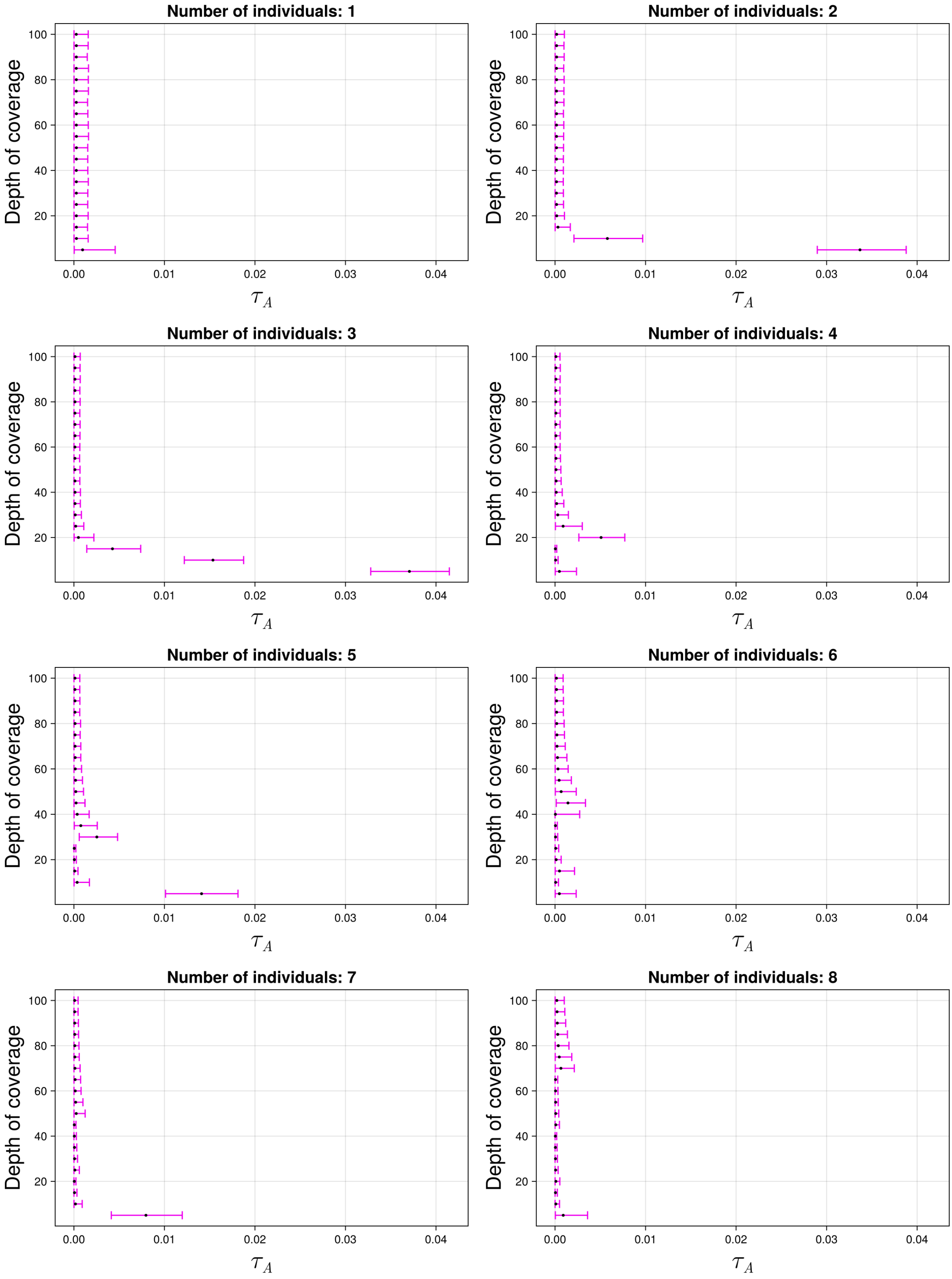

Figure S31: The 95% credible intervals for $\tau_A$ with ancient population YRIS and anchor population CEU. The black dot indicates the median.

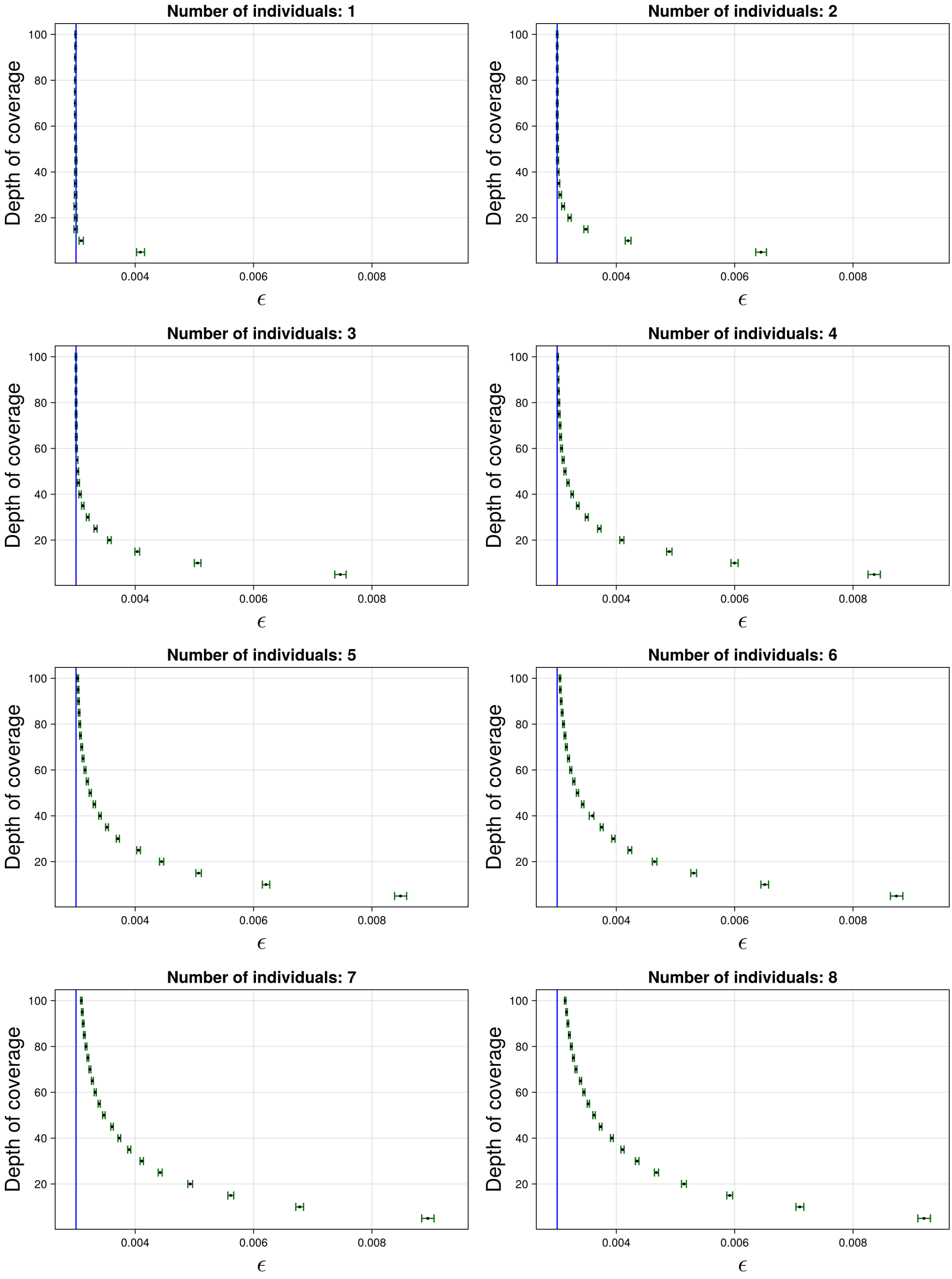

Figure S32: The 95% credible intervals for $\varepsilon$ with ancient population YRIS and anchor population CEU. The black dot indicates the median. The blue line indicates the true simulated value.

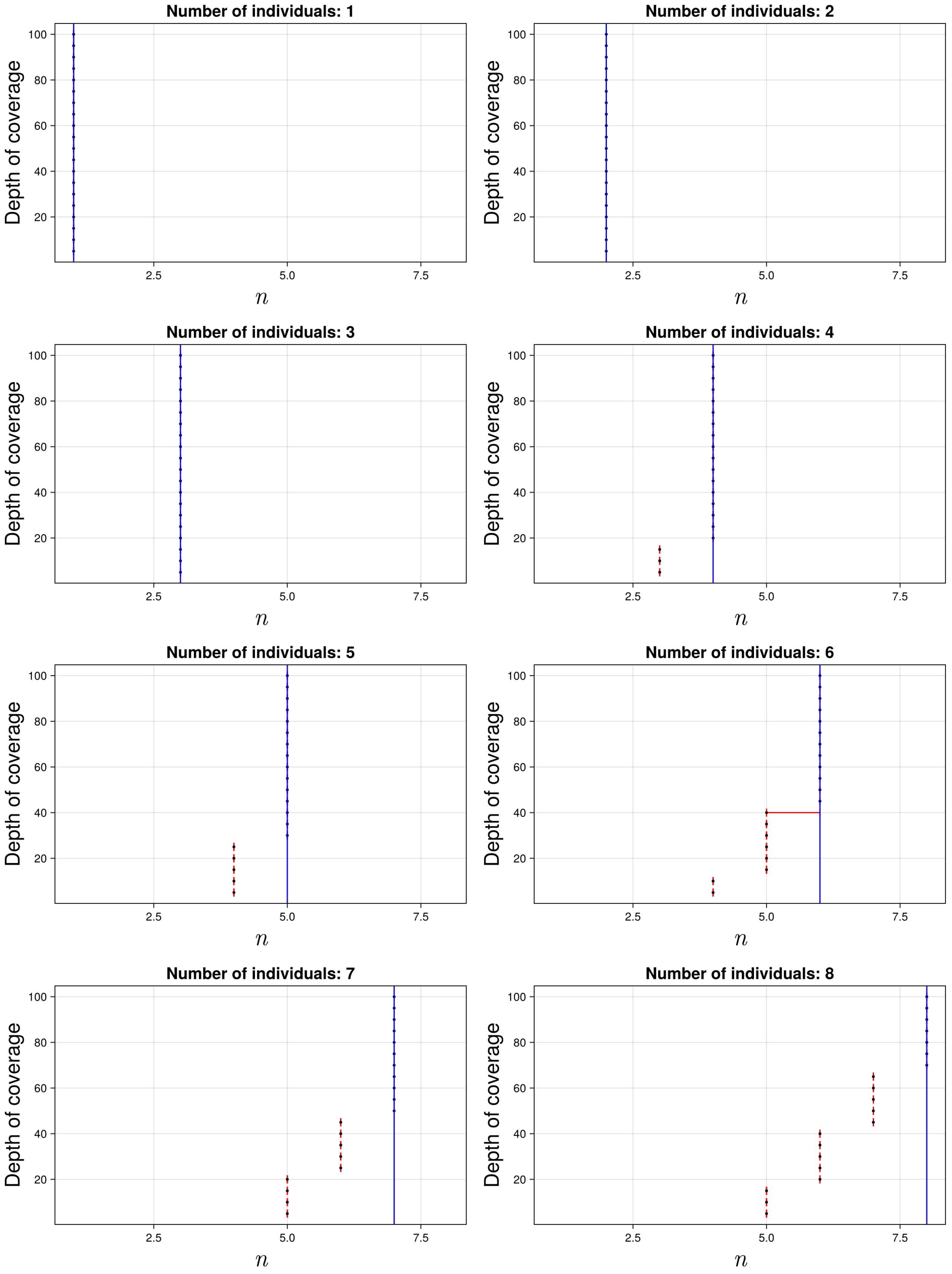

Figure S33: The 95% credible intervals for $n$ with ancient population YRIS and anchor population CEU. The black dot indicates the median. The blue line indicates the true simulated value.

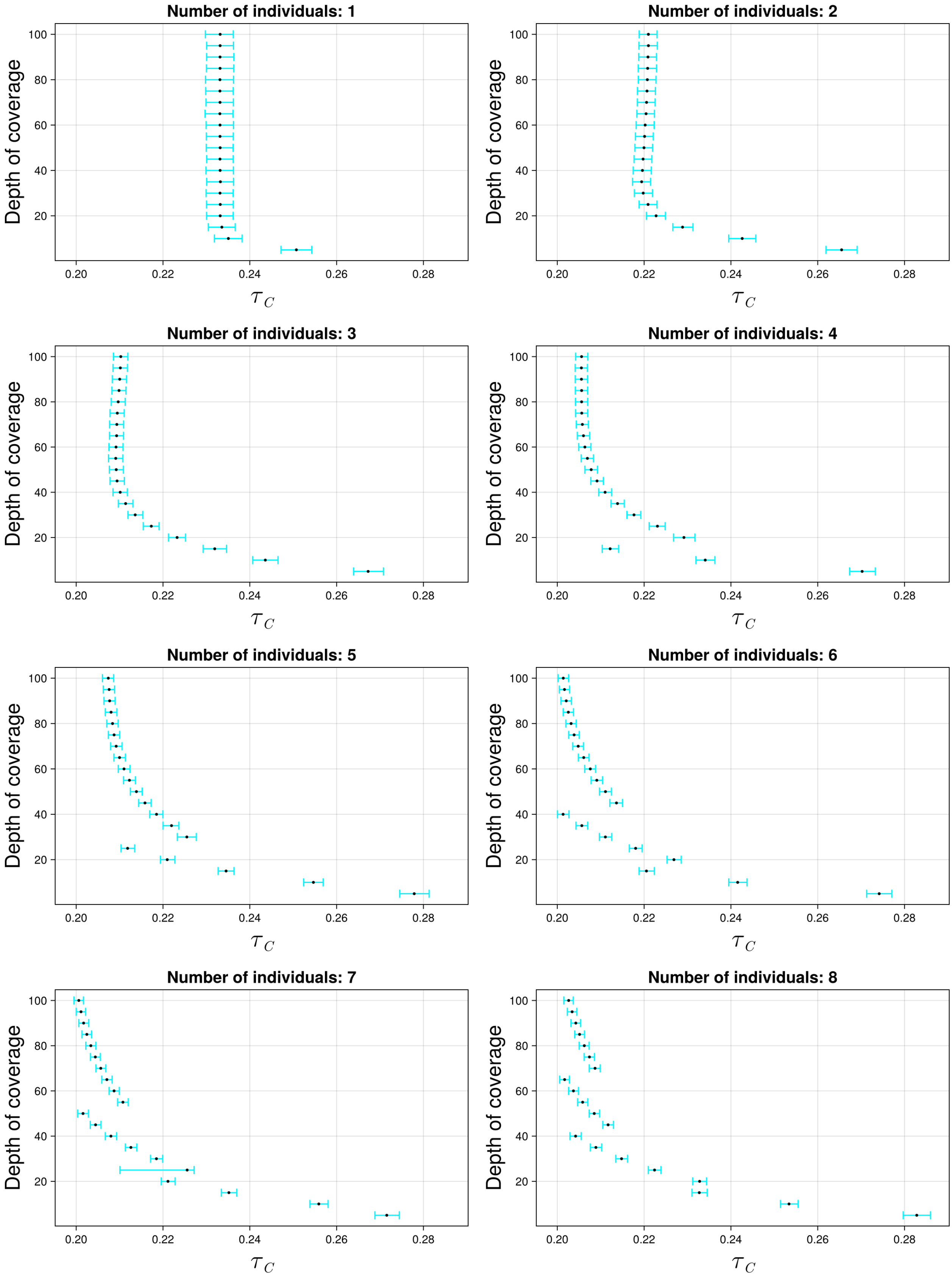

Figure S34: The 95% credible intervals for $\tau_C$ with ancient population YRIS and anchor population CHB. The black dot indicates the median.

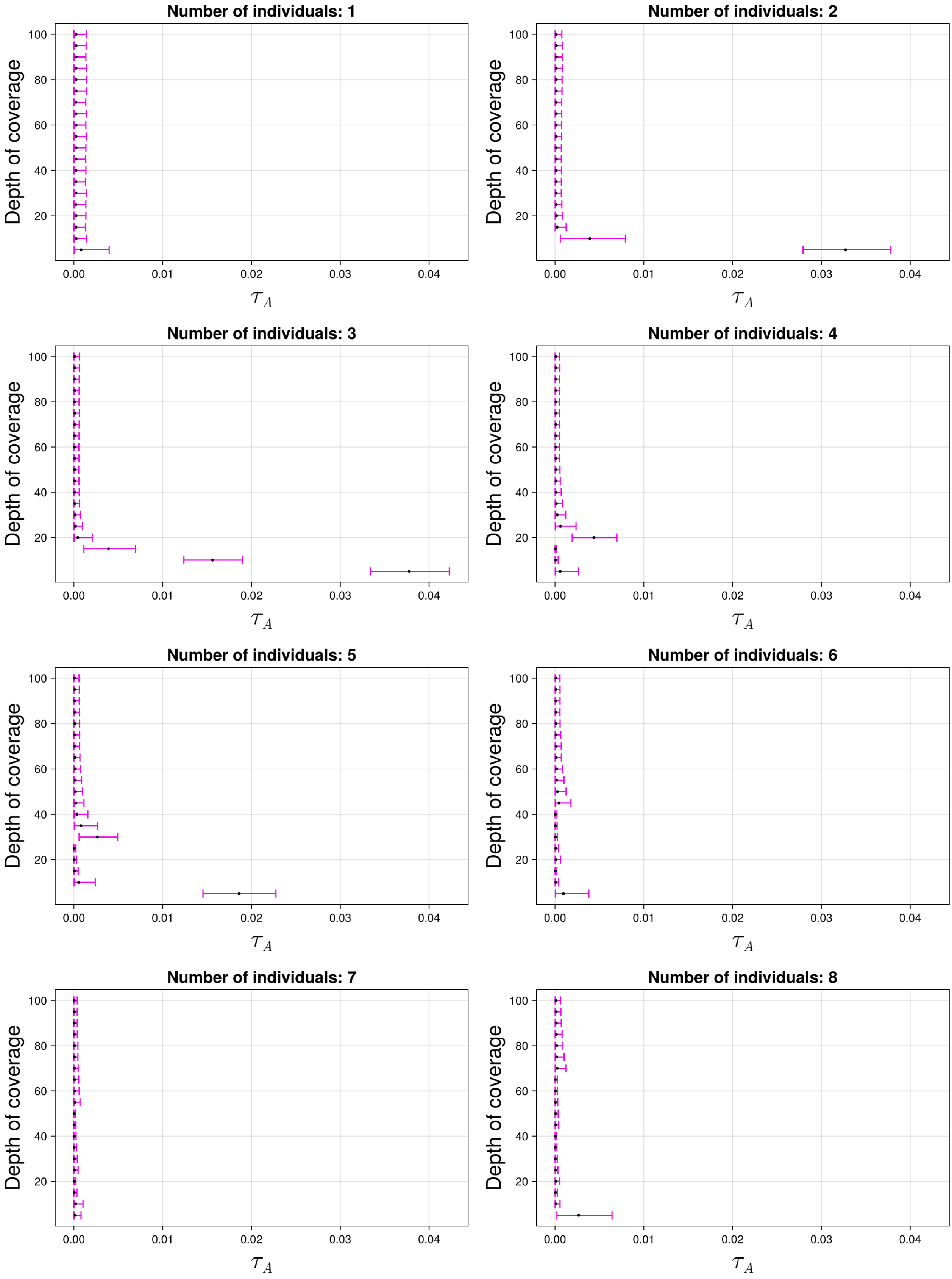

Figure S35: The 95% credible intervals for $\tau_A$ with ancient population YRIS and anchor population CHB. The black dot indicates the median.

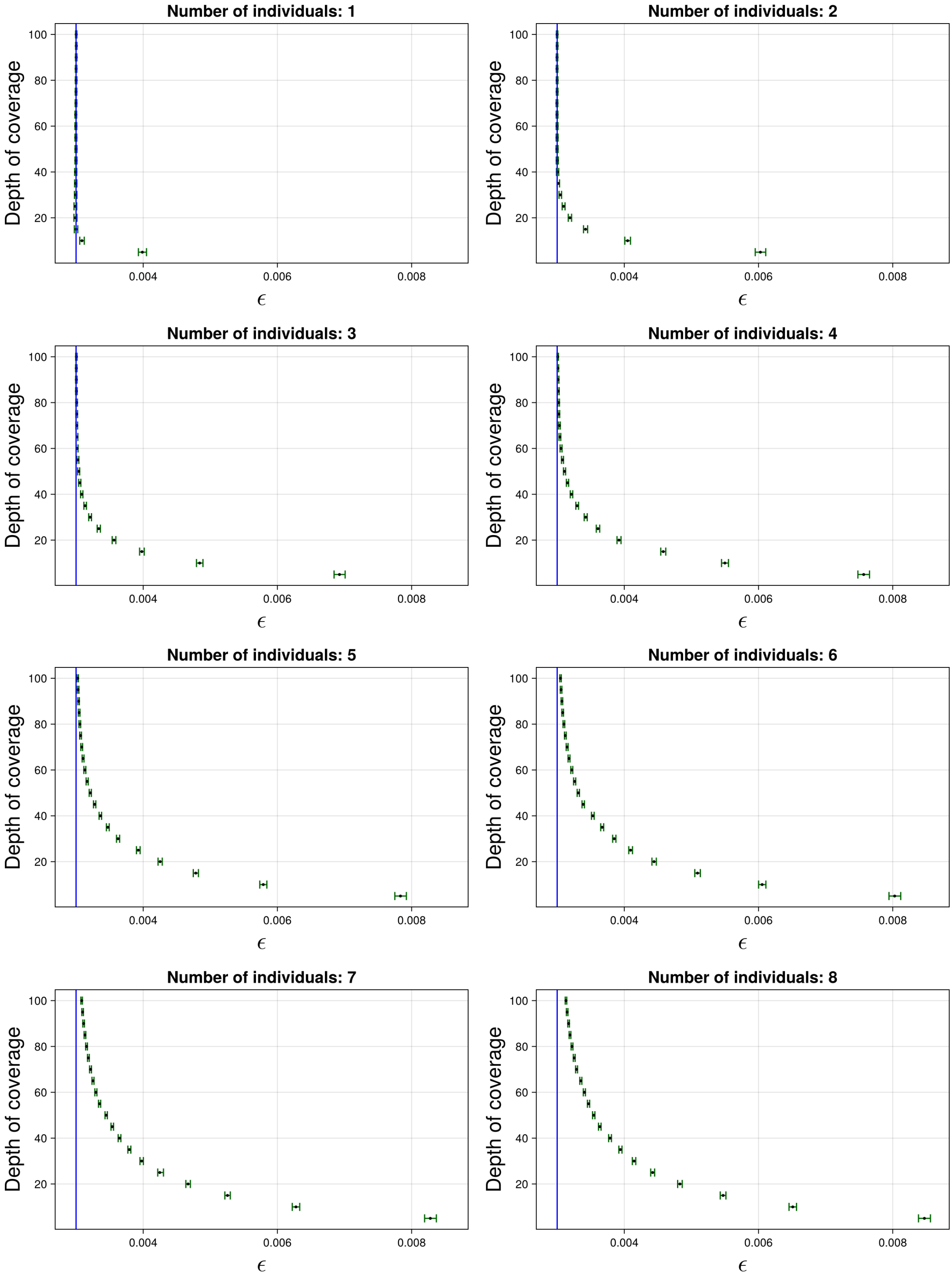

Figure S36: The 95% credible intervals for $\varepsilon$ with ancient population YRIS and anchor population CHB. The black dot indicates the median. The blue line indicates the true simulated value.

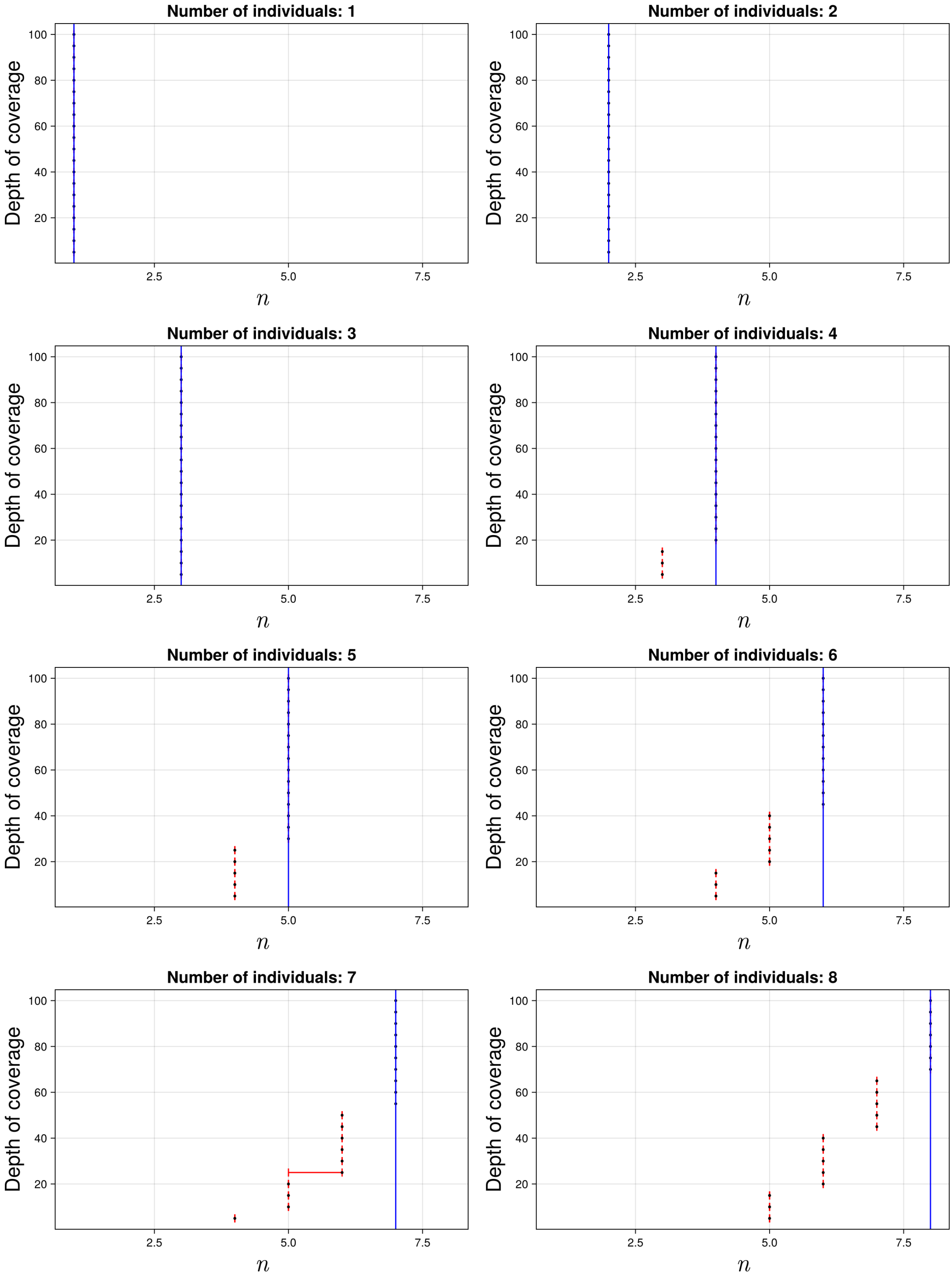

Figure S37: The 95% credible intervals for $n$ with ancient population YRIS and anchor population CHB. The black dot indicates the median. The blue line indicates the true simulated value.

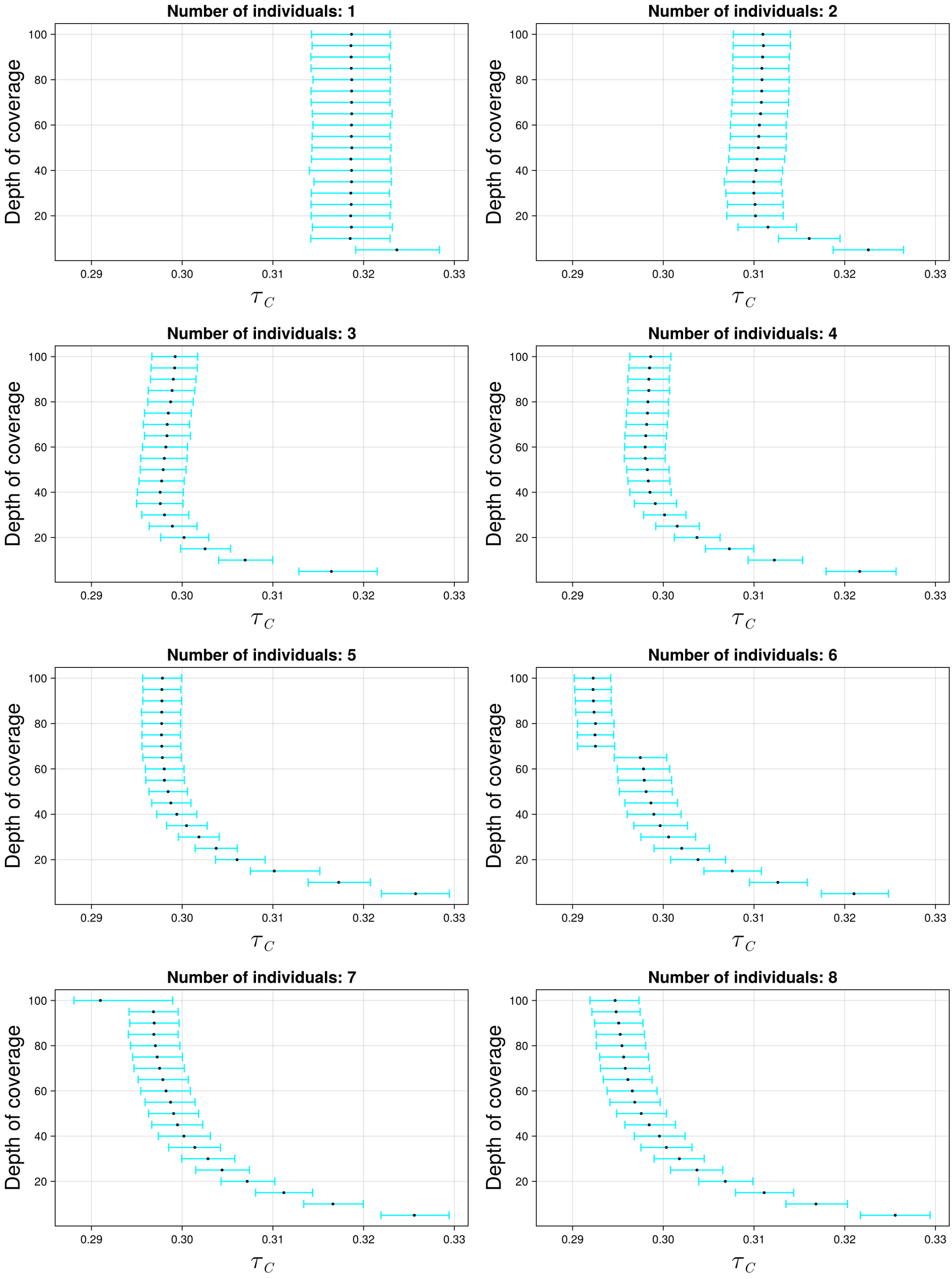

Figure S38: The 95% credible intervals for $\tau_C$ with ancient population YRIS and anchor population KAR. The black dot indicates the median.

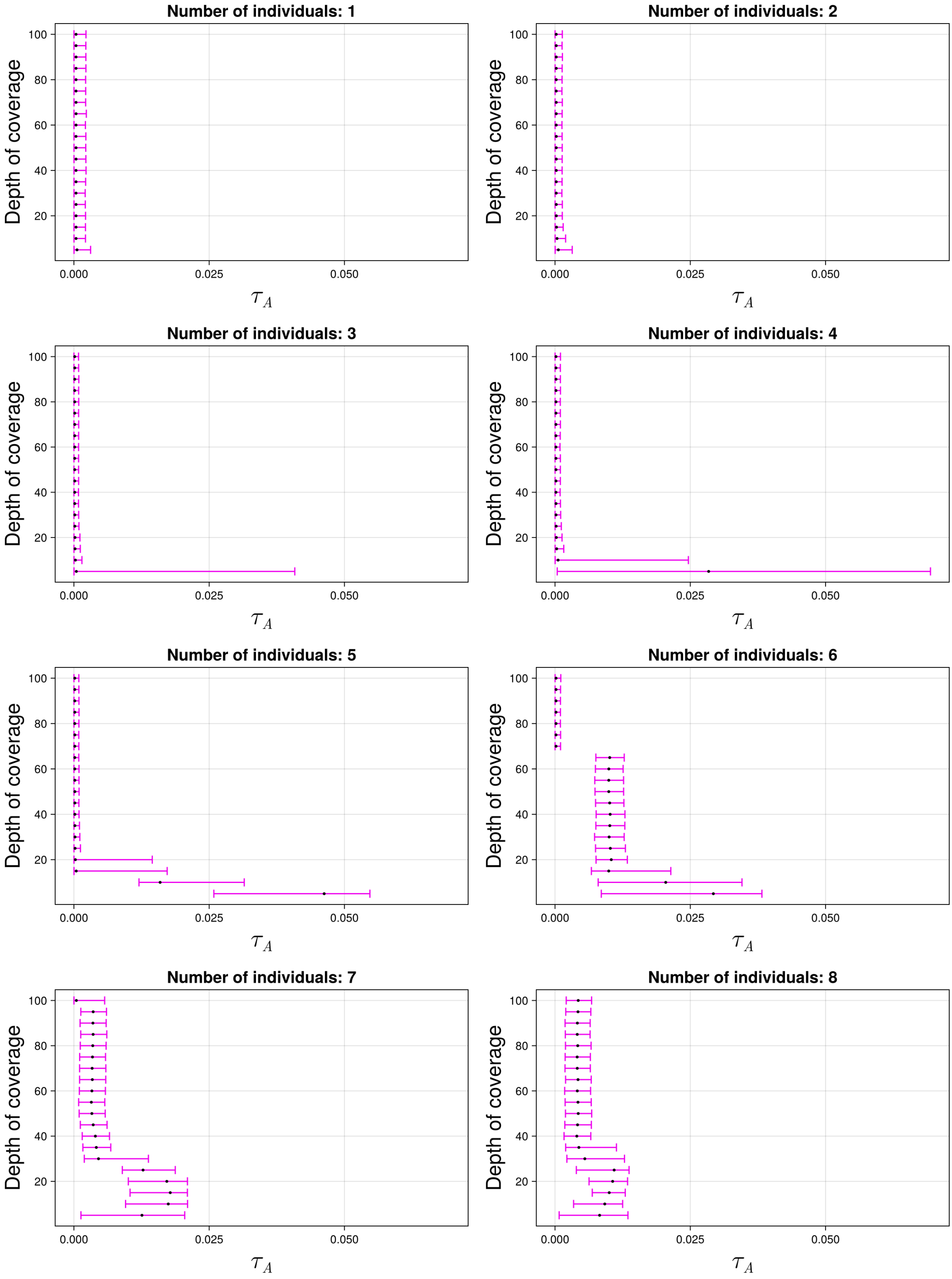

Figure S39: The 95% credible intervals for $\tau_A$ with ancient population YRIS and anchor population KAR. The black dot indicates the median.

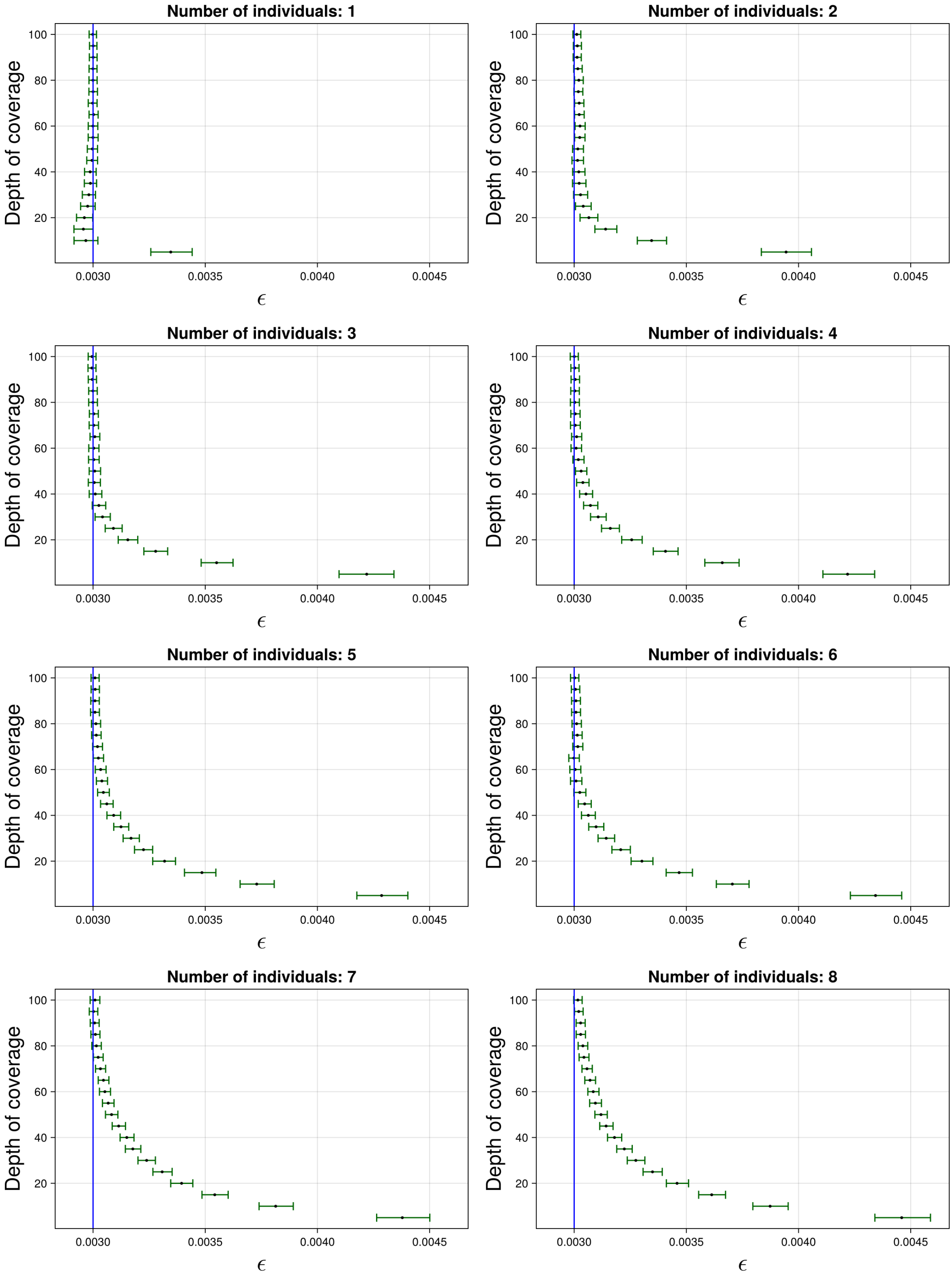

Figure S40: The 95% credible intervals for $\varepsilon$ with ancient population YRIS and anchor population KAR. The black dot indicates the median. The blue line indicates the true simulated value.

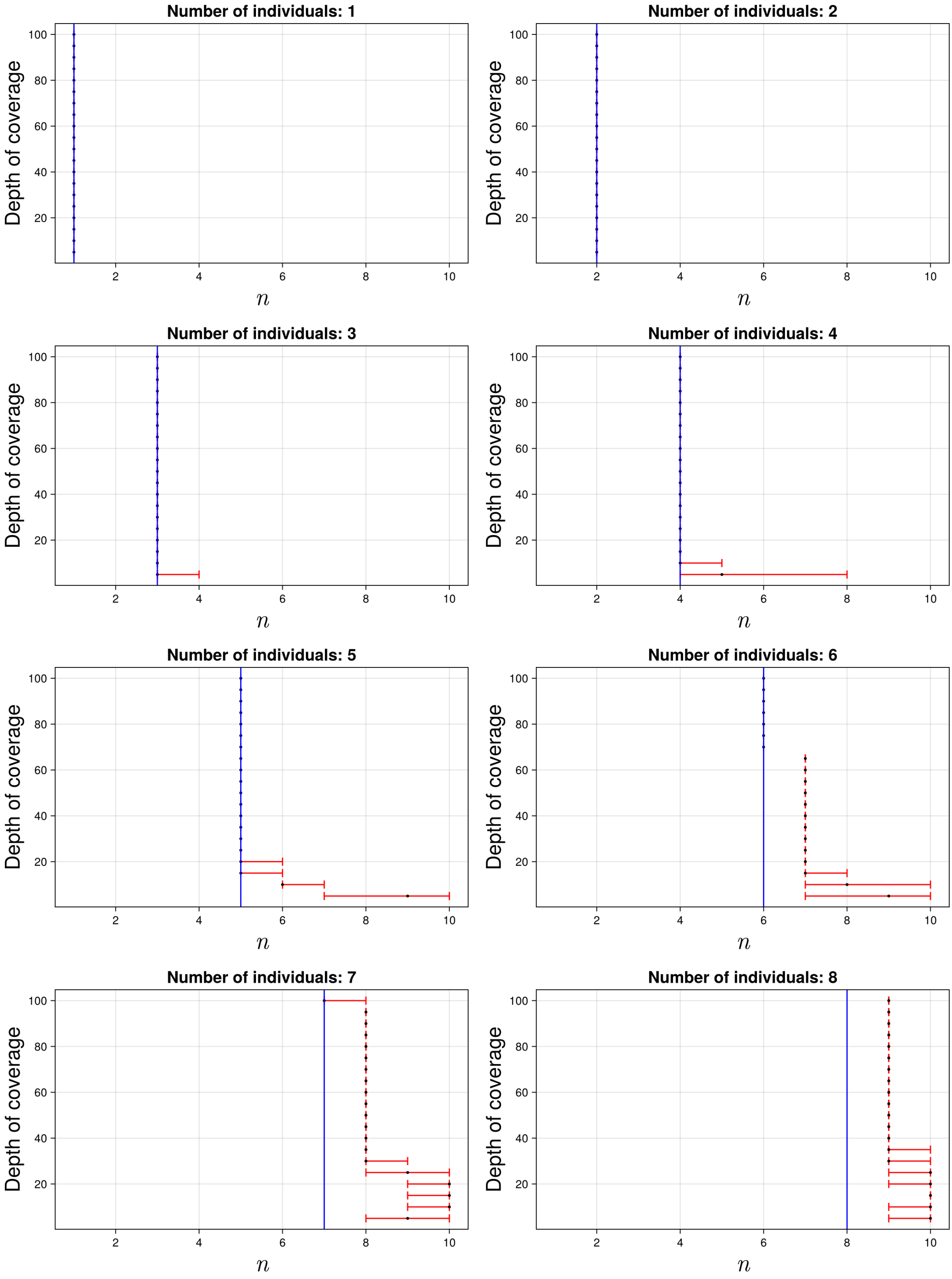

Figure S41: The 95% credible intervals for $n$ with ancient population YRIS and anchor population KAR. The black dot indicates the median. The blue line indicates the true simulated value.

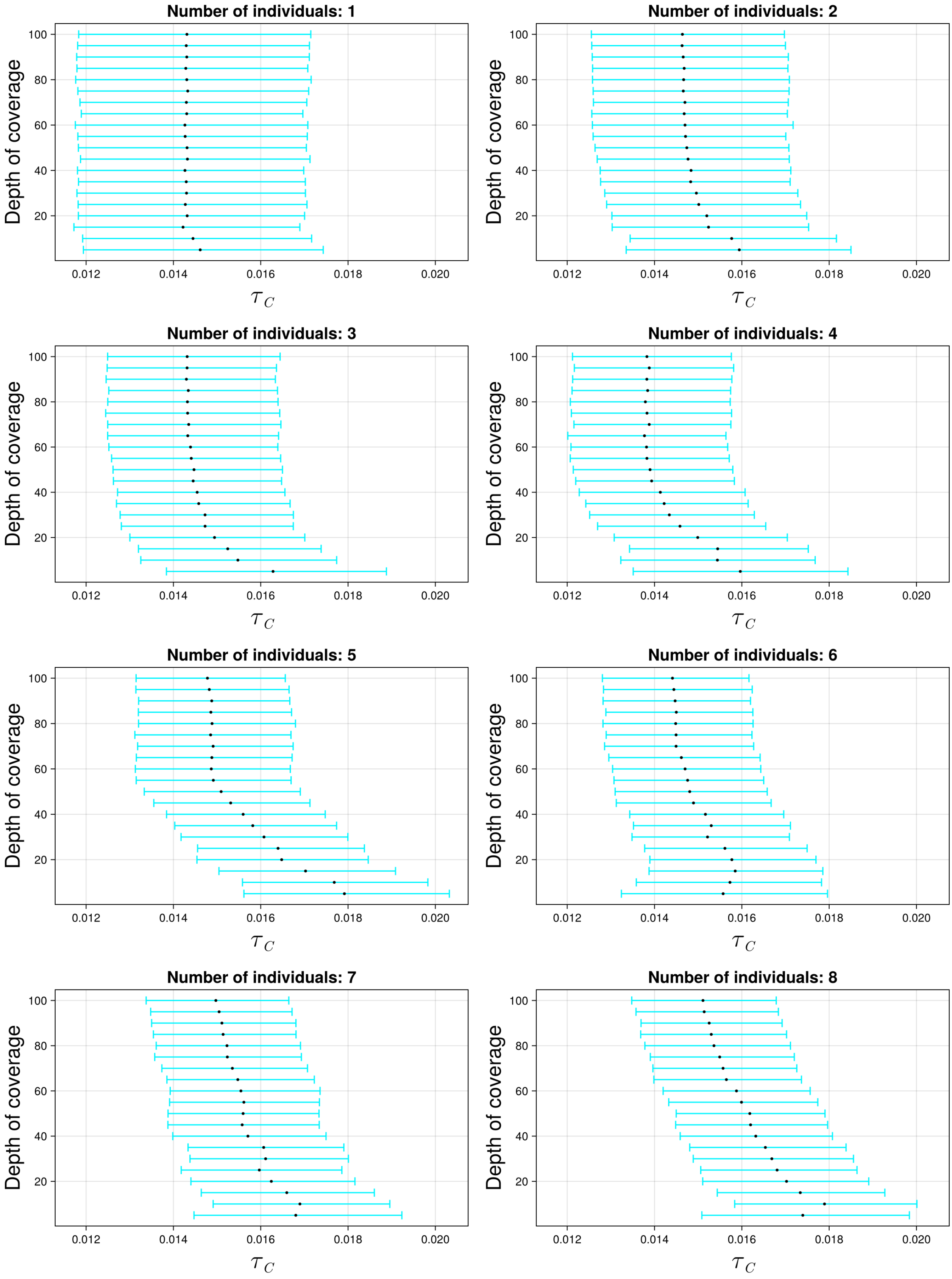

Figure S42: The 95% credible intervals for $\tau_C$ with ancient population CEUS and anchor population YRI. The black dot indicates the median.

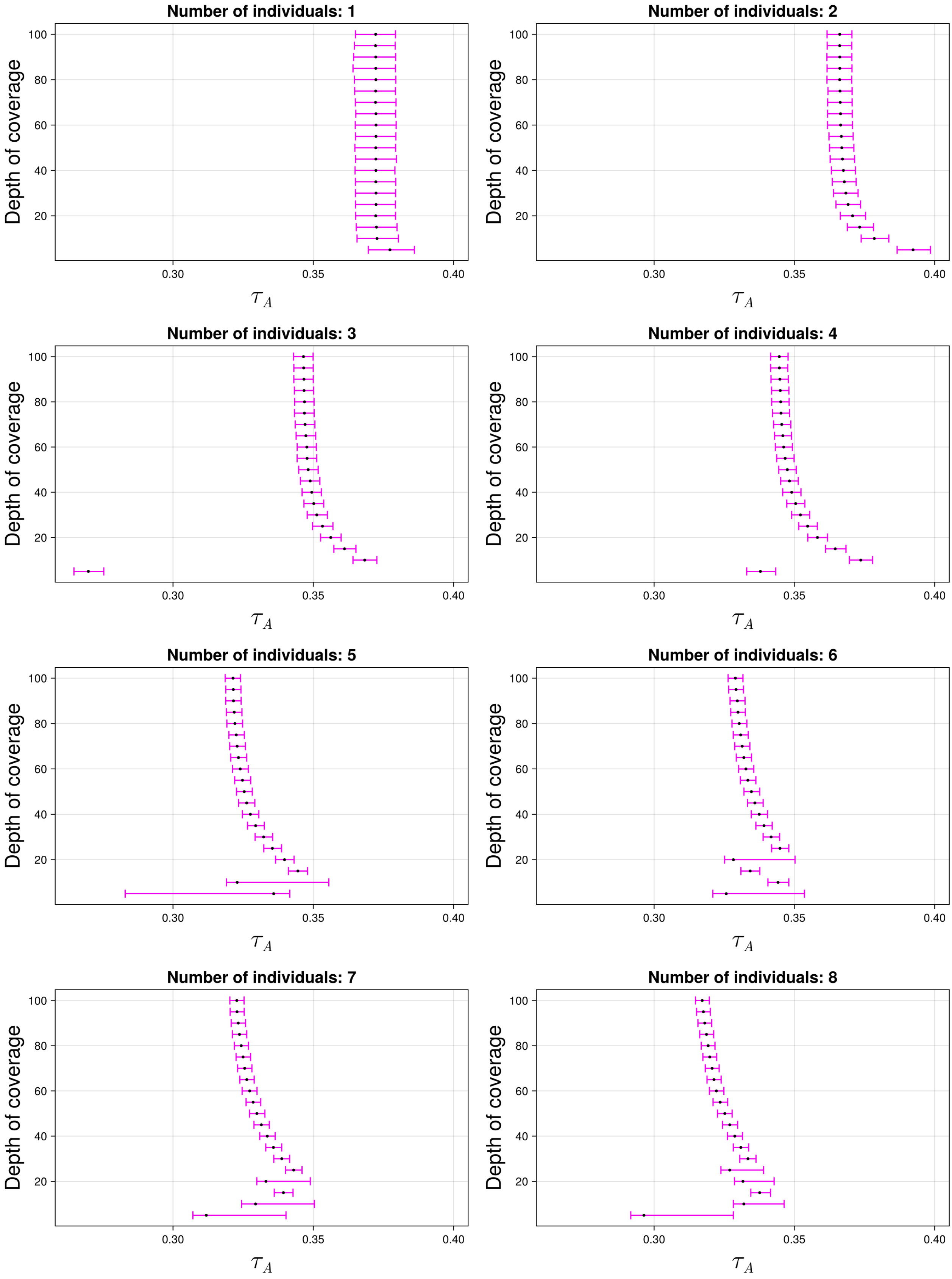

Figure S43: The 95% credible intervals for $\tau_A$ with ancient population CEUS and anchor population YRI. The black dot indicates the median.

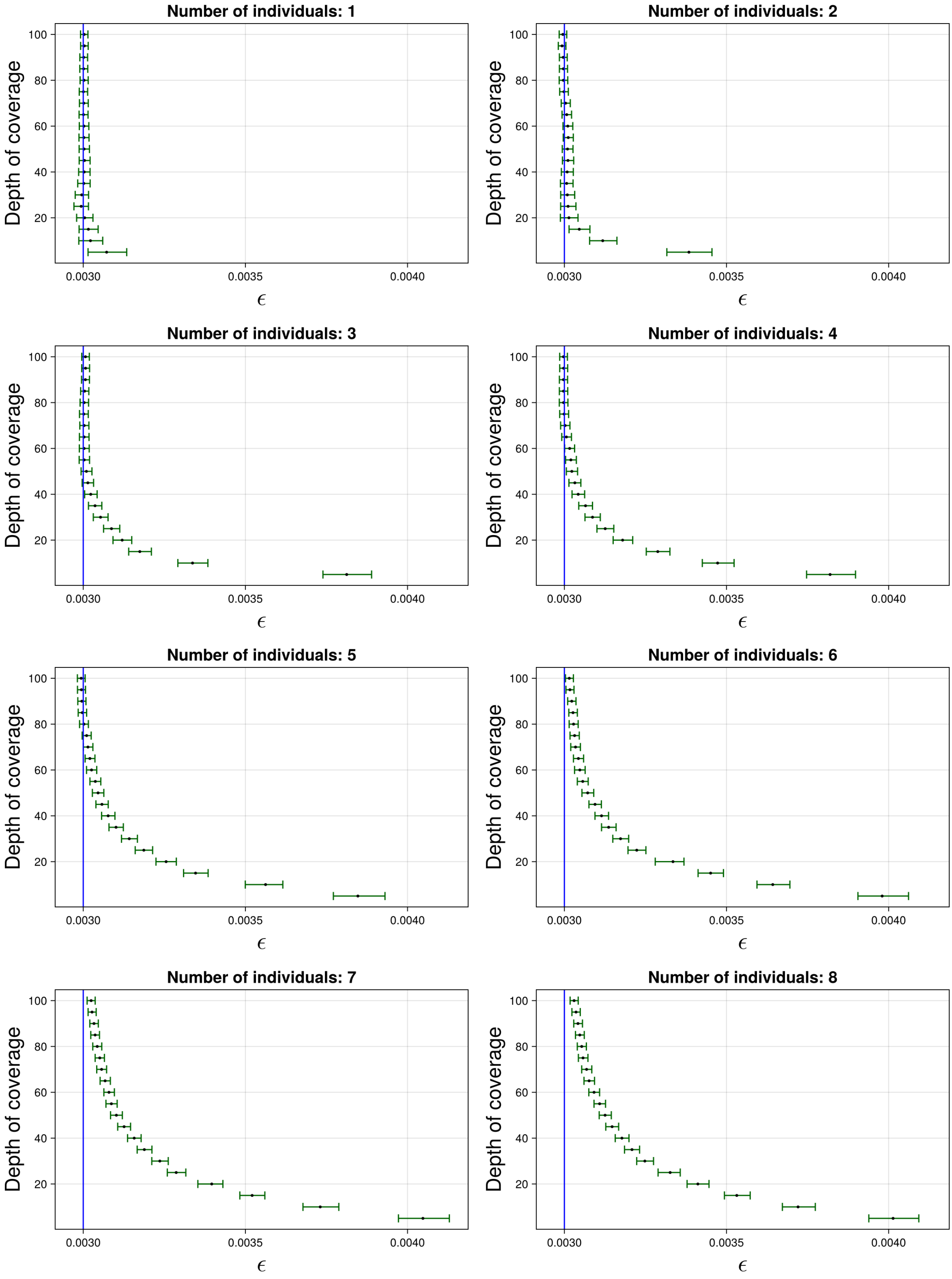

Figure S44: The 95% credible intervals for $\varepsilon$ with ancient population CEUS and anchor population YRI. The black dot indicates the median. The blue line indicates the true simulated value.

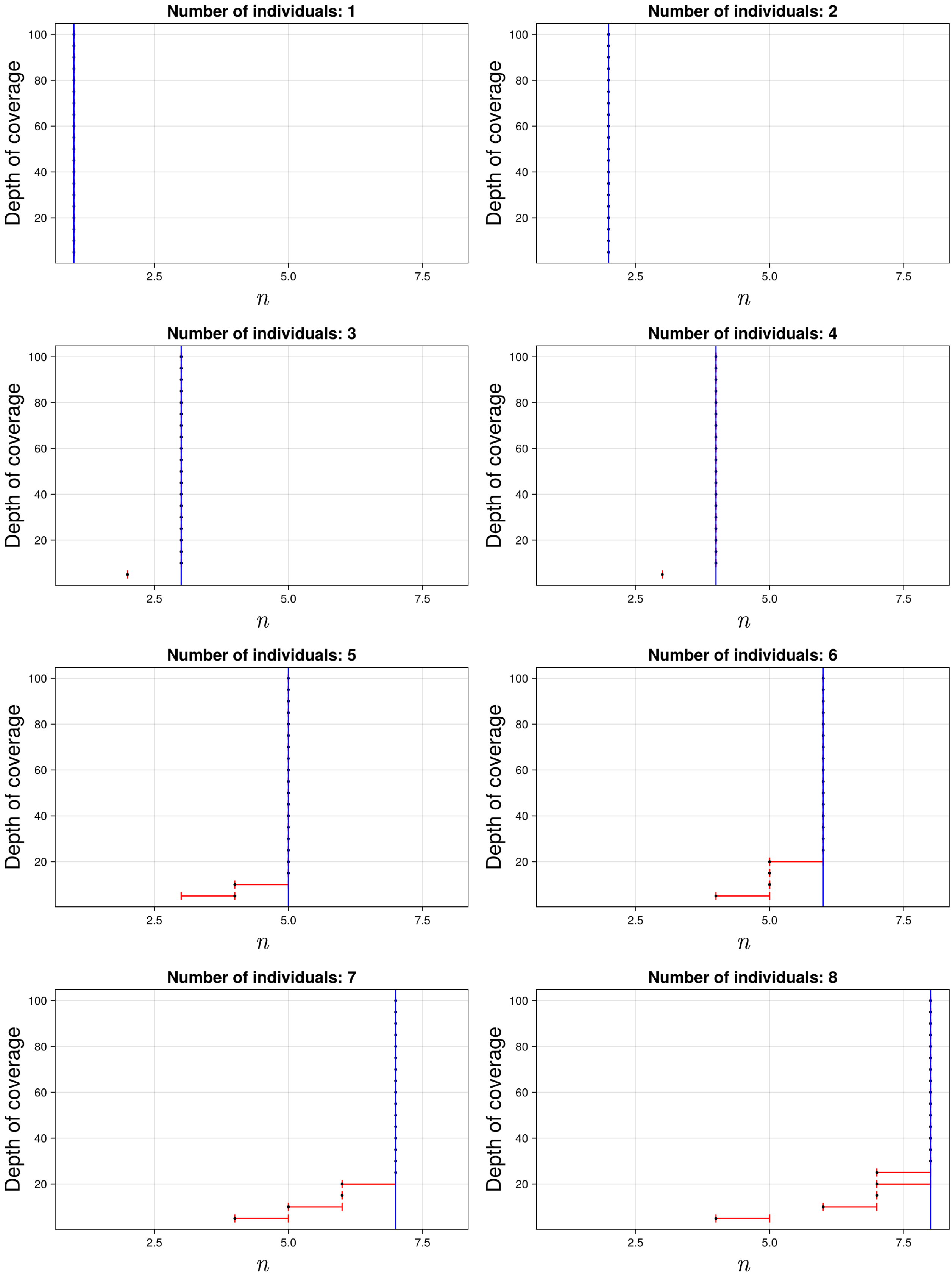

Figure S45: The 95% credible intervals for $n$ with ancient population CEUS and anchor population YRI. The black dot indicates the median. The blue line indicates the true simulated value.

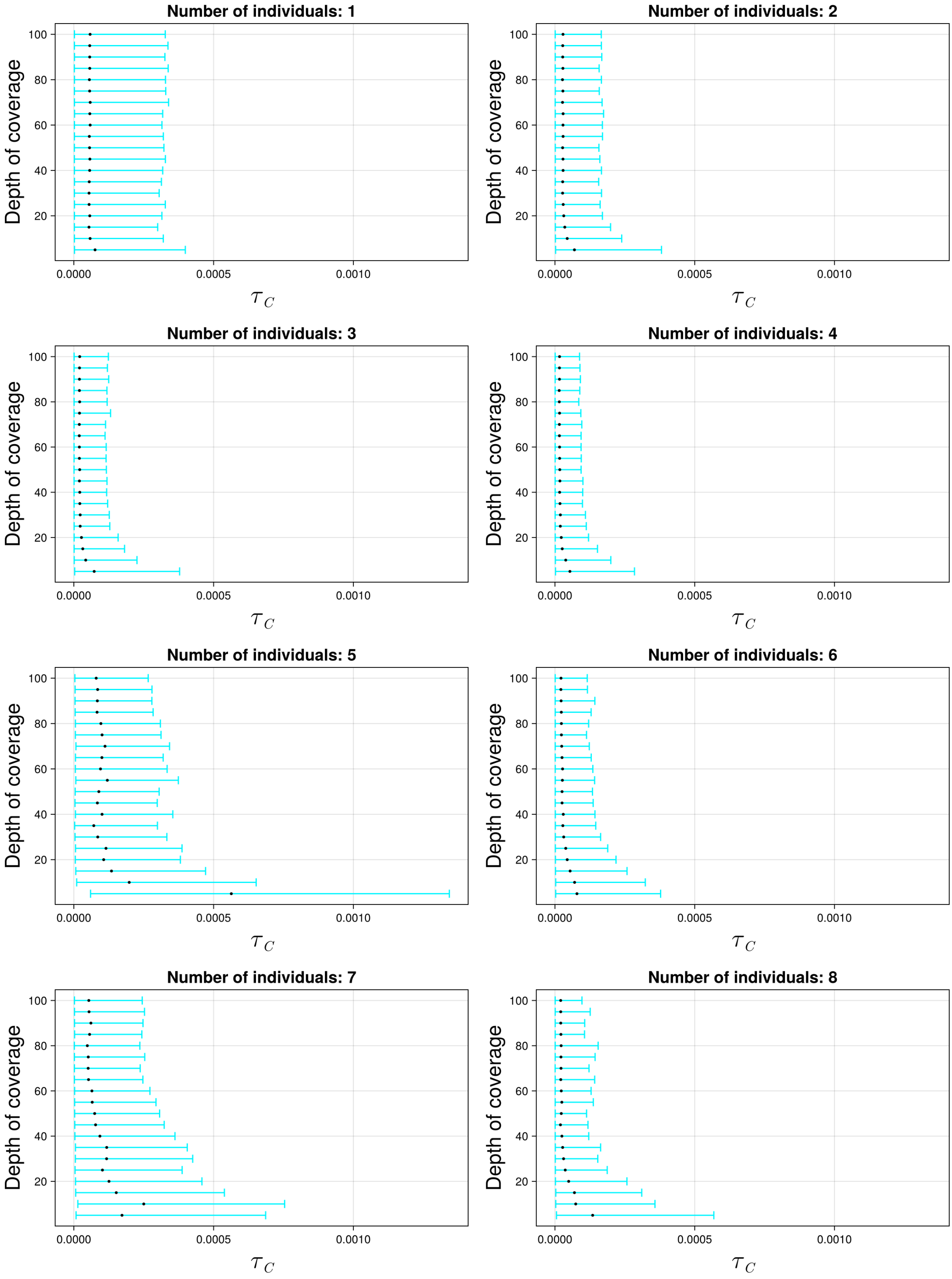

Figure S46: The 95% credible intervals for $\tau_C$ with ancient population CEUS and anchor population CEU. The black dot indicates the median.

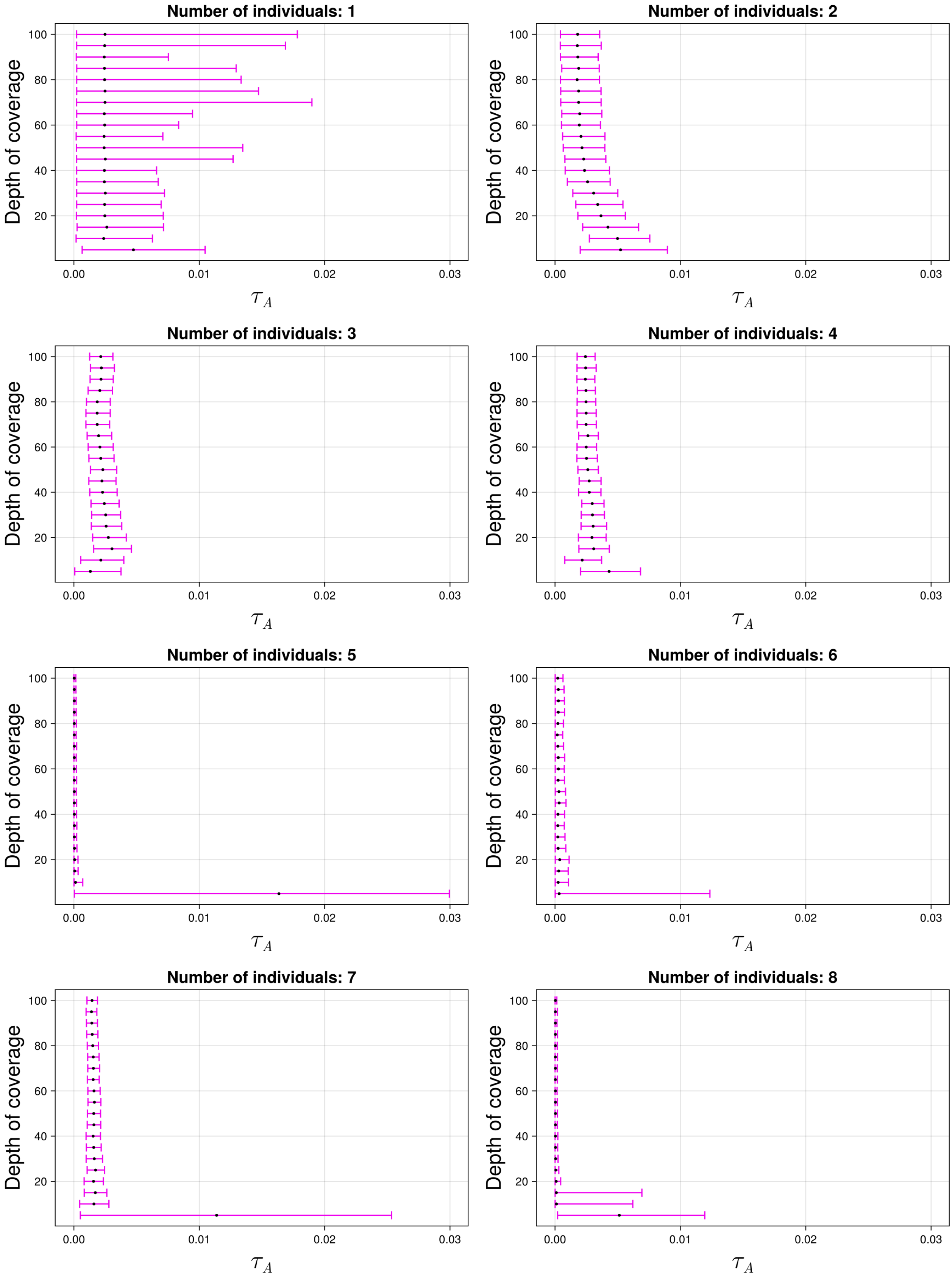

Figure S47: The 95% credible intervals for $\tau_A$ with ancient population CEUS and anchor population CEU. The black dot indicates the median.

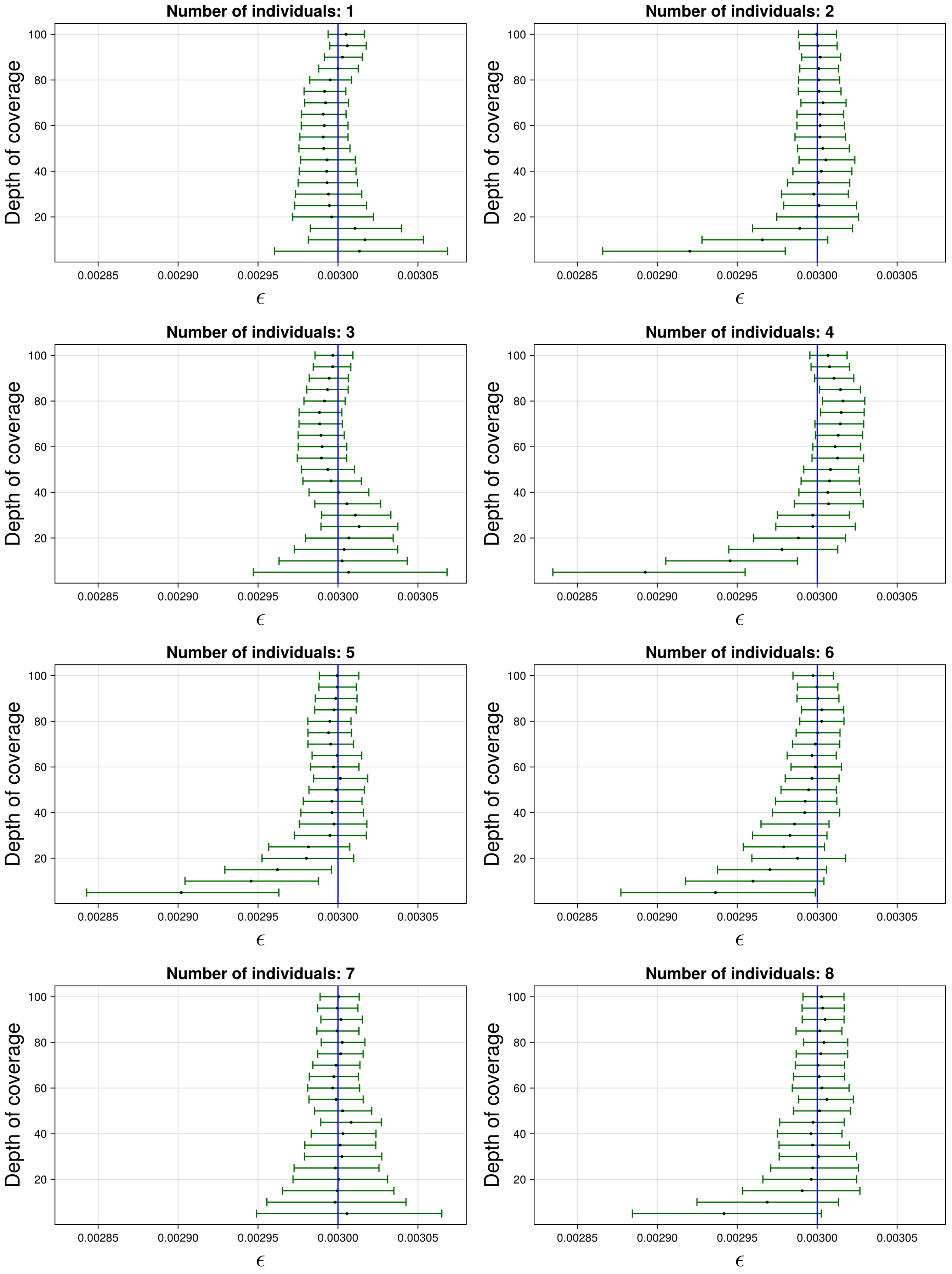

Figure S48: The 95% credible intervals for $\varepsilon$ with ancient population CEUS and anchor population CEU. The black dot indicates the median. The blue line indicates the true simulated value.

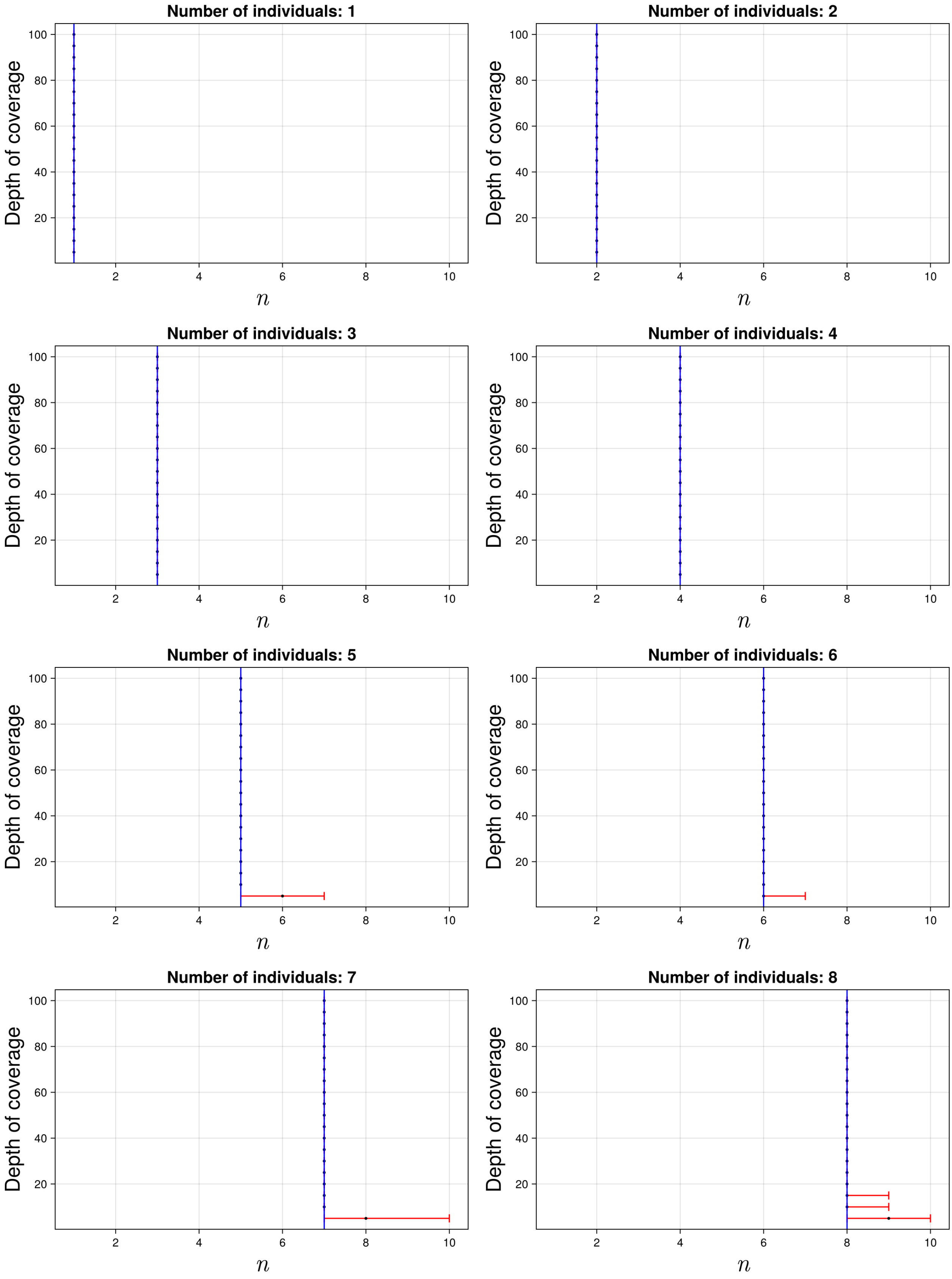

Figure S49: The 95% credible intervals for $n$ with ancient population CEUS and anchor population CEU. The black dot indicates the median. The blue line indicates the true simulated value.

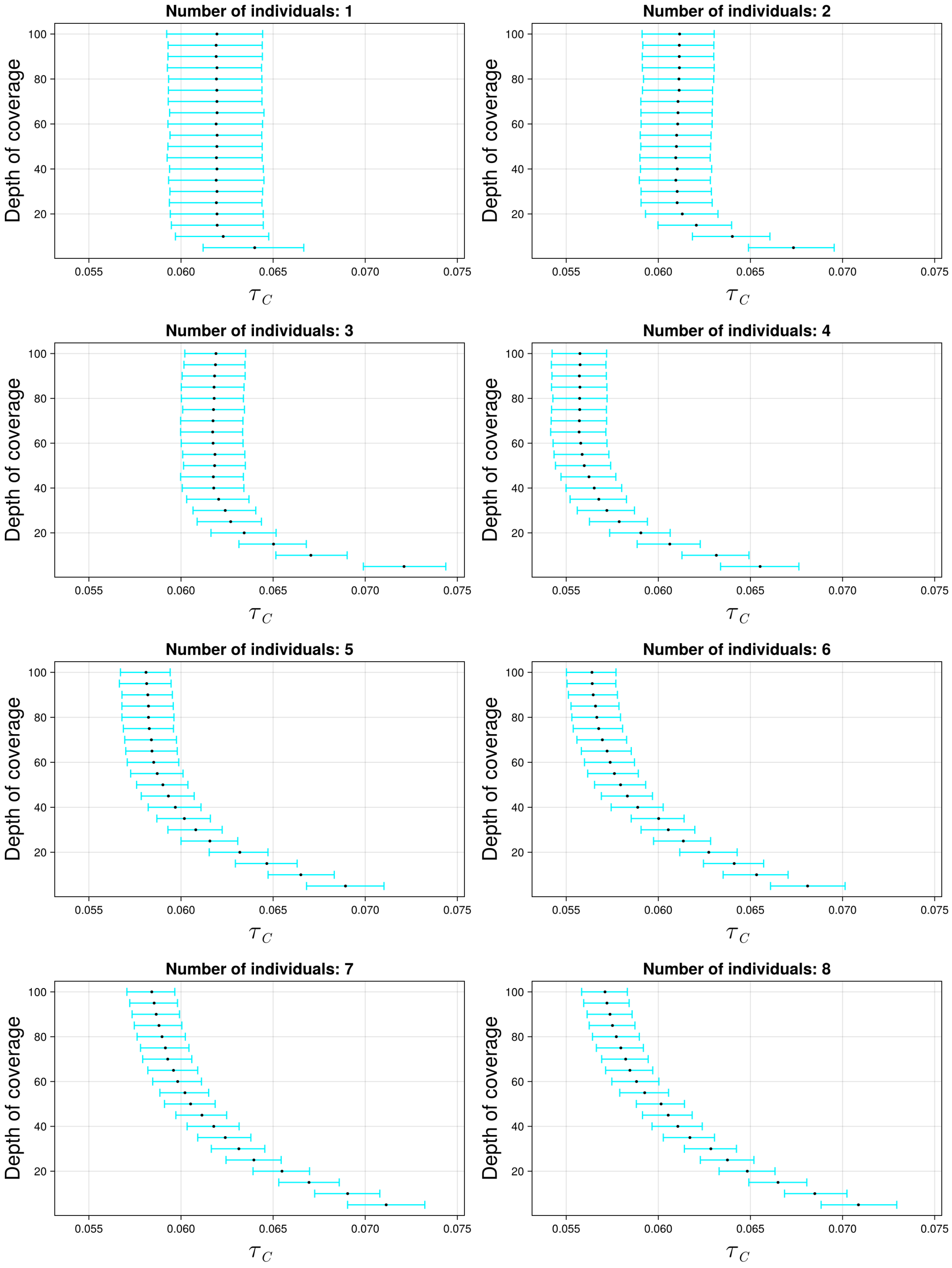

Figure S50: The 95% credible intervals for $\tau_C$ with ancient population CEUS and anchor population CHB. The black dot indicates the median.

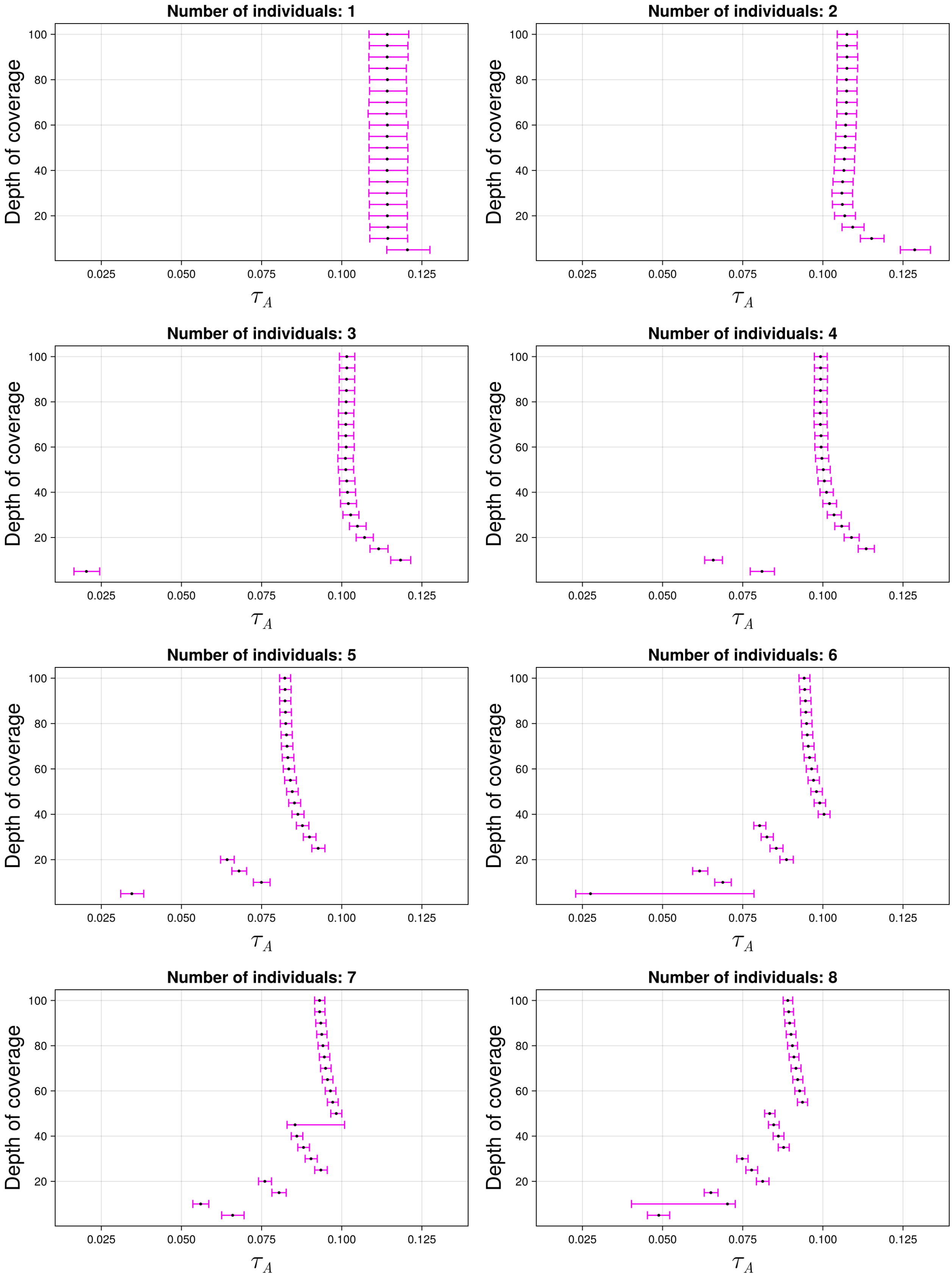

Figure S51: The 95% credible intervals for $\tau_A$ with ancient population CEUS and anchor population CHB. The black dot indicates the median.

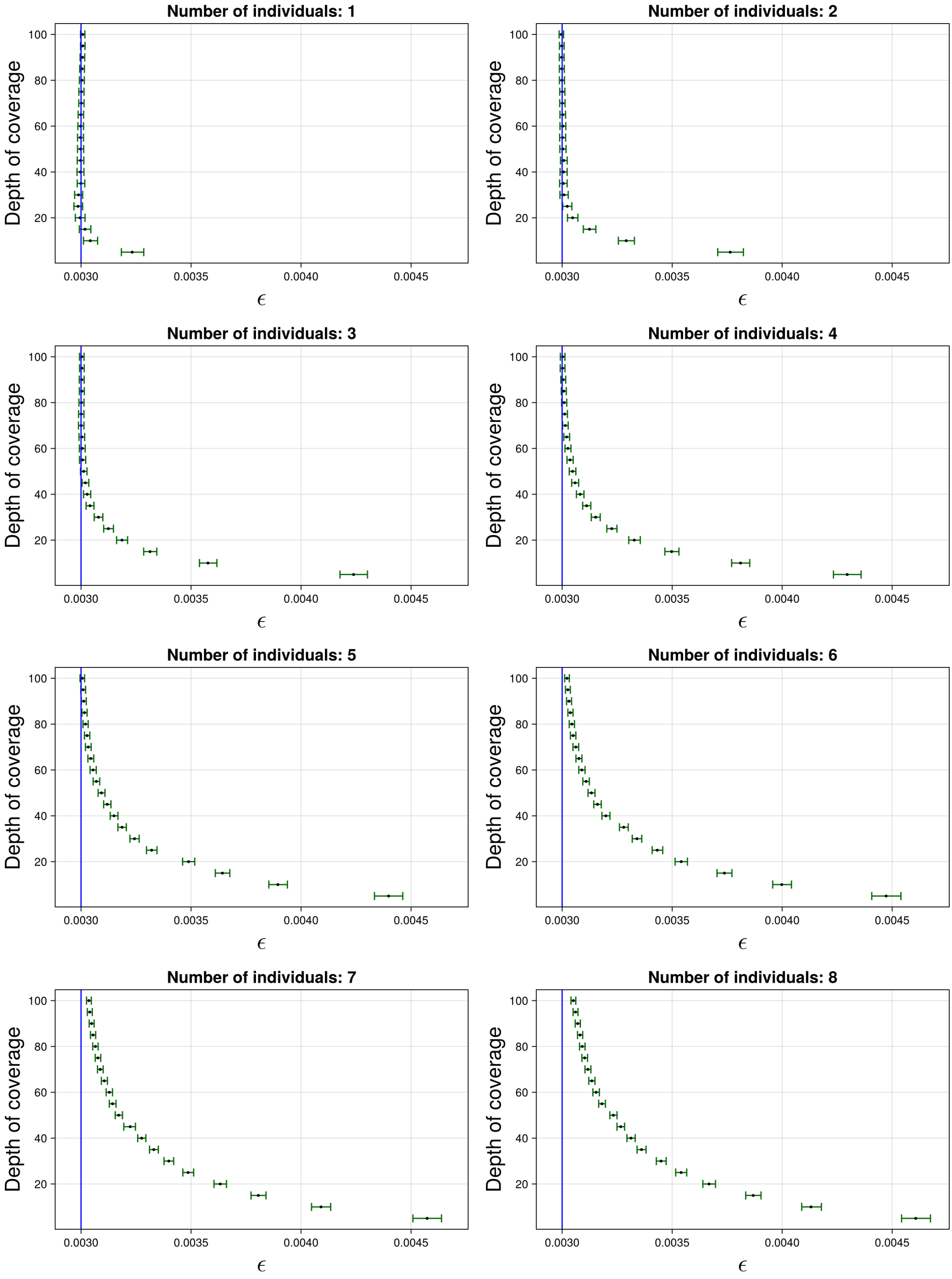

Figure S52: The 95% credible intervals for $\varepsilon$ with ancient population CEUS and anchor population CHB. The black dot indicates the median. The blue line indicates the true simulated value.

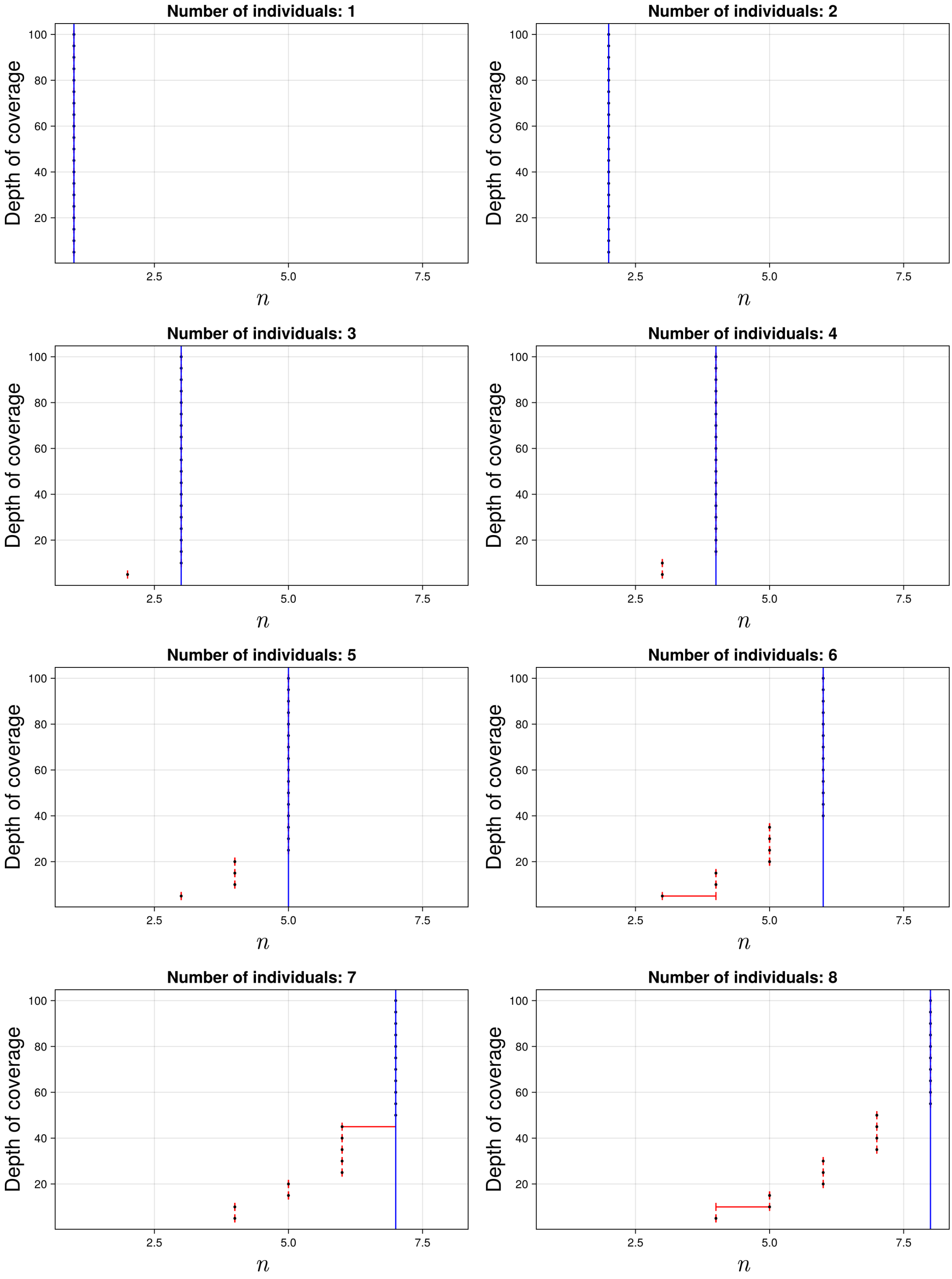

Figure S53: The 95% credible intervals for $n$ with ancient population CEUS and anchor population CHB. The black dot indicates the median. The blue line indicates the true simulated value.

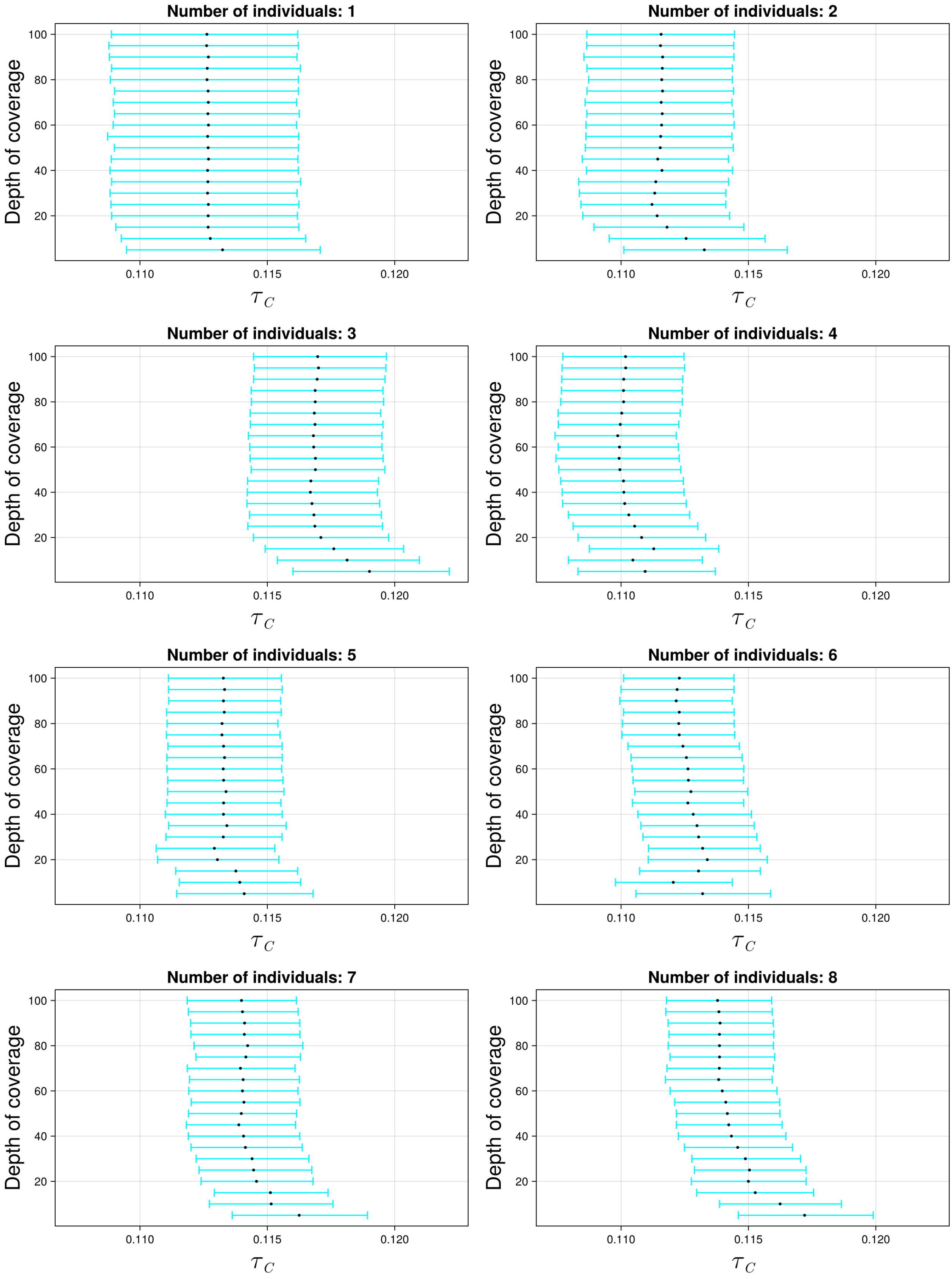

Figure S54: The 95% credible intervals for $\tau_C$ with ancient population CEUS and anchor population KAR. The black dot indicates the median.

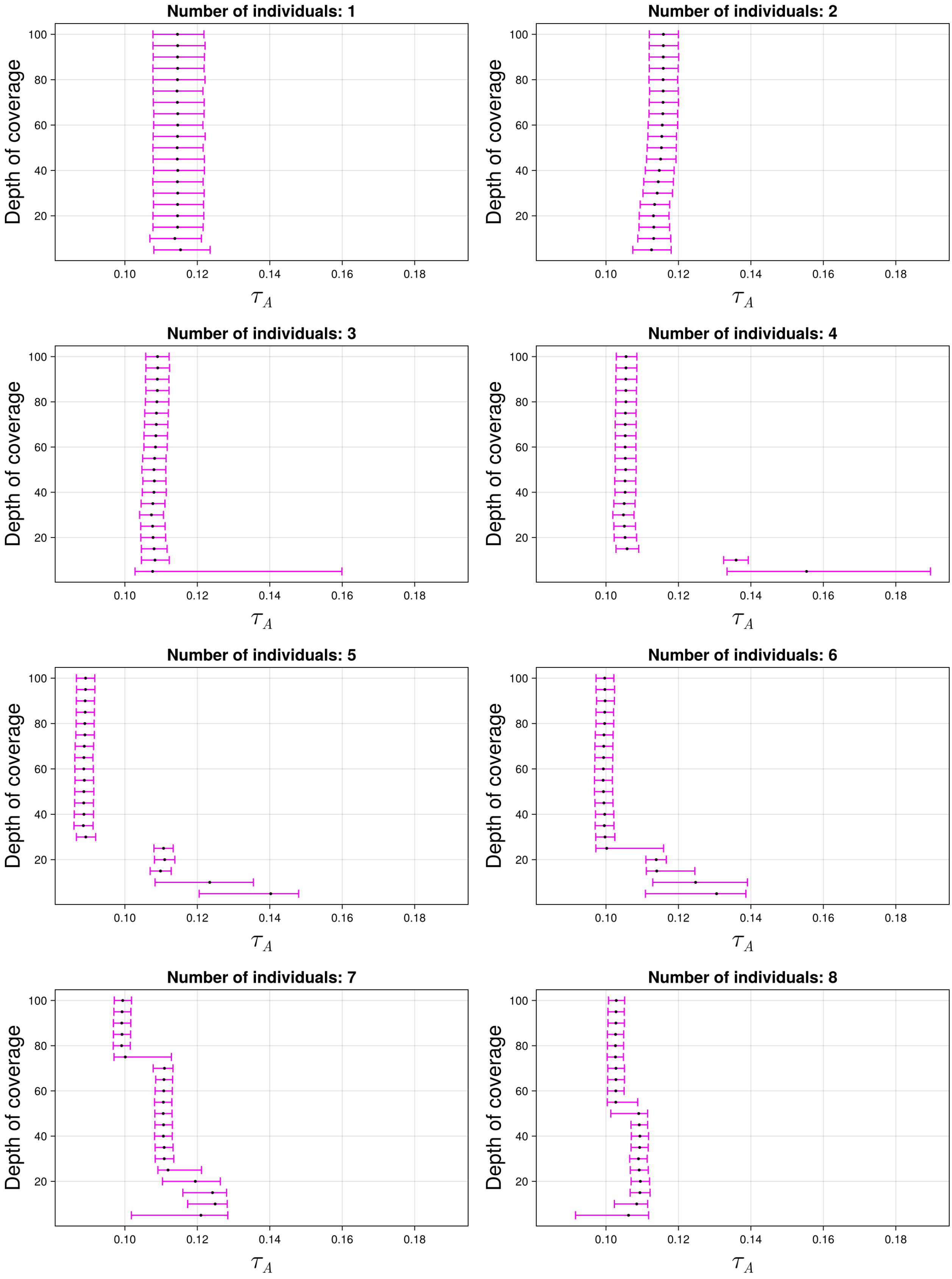

Figure S55: The 95% credible intervals for $\tau_A$ with ancient population CEUS and anchor population KAR. The black dot indicates the median.

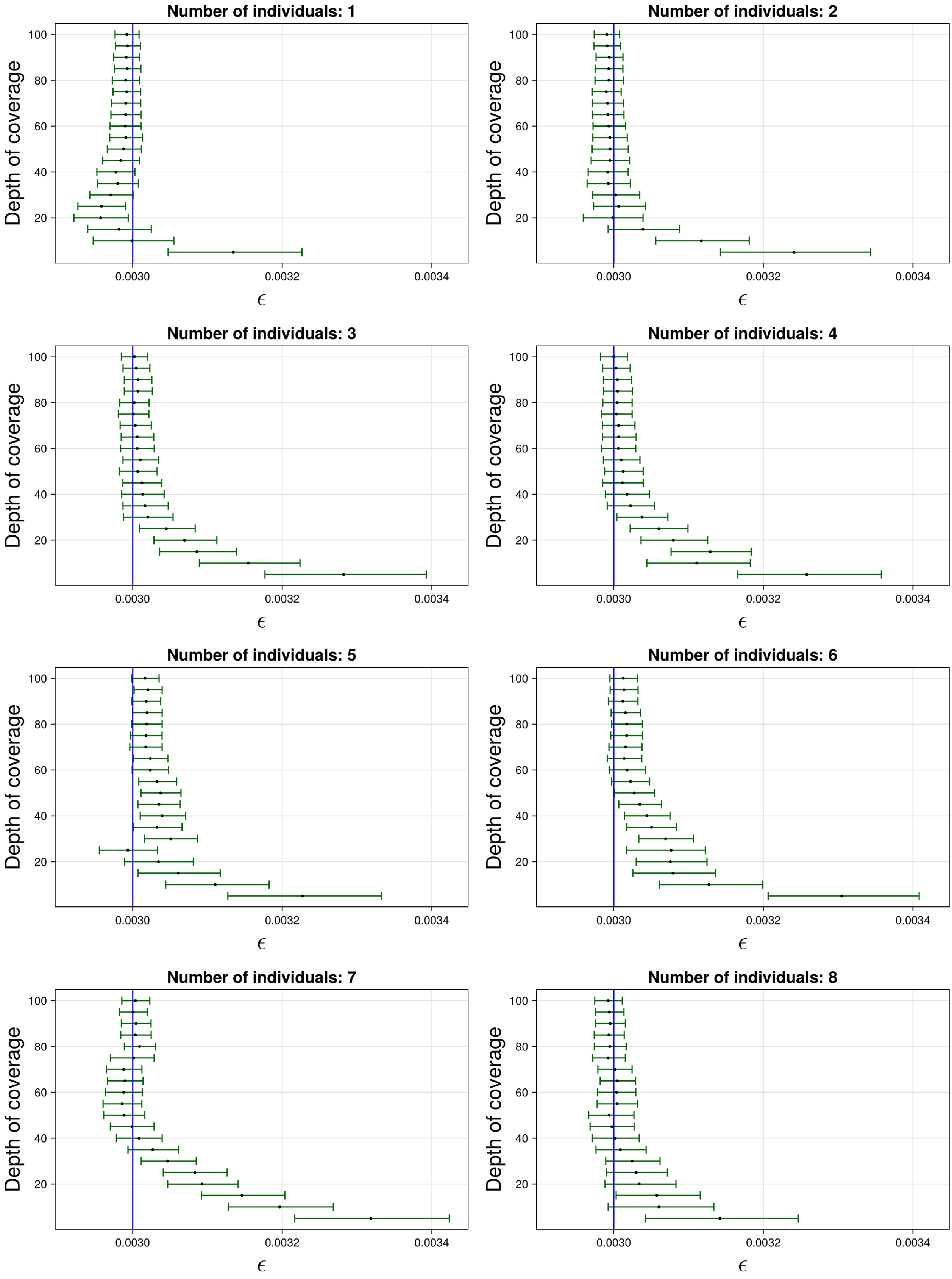

Figure S56: The 95% credible intervals for $\varepsilon$ with ancient population CEUS and anchor population KAR. The black dot indicates the median. The blue line indicates the true simulated value.

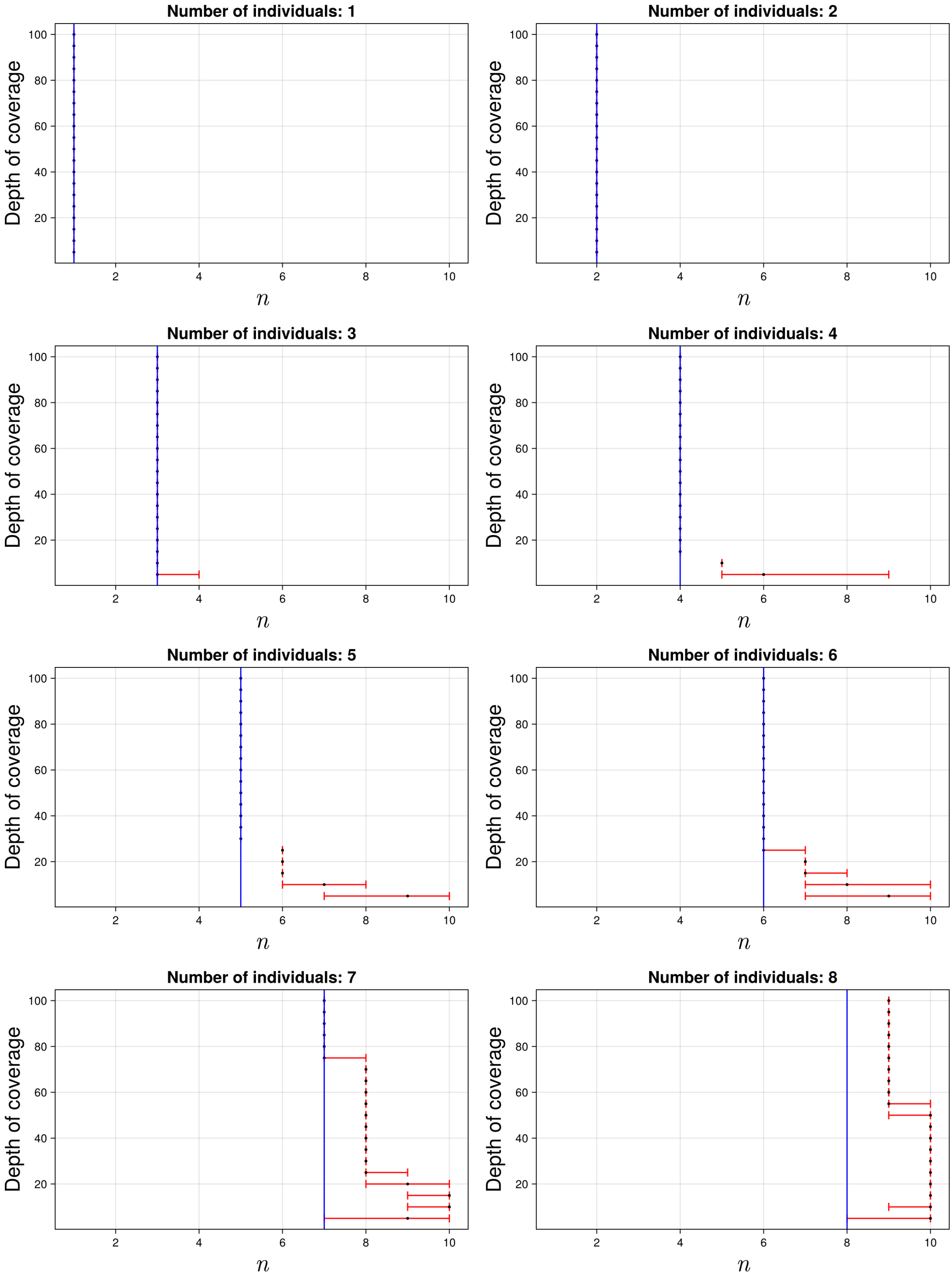

Figure S57: The 95% credible intervals for $n$ with ancient population CEUS and anchor population KAR. The black dot indicates the median. The blue line indicates the true simulated value.

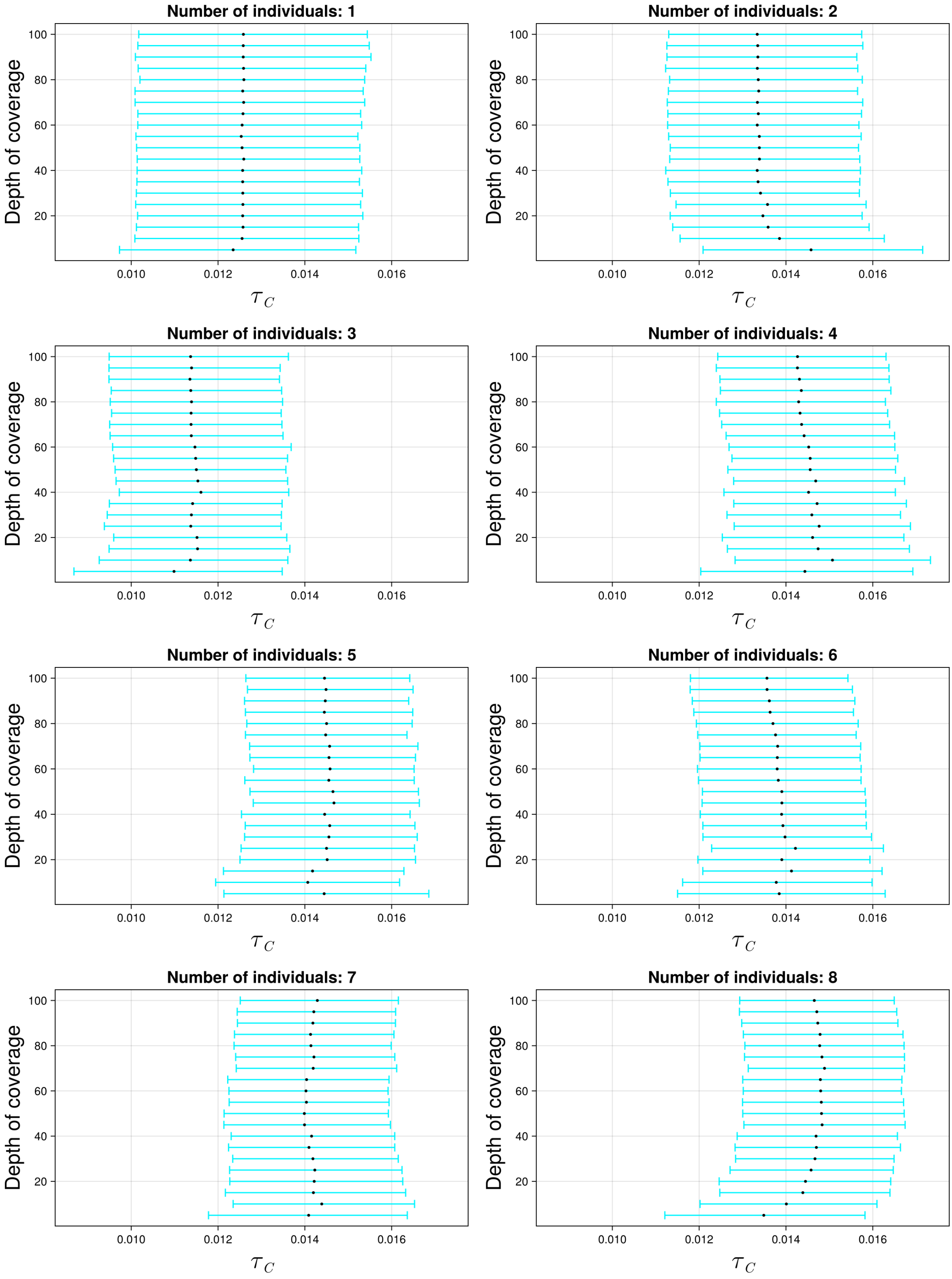

Figure S58: The 95% credible intervals for $\tau_C$ with ancient population KARS and anchor population YRI. The black dot indicates the median.

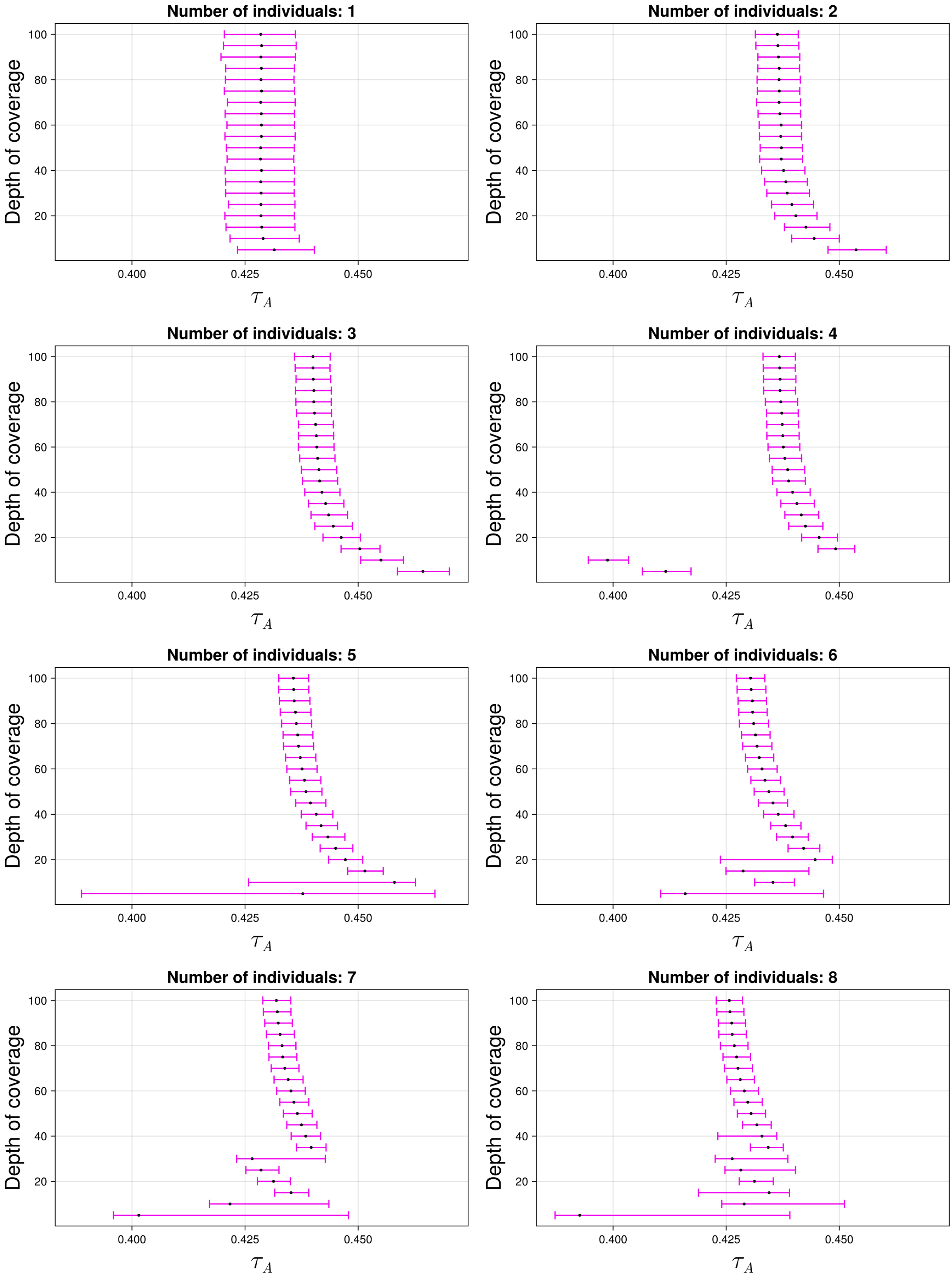

Figure S59: The 95% credible intervals for $\tau_A$ with ancient population KARS and anchor population YRI. The black dot indicates the median.

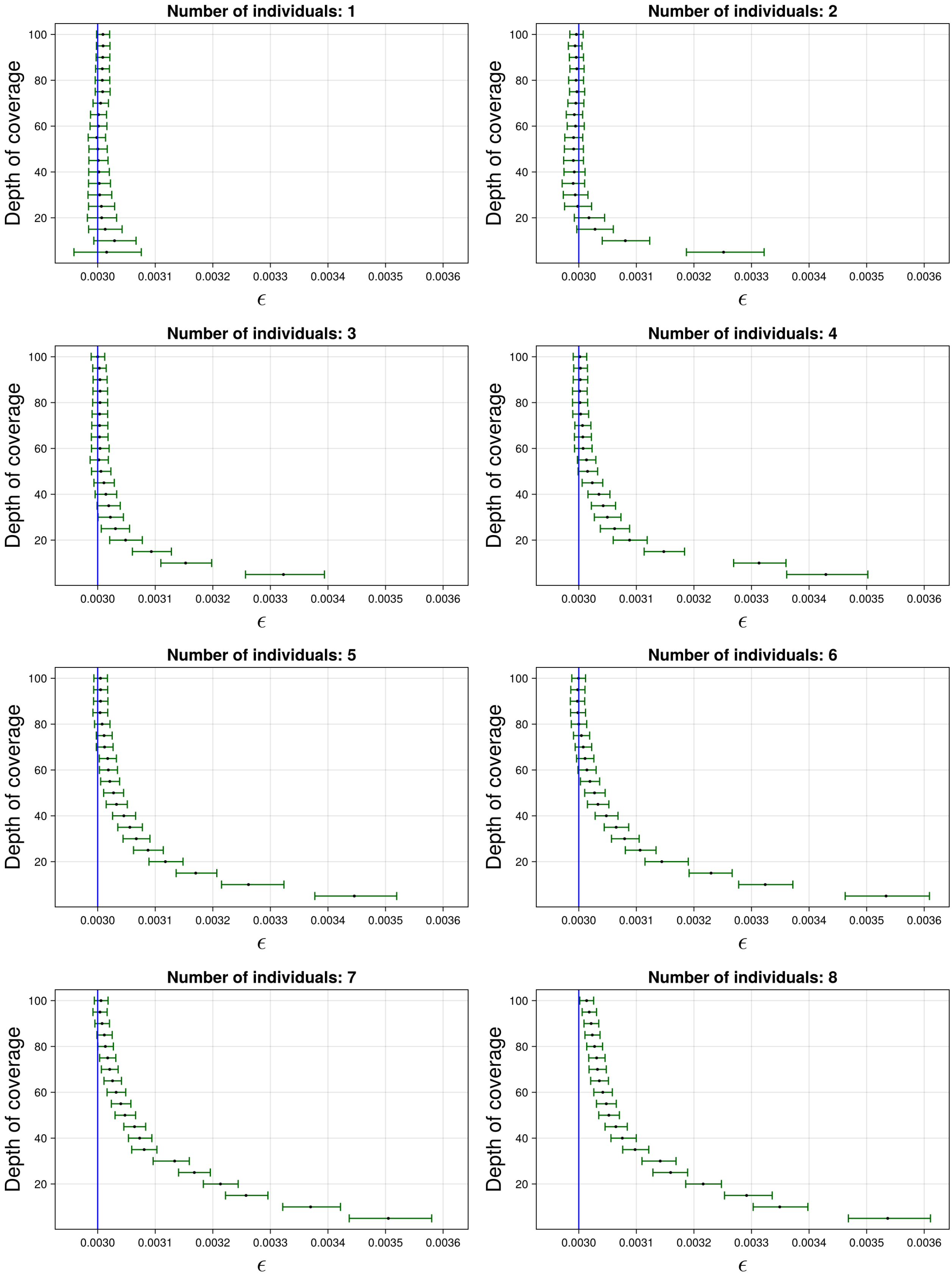

Figure S60: The 95% credible intervals for $\varepsilon$ with ancient population KARS and anchor population YRI. The black dot indicates the median. The blue line indicates the true simulated value.

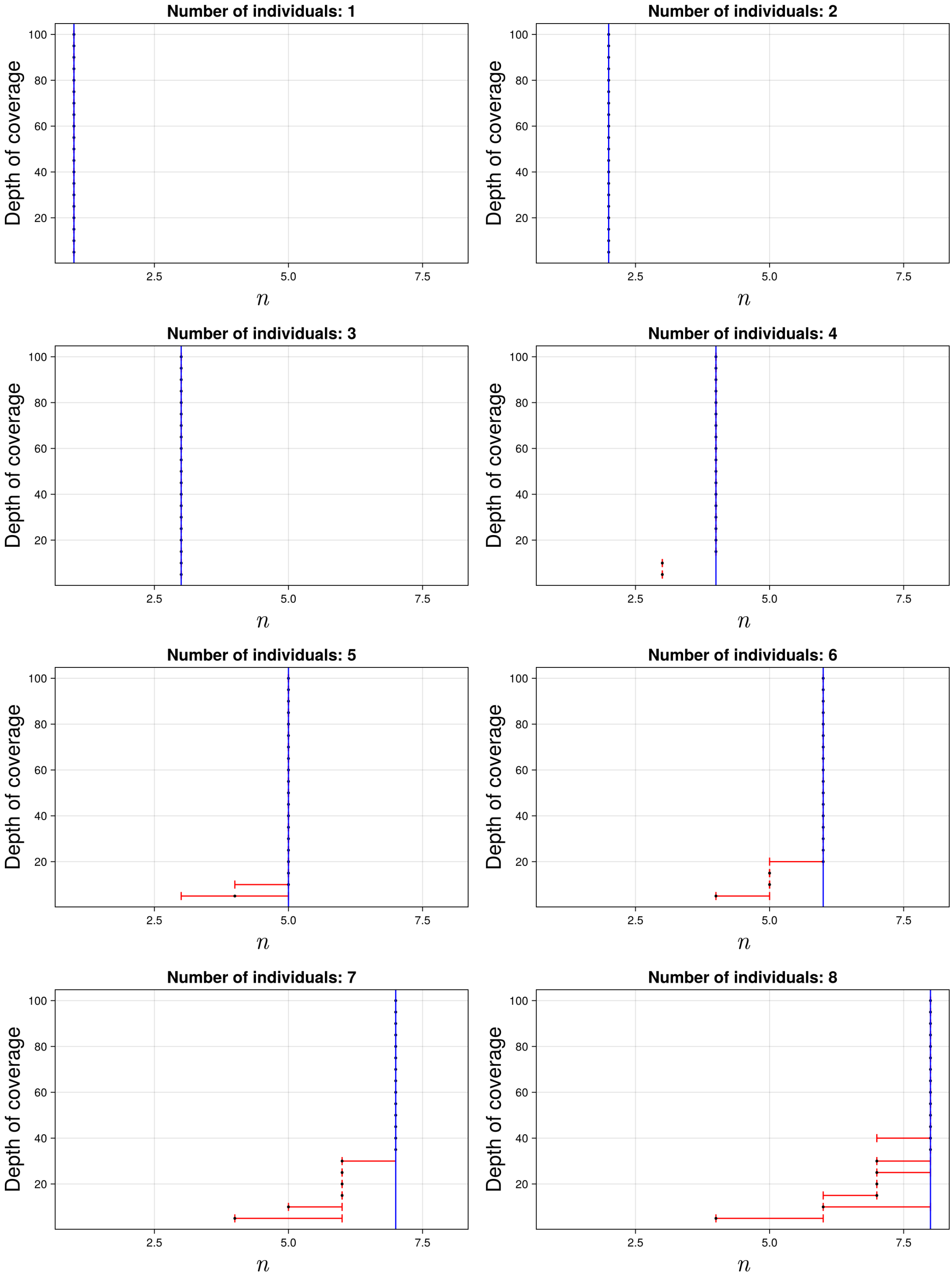

Figure S61: The 95% credible intervals for $n$ with ancient population KARS and anchor population YRI. The black dot indicates the median. The blue line indicates the true simulated value.

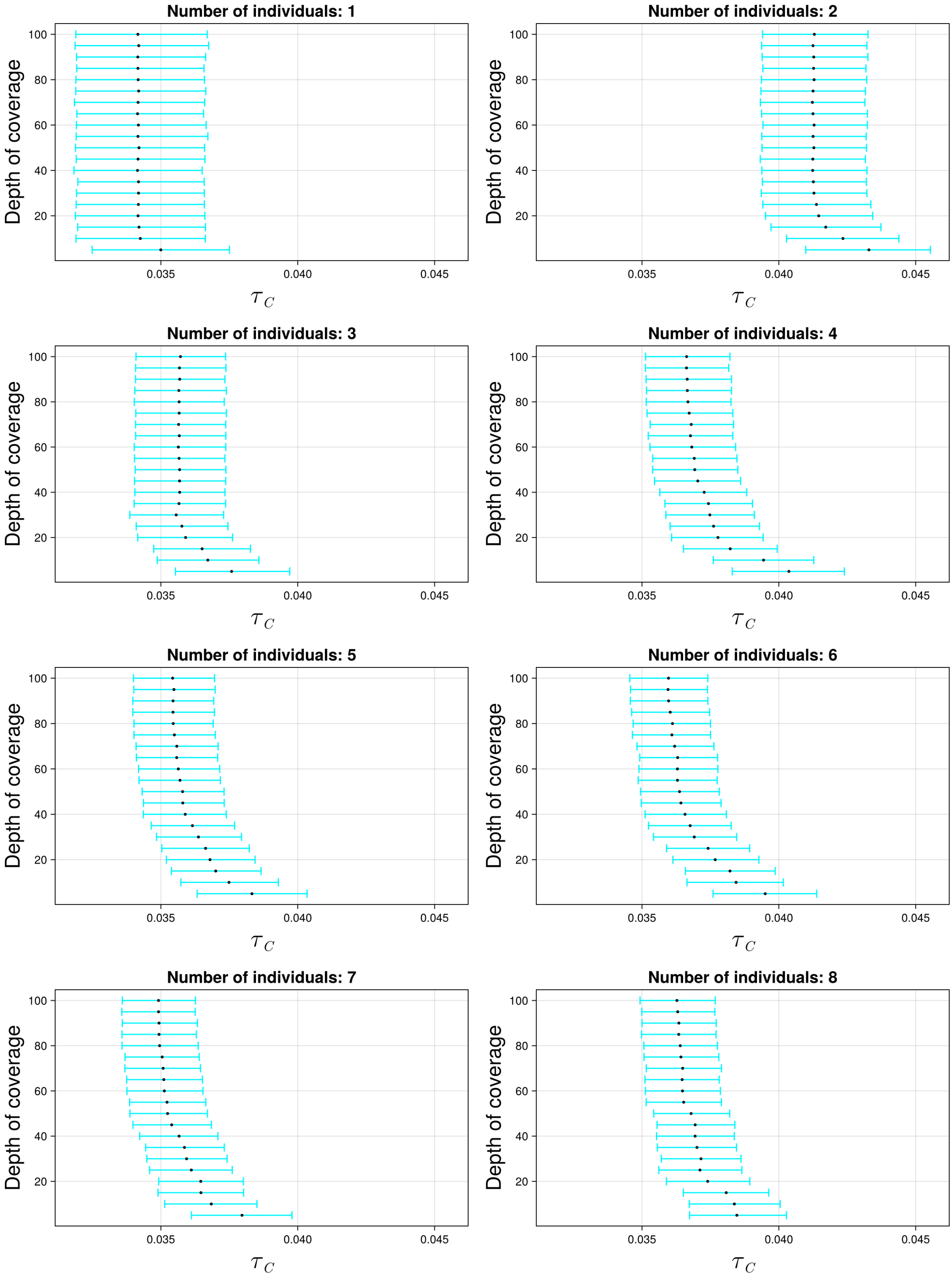

Figure S62: The 95% credible intervals for $\tau_C$ with ancient population KARS and anchor population CEU. The black dot indicates the median.

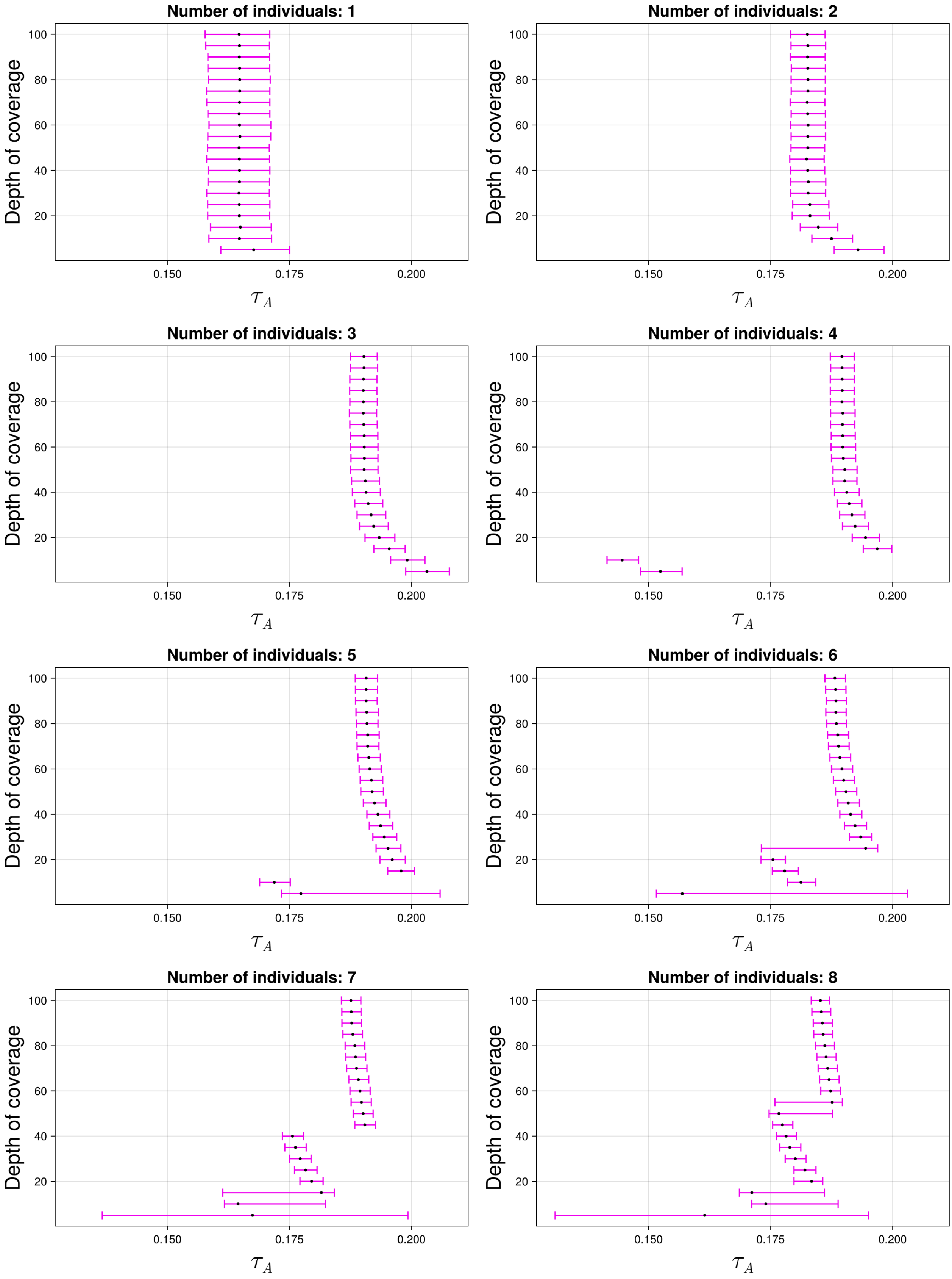

Figure S63: The 95% credible intervals for $\tau_A$ with ancient population KARS and anchor population CEU. The black dot indicates the median.

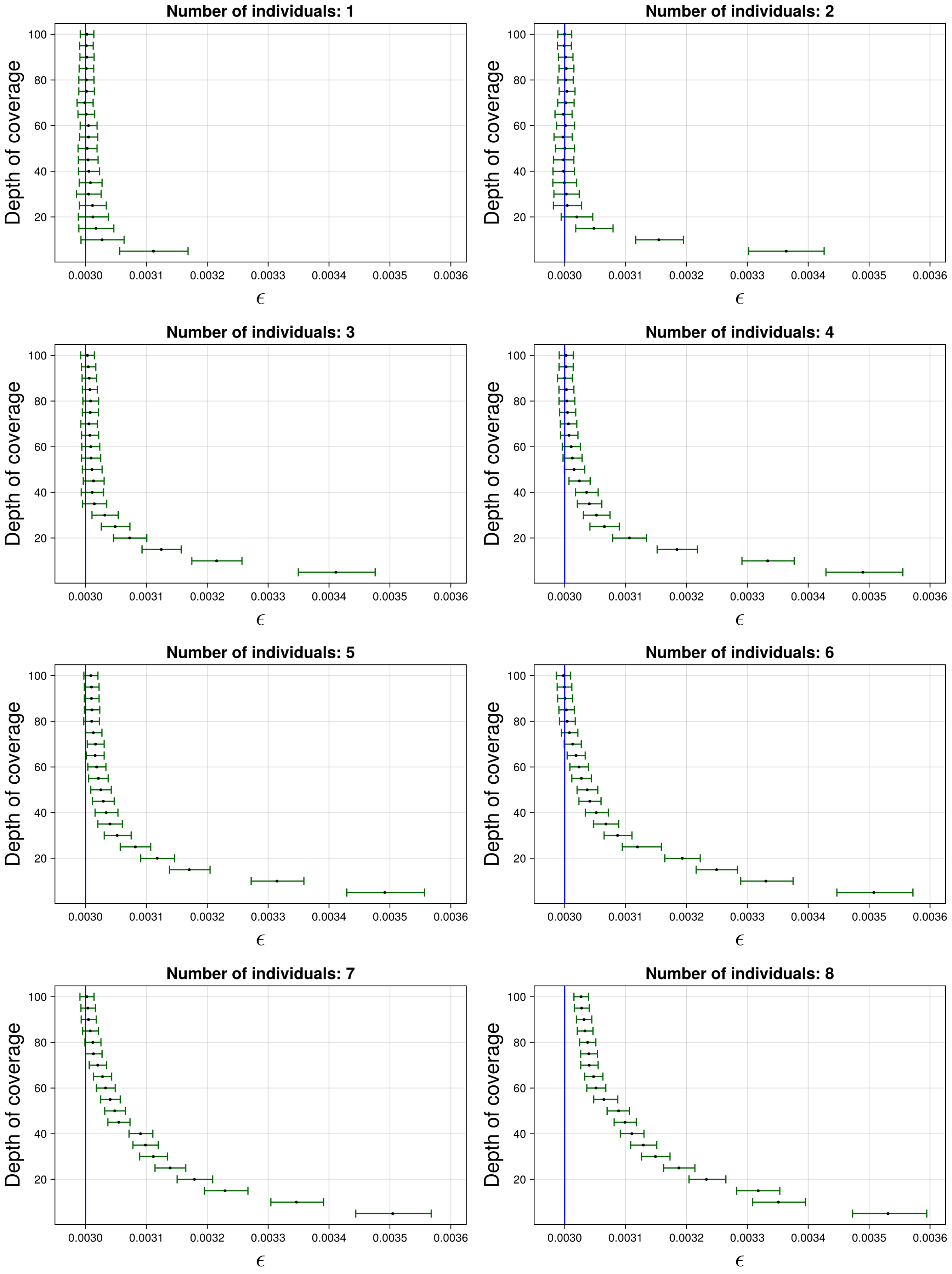

Figure S64: The 95% credible intervals for $\varepsilon$ with ancient population KARS and anchor population CEU. The black dot indicates the median. The blue line indicates the true simulated value.

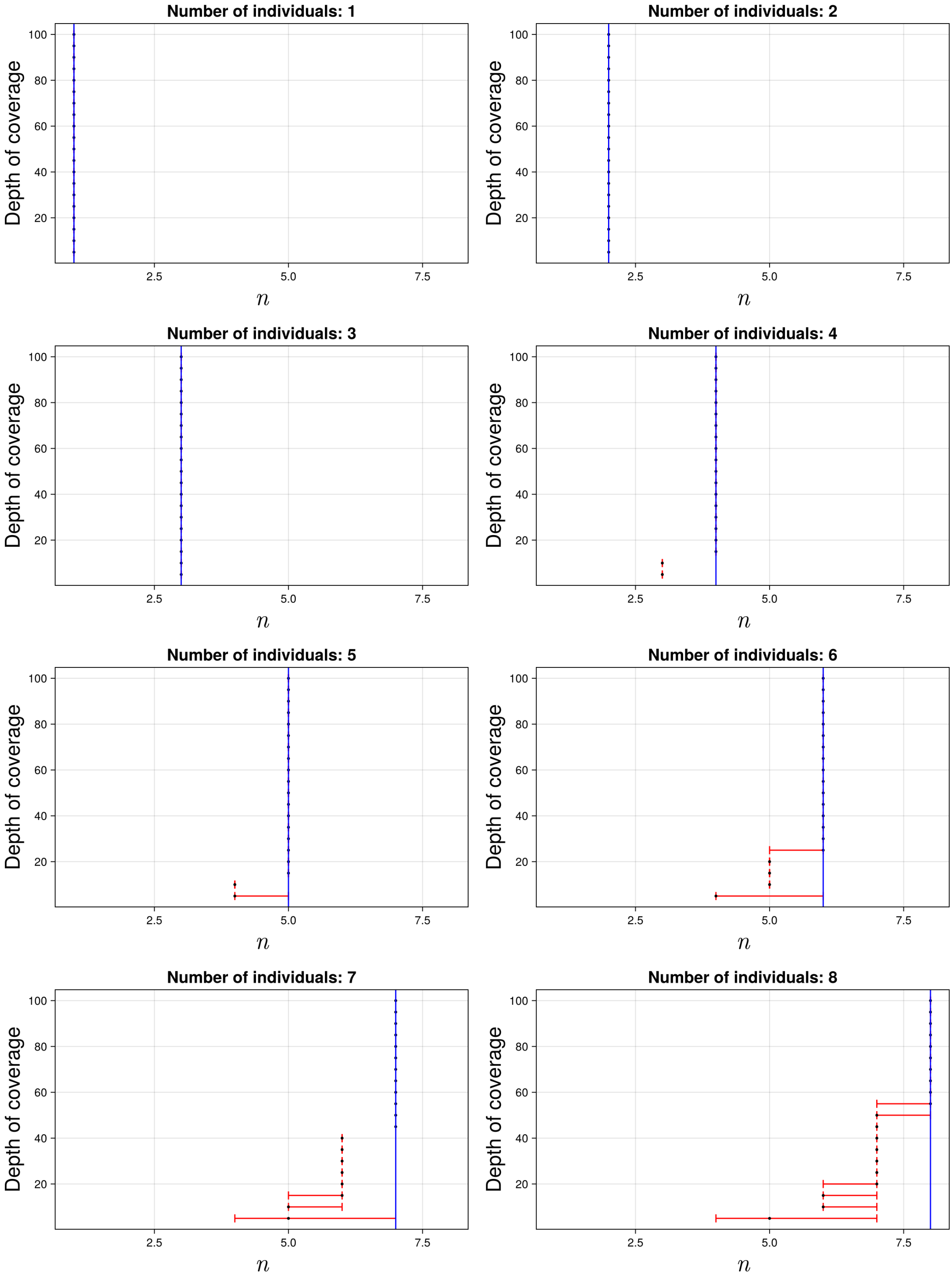

Figure S65: The 95% credible intervals for $n$ with ancient population KARS and anchor population CEU. The black dot indicates the median. The blue line indicates the true simulated value.

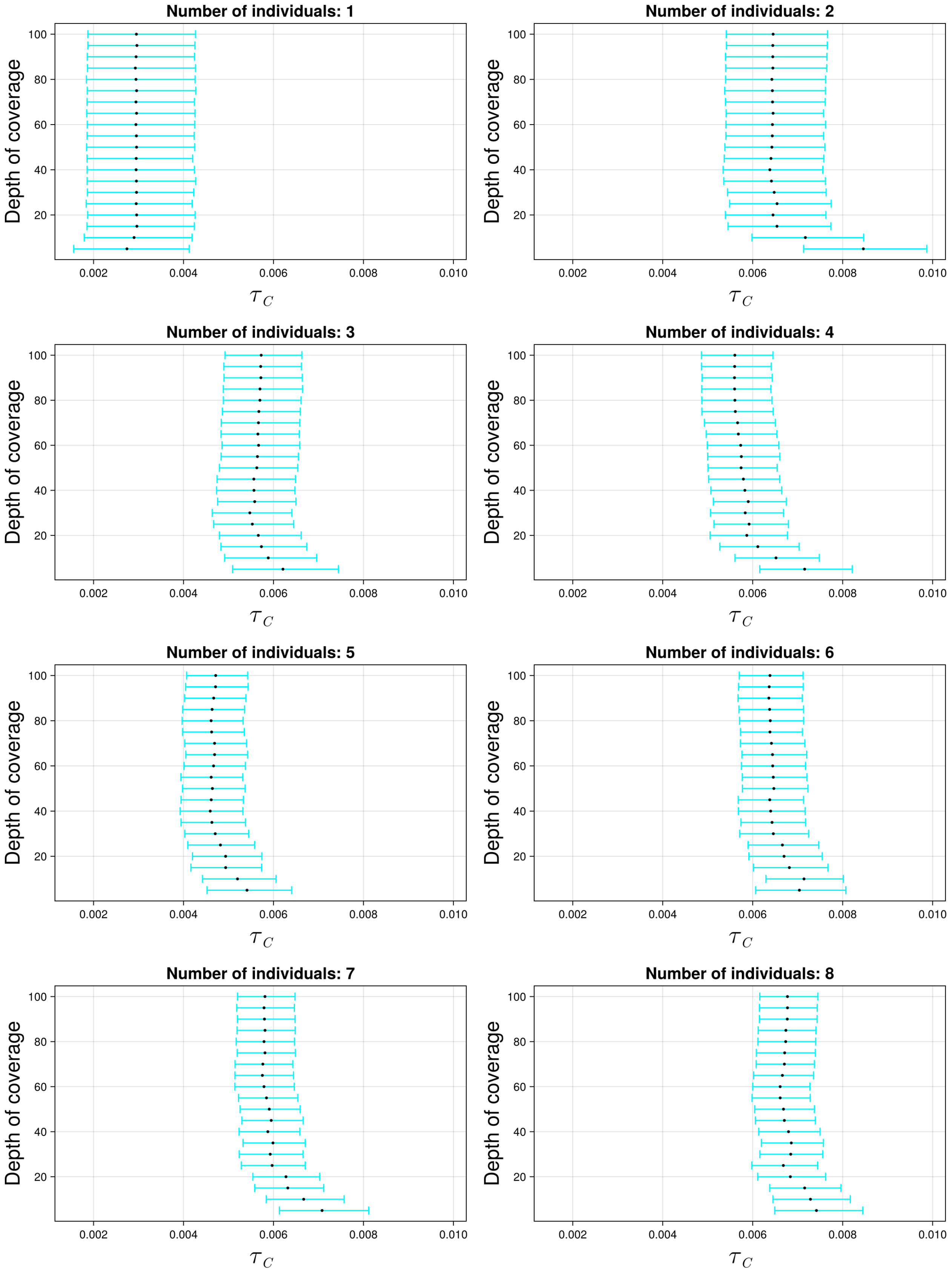

Figure S66: The 95% credible intervals for $\tau_C$ with ancient population KARS and anchor population CHB. The black dot indicates the median.

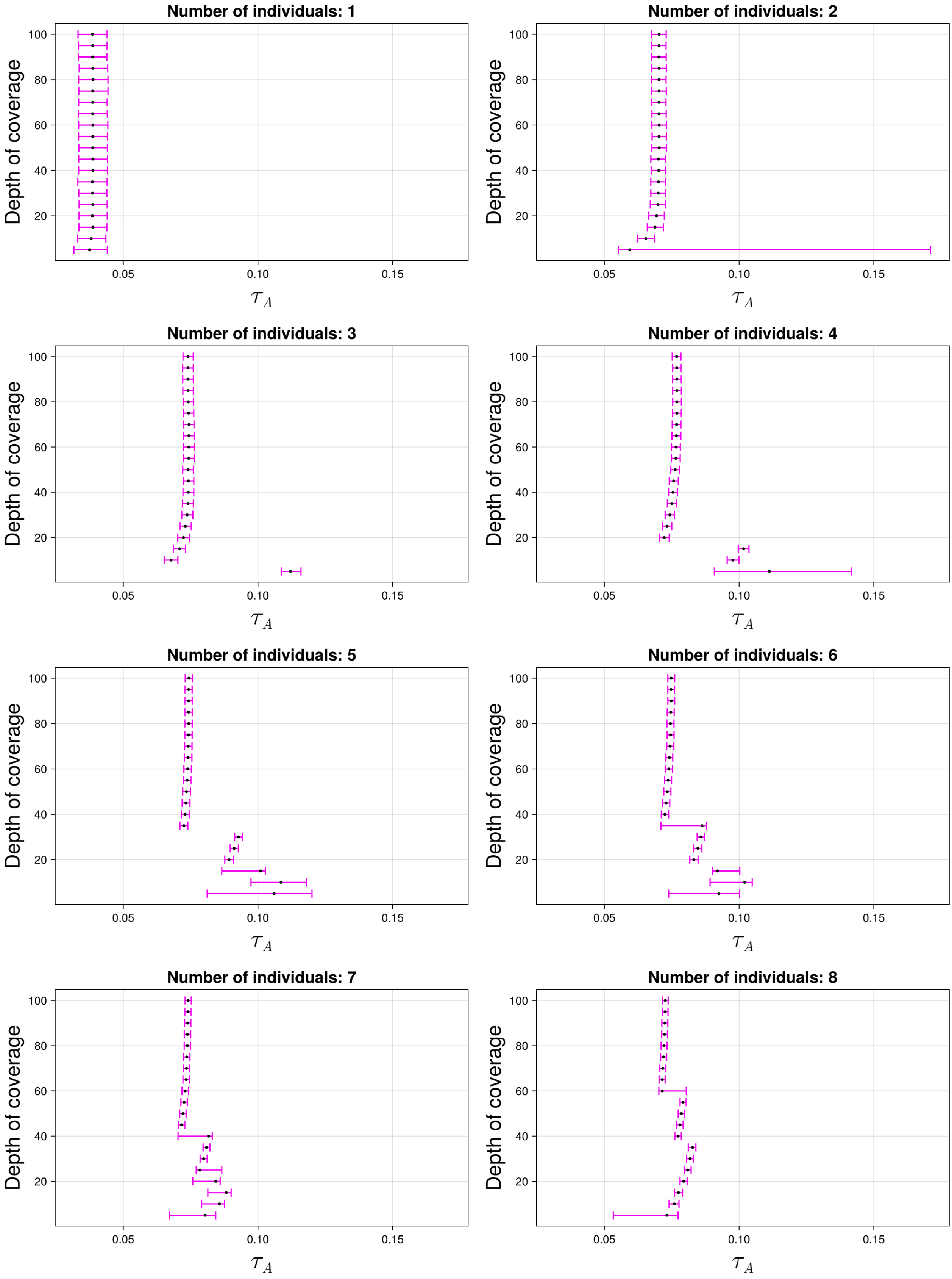

Figure S67: The 95% credible intervals for $\tau_A$ with ancient population KARS and anchor population CHB. The black dot indicates the median.

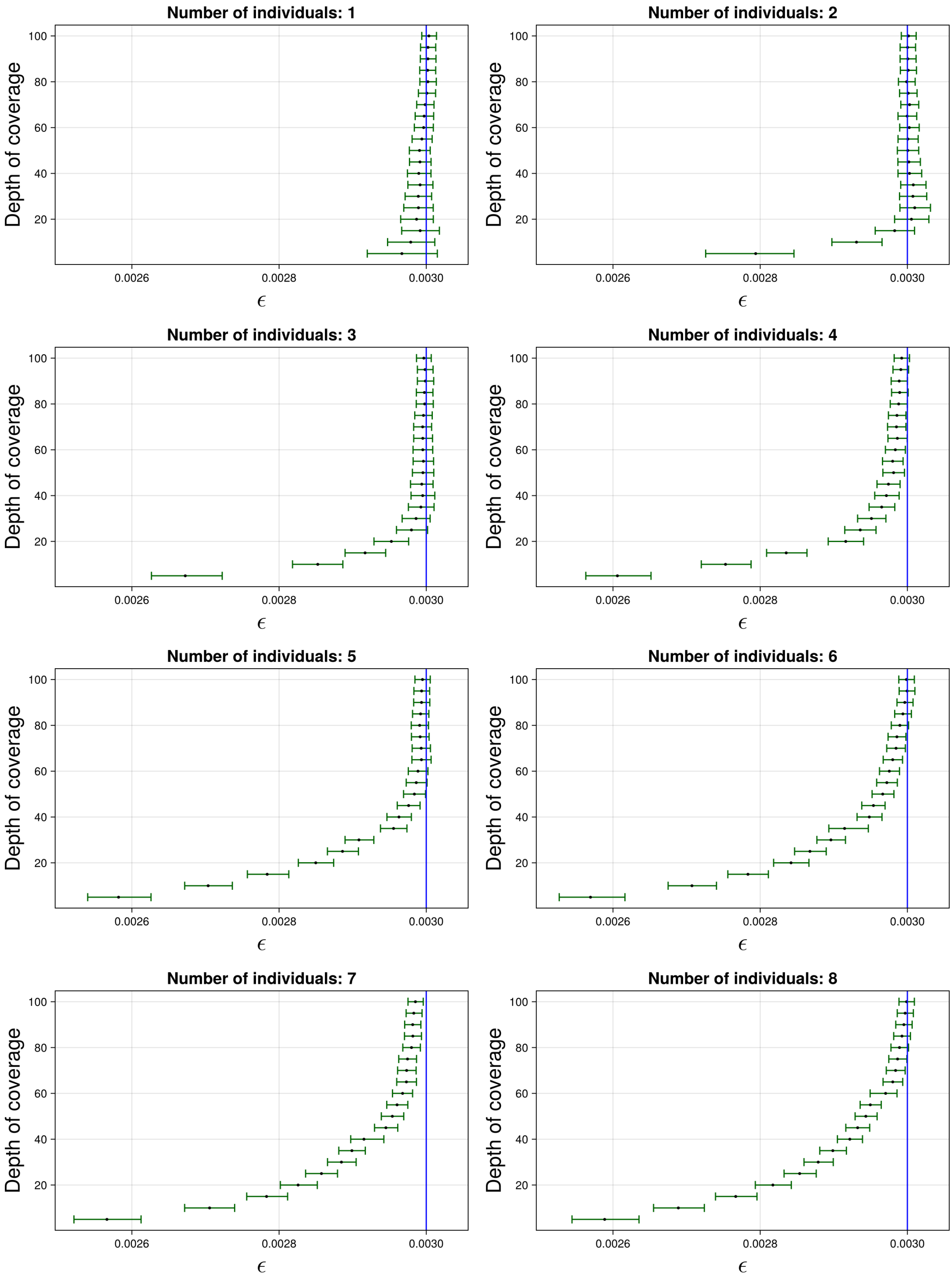

Figure S68: The 95% credible intervals for $\varepsilon$ with ancient population KARS and anchor population CHB. The black dot indicates the median. The blue line indicates the true simulated value.

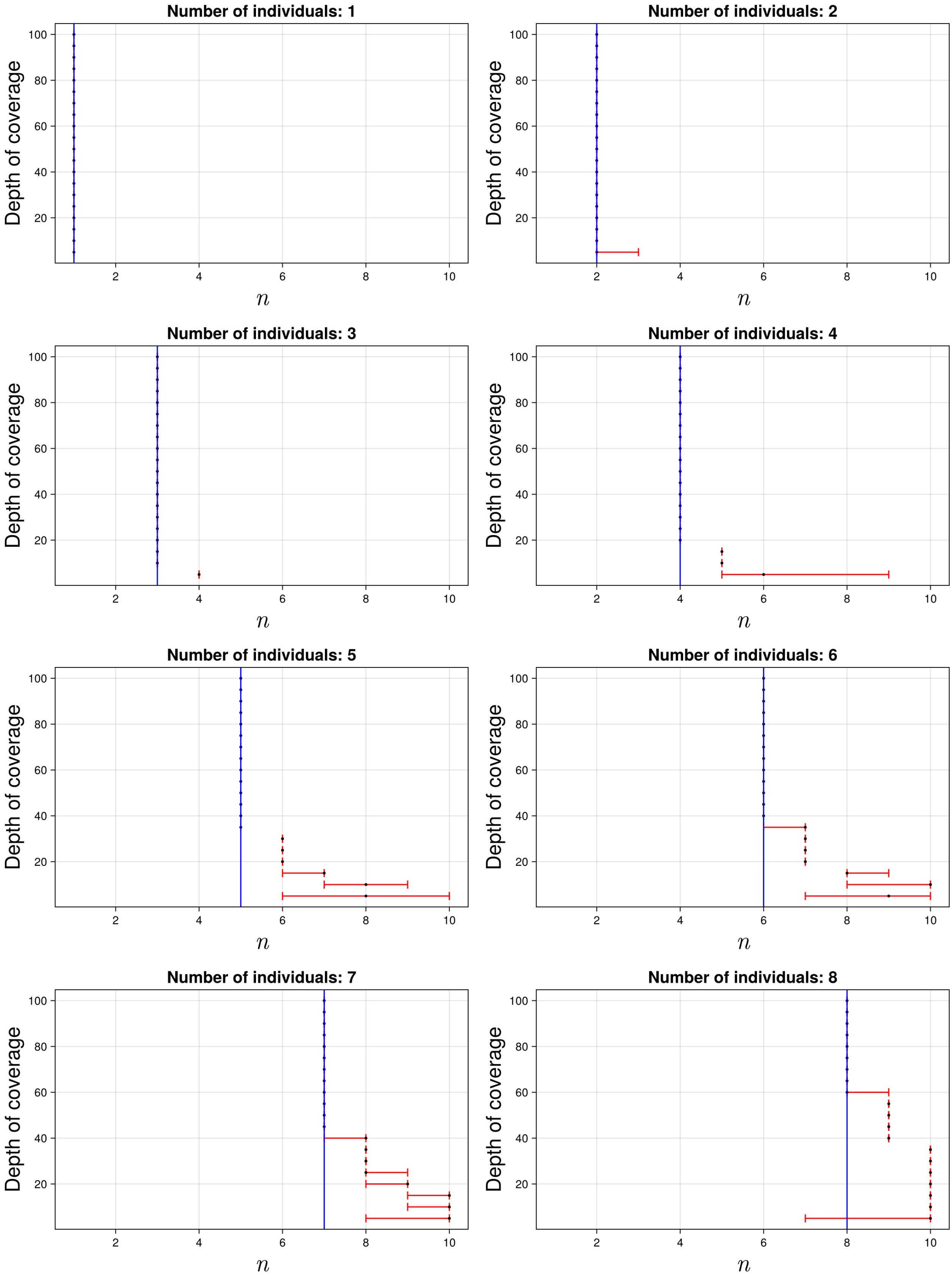

Figure S69: The 95% credible intervals for $n$ with ancient population KARS and anchor population CHB. The black dot indicates the median. The blue line indicates the true simulated value.

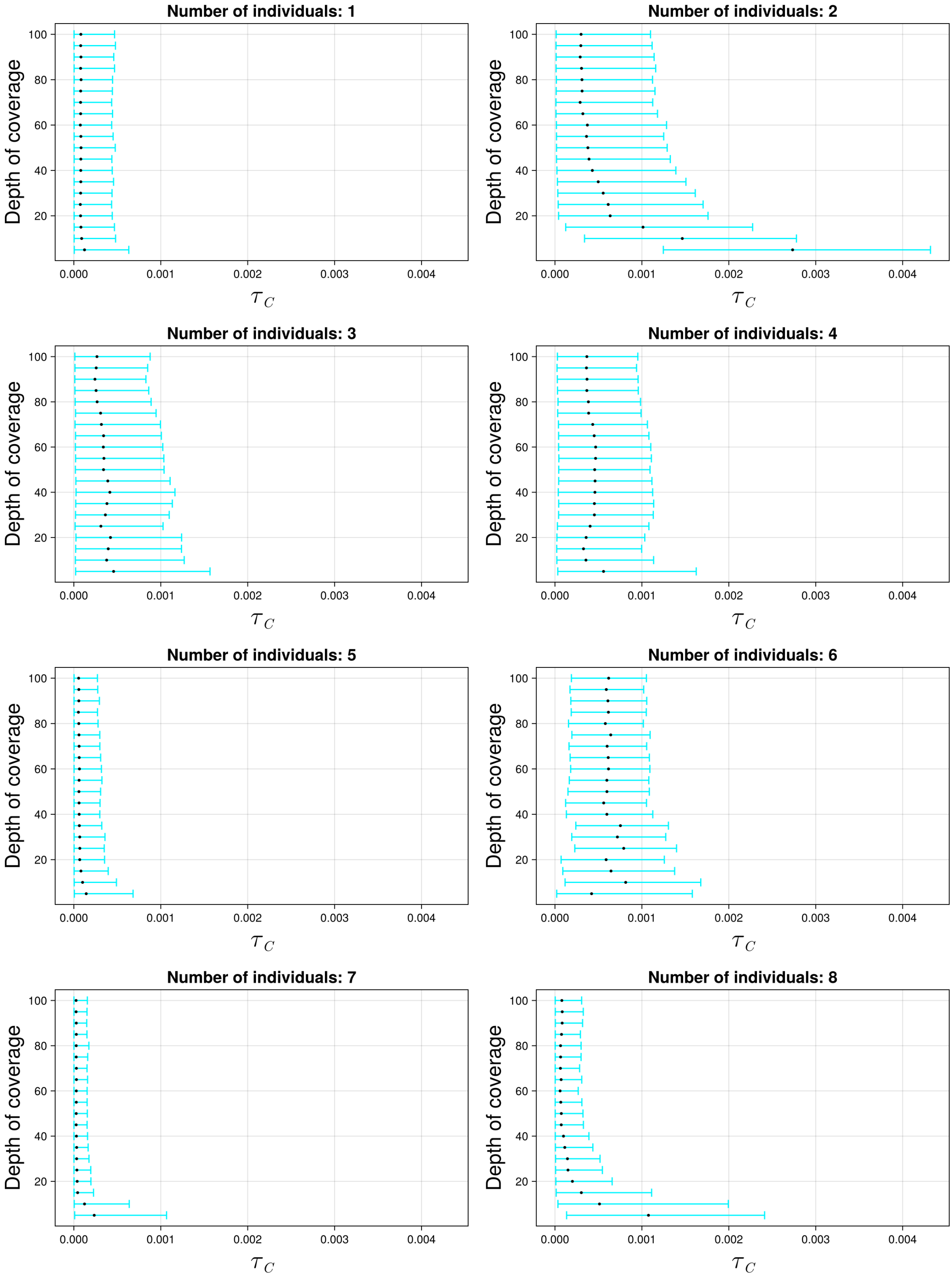

Figure S70: The 95% credible intervals for $\tau_C$ with ancient population KARS and anchor population KAR. The black dot indicates the median.

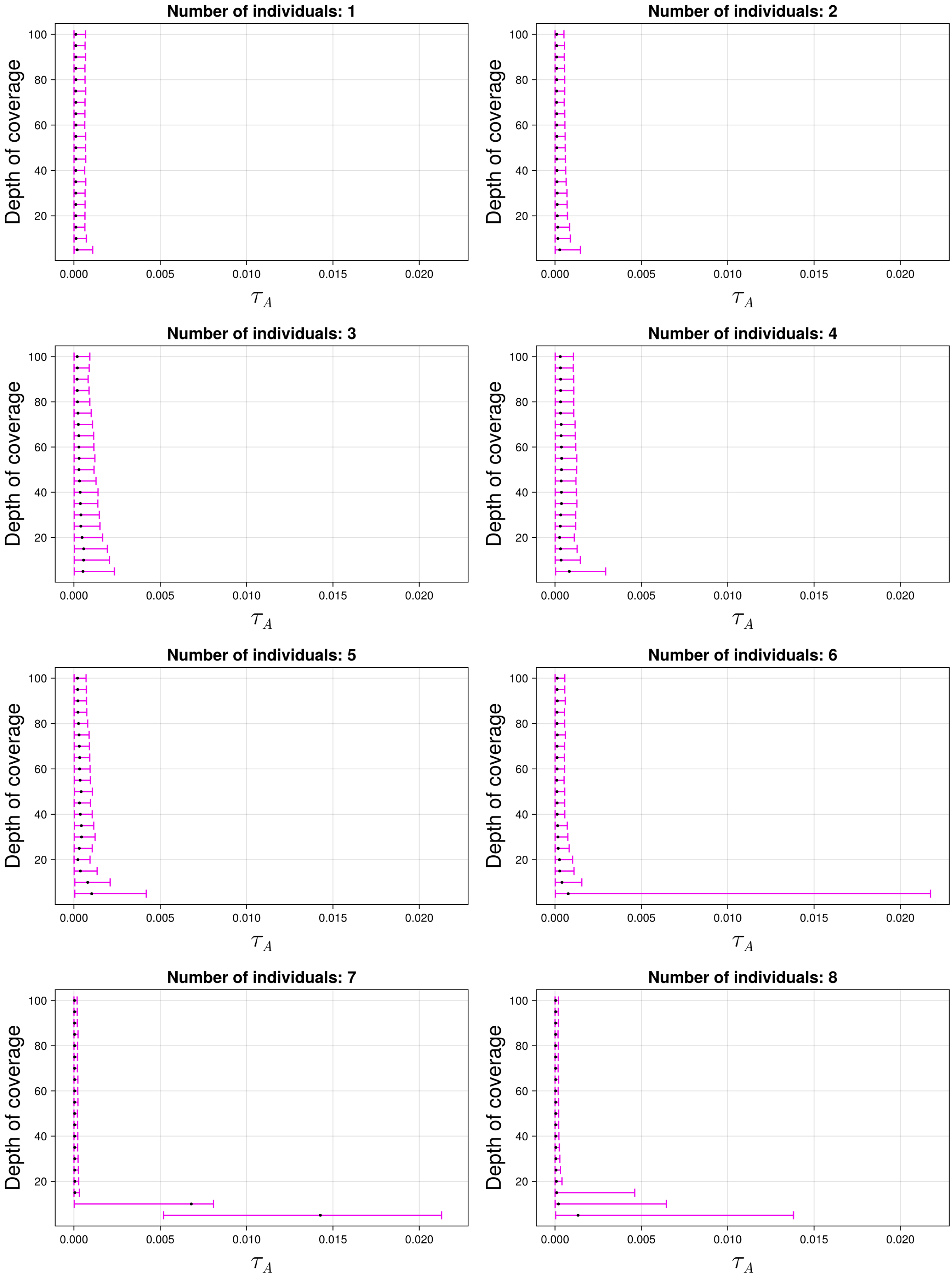

Figure S71: The 95% credible intervals for $\tau_A$ with ancient population KARS and anchor population KAR. The black dot indicates the median.

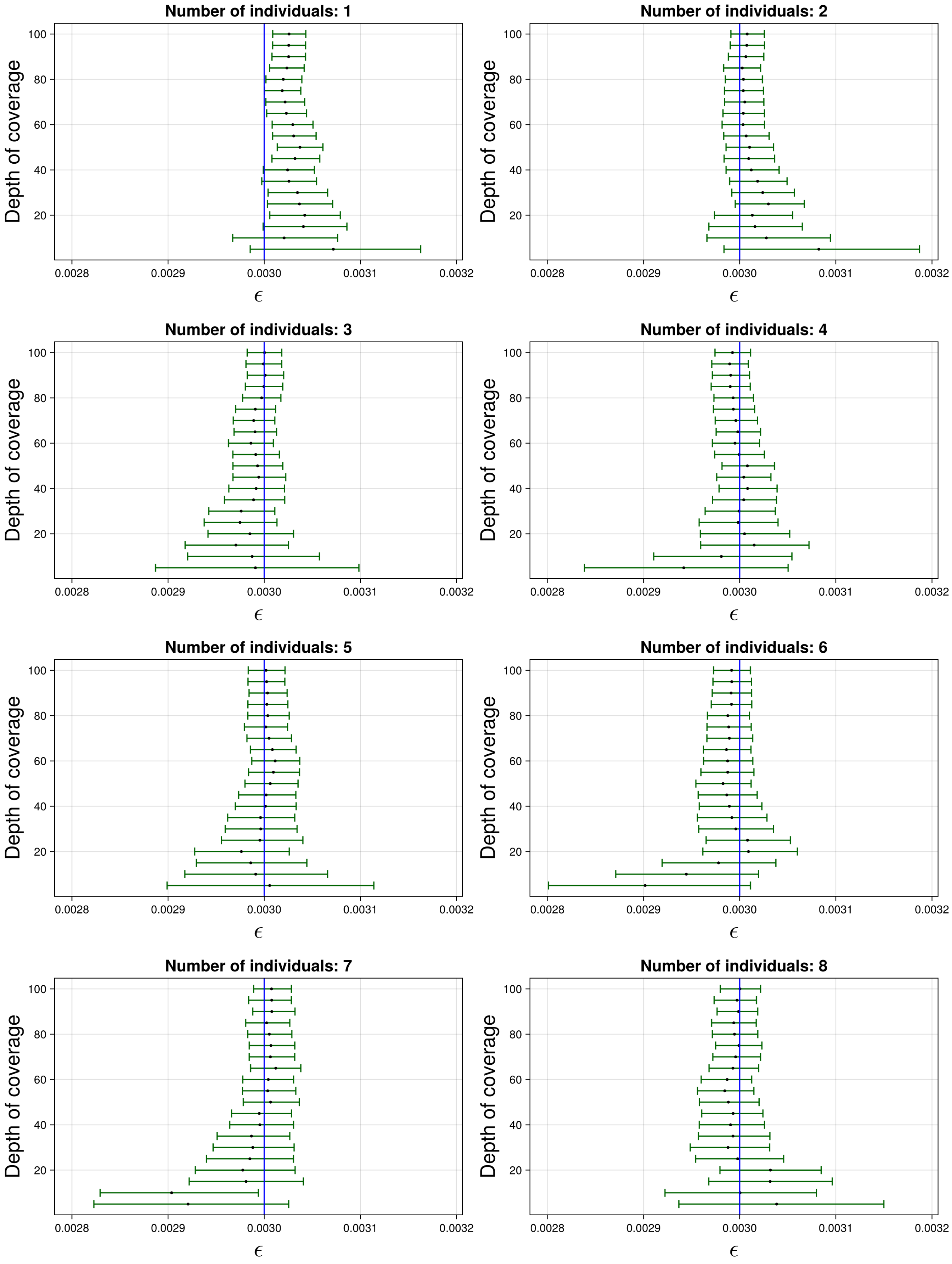

Figure S72: The 95% credible intervals for $\varepsilon$ with ancient population KARS and anchor population KAR. The black dot indicates the median. The blue line indicates the true simulated value.

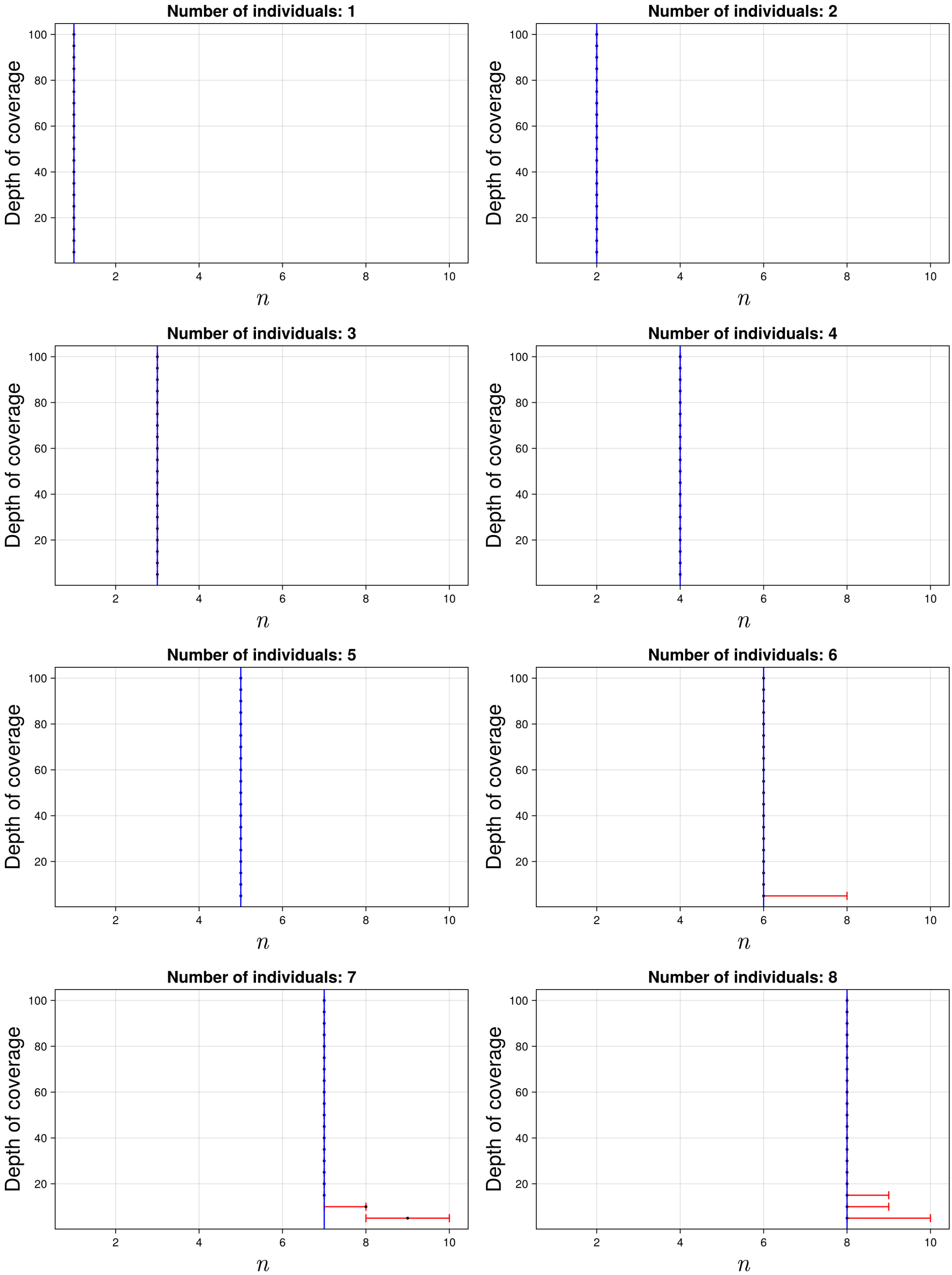

Figure S73: The 95% credible intervals for $n$ with ancient population KARS and anchor population KAR. The black dot indicates the median. The blue line indicates the true simulated value.

#### S5.4 Plot syntetic empirical data

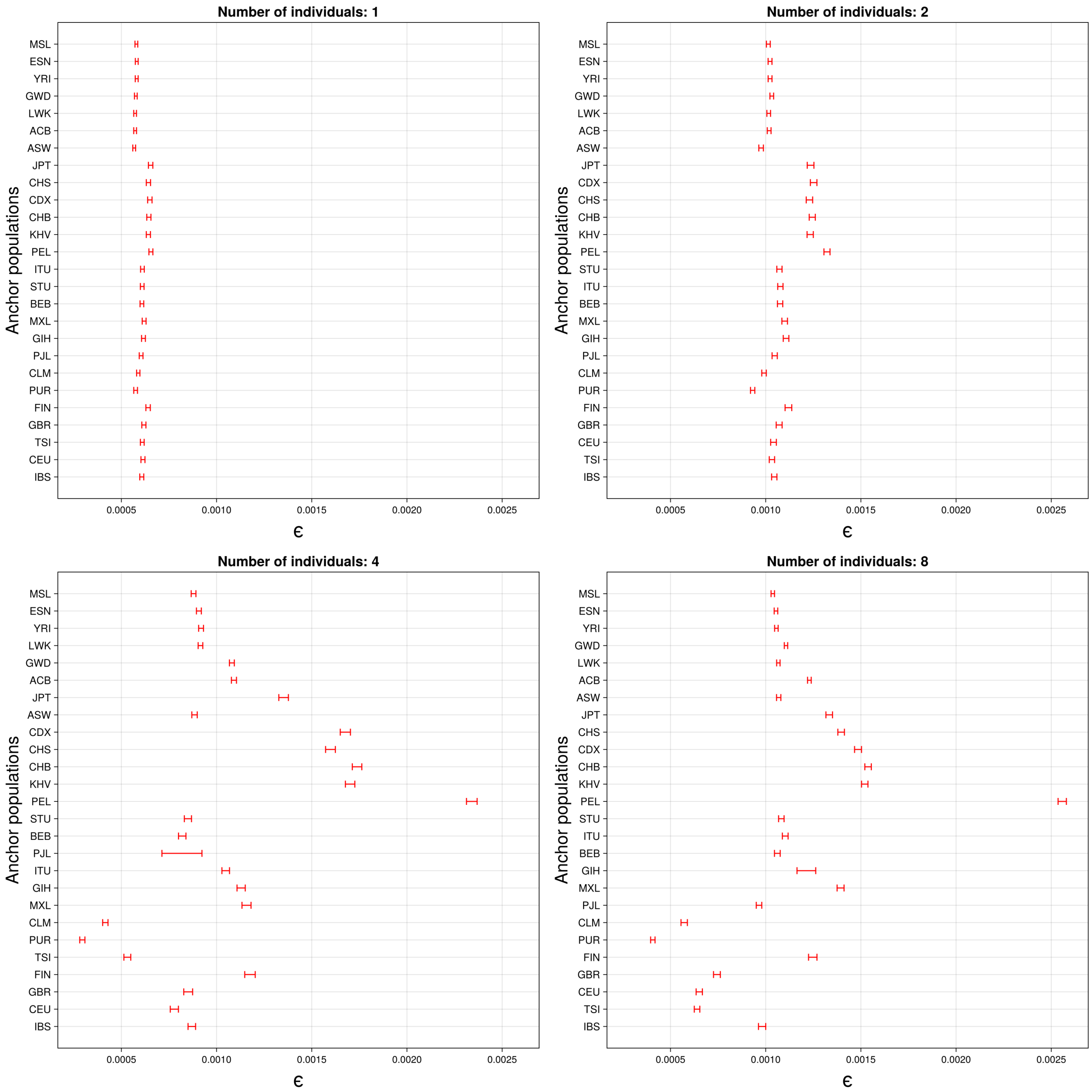

Figure S74: These are the credible intervals of the error rate $\varepsilon$. The anchor populations are ordered according to the median of $\tau_C + \tau_A$, with the lowest value at the bottom.

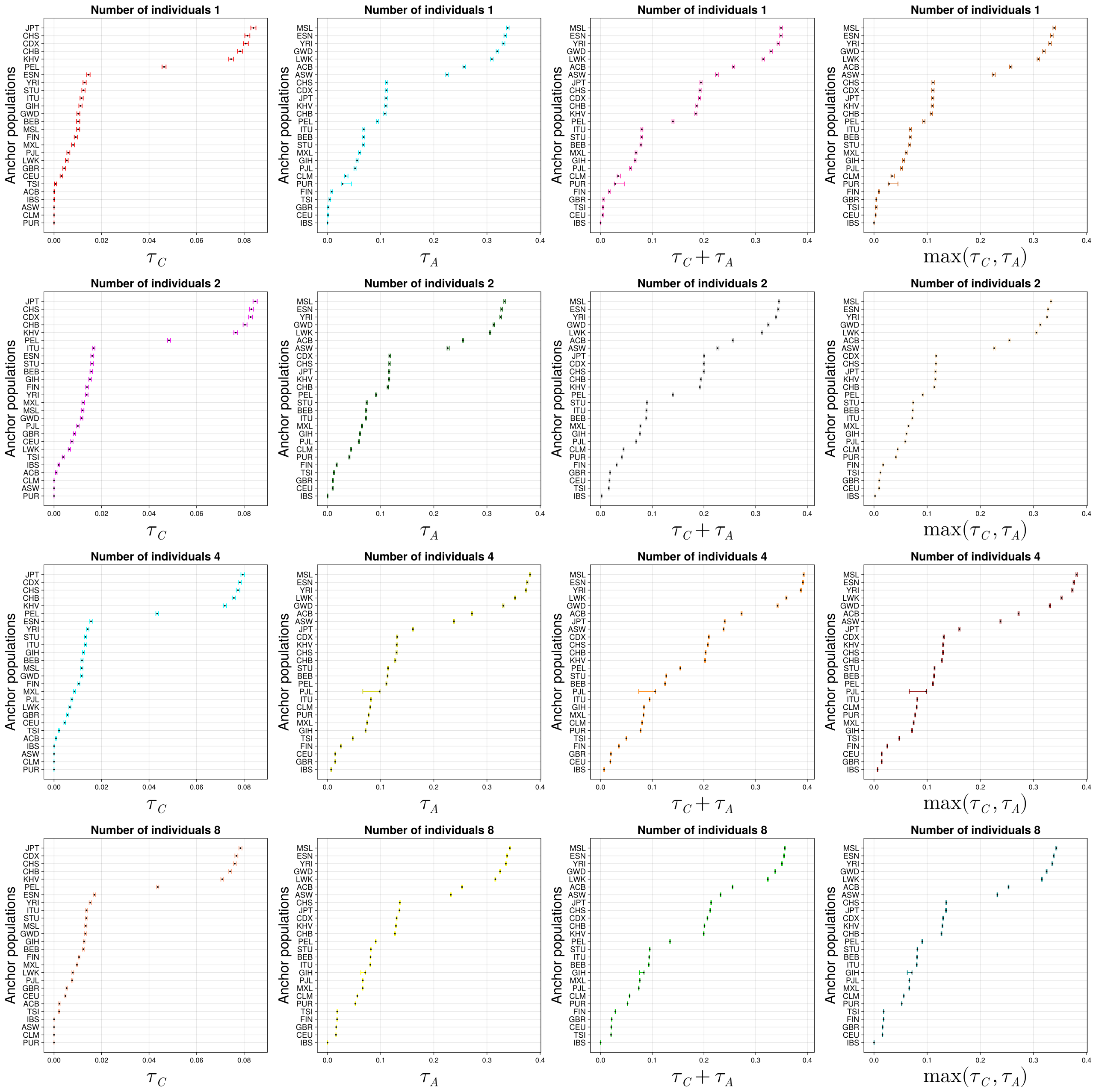

Figure S75: Populations ordered by $\tau_C$, $\tau_A$, $\tau_C + \tau_A$ and $\max(\tau_C\,,\,\tau_A)$ (with the lowest value at the bottom) for the samples simulated with IBS as source population, respectively, along with 95% credible intervals.

### S6 Code

#### S6.1 Estimating parameters

Demes file:

```
---
time_units: generations
demes:
  - name: common_ancestor
    epochs:
      - end_time: 1000
        start_size: 2000
```

```
  - name: ancient
    ancestors:
      - common_ancestor
    epochs:
      - start_size: 2000
  - name: anchor
    ancestors:
      - common_ancestor
    epochs:
      - start_size: 2000
```

Python code:

```
import sys # for command line arguments.
import msprime
import demes
import subprocess # to call gzip from within Python.

def two_demes(demesfile, demesfiledescription, random_seed=None):
    graph = demes.load(demesfile)
    demography = msprime.Demography.from_demes(graph)
    ts = msprime.sim_ancestry(samples={"ancient":5_000, "anchor":5_000},
    demography=demography, random_seed=random_seed, recombination_rate=1e-8,
    sequence_length=10e7)
    mts = msprime.sim_mutations(ts, rate=2e-8, random_seed=random_seed)
    filenamewithoutextension = "two_demes_with_demes_" +
    demesfiledescription +str(random_seed)
    mts.dump( filenamewithoutextension+".tree")
    vcffilename=filenamewithoutextension+".vcf"
    with open(vcffilename, "w") as vcf_file:
        mts.write_vcf(vcf_file)
    subprocess.call("/home/ctools/bin/bgzip -f "+vcffilename, shell=True)

if len(sys.argv) == 4:
    two_demes( sys.argv[1], sys.argv[2], int(sys.argv[3]))
elif len(sys.argv) == 3:
    two_demes(sys.argv[1], sys.argv[2])
else:
    println("Something went wrong, use one or two arguments:
    the first argument should be the demes file and the second is
    optionally the seed.")
```

#### S6.2 Continental demes

Demes file:

```
description: Based on the Gutenkunst et al. (2009) OOA model.
doi:
- https://doi.org/10.1371/journal.pgen.1000695
time_units: years
generation_time: 25
```

```
demes:
- name: ancestral
  description: Equilibrium/root population
  epochs:
  - {end_time: 220e3, start_size: 7300}
- name: AMH
  description: Anatomically modern humans
  ancestors: [ancestral]
  epochs:
  - {end_time: 140e3, start_size: 12300}
- name: OOA
  description: Bottleneck out-of-Africa population
  ancestors: [AMH]
  epochs:
  - {end_time: 21.2e3, start_size: 2100}
- name: YRI
  description: Yoruba in Ibadan, Nigeria
  ancestors: [AMH]
  epochs:
  - start_size: 12300
- name: YRIS
  description: Yoruba in Ibadan, Nigeria - Split
  ancestors: [YRI]
  start_time: 75
  epochs:
  - {start_size: 12300}
- name: CEU
  description: Utah Residents (CEPH) with Northern and Western European Ancestry
  ancestors: [OOA]
  epochs:
  - {start_size: 1000, end_size: 29725}
- name: CEUS
  description: Utah Residents (CEPH) with Northern and Western European Ancestry - Split
  ancestors: [CEU]
  start_time: 75
  epochs:
  - {start_size: 29725}
- name: CHB
  description: Han Chinese in Beijing, China
  ancestors: [OOA]
  epochs:
  - {start_size: 510, end_size: 54090}
- name: CHBS
  description: Han Chinese in Beijing, China - Split
  ancestors: [CHB]
  start_time: 75
  epochs:
  - {start_size: 54090}
- name: KAR
```

```
  description: Karitiana native Brazilians
  ancestors: [CHB]
  start_time: 15000
  #https://doi.org/10.1038/ncomms13175
  epochs:
  - {start_size: 13975, end_size: 6006}
- name: KARS
  description: Karitiana native Brazilians - Split
  ancestors: [KAR]
  start_time: 75
  epochs:
  - {start_size: 6006}

migrations:
- {demes: [YRI, OOA], rate: 25e-5}
- {demes: [YRI, CEU], rate: 3e-5}
- {demes: [YRI, CHB], rate: 1.9e-5}
- {demes: [CEU, CHB], rate: 9.6e-5}
```

Python code:

```
    import os
import msprime
import demes
import sys

if len(sys.argv) != 3:
    print("Usage: python simulate.py <demes_file_path> <output_file_path>")
    sys.exit(1)

demes_file_path = sys.argv[1]
if not os.path.exists(demes_file_path):
    print(f"File {demes_file_path} not found.")
    sys.exit(1)
output_file_path = sys.argv[2]

mutation_rate = 1e-8 #set mutation rate
sq_len = 248387328 #length of chromosome
n_individuals = 1000*2
rate_recomb = 1e-8 #average recombination rate in humans
static_seed = 5555 #seed 5555 to eliminate randomness in simulation

graph = demes.load(demes_file_path)
demography = msprime.Demography.from_demes(graph)
ts = msprime.sim_ancestry({"CEU": n_individuals,
                           "YRI": n_individuals,
                           "CHB": n_individuals,
                           "KAR": n_individuals,
                           "CEUS": n_individuals,
                           "YRIS": n_individuals,
                           "CHBS": n_individuals,
```

```
                                    "KARS": n_individuals},
                                    sequence_length=sq_len,
                                    demography=demography,
                                    random_seed=static_seed,
                                    recombination_rate=rate_recomb)
mts = msprime.sim_mutations(ts, rate = mutation_rate, random_seed = static_seed)

mts.write_vcf(sys.stdout)
```

### References

[1] F. Racimo, G. Renaud, and M. Slatkin. "Joint Estimation of Contamination, Error and Demography for Nuclear DNA from Ancient Humans". In: *PLOS Genetics* 12.4 (2016), pp. 1–27. DOI: 10.1371/journal.pgen.1005972.

[2] C. P. Robert and G. Casella. *Monte Carlo Statistical Methods*. Springer New York, NY, 2004. DOI: 10.1007/978-1-4757-4145-2.

[3] J. S. Rosenthal. "Optimal Proposal Distributions and Adaptive MCMC". In: *Handbook of Markov Chain Monte Carlo*. Ed. by S. Brooks et al. Chapman and Hall/CRC, 2011. Chap. 4, pp. 93–111. DOI: 10.1201/b10905.

[4] J. G. Schraiber. "Assessing the Relationship of Ancient and Modern Populations". In: *Genetics* 208.1 (2018), pp. 383–398. ISSN: 1943-2631. DOI: 10.1534/genetics.117.300448.

[5] A. W. van der Vaart. *Asymptotic Statistics*. Cambridge Series in Statistical and Probabilistic Mathematics. Cambridge University Press, 1998.
